# Mycobacterial Phenolic Glycolipid Triggers ATP-Mediated Neuronal P2X3 Signaling and Cough

**DOI:** 10.1101/2025.05.01.651726

**Authors:** Kubra F. Naqvi, Lois L. Warden, J. Hessel M. van Dijk, Dhananjay K. Naik, Felipe de Jesus Espinosa-Becerra, Ifunanya M. Okolie, Vincent V. F. Verhoeks, Kathryn C. Rahlwes, Christopher P. Xavier, Jeroen D. C. Codee, Theodore J. Price, Michael U. Shiloh

## Abstract

Cough drives respiratory pathogen transmission, yet how microbes directly engage host sensory neurons to trigger cough is largely unknown. We previously demonstrated that the *Mycobacterium tuberculosis* (Mtb) glycolipid sulfolipid-1 (SL-1) activates neurons and induces cough. Here, we reveal that phenolic glycolipid (PGL) produced by the hypertransmissible HN878 Mtb strain activates both mouse and human nociceptive neurons in vitro using calcium imaging and electrophysiology and is sufficient to induce cough using plethysmography. Combined with SL-1, PGL potently triggers neuronal activation. By synthesizing various PGL analogs, we show that neuroactivity is proportional to saccharide chain length and structure. Mechanistically, PGL stimulates rapid extracellular ATP release, which engages neuronal P2X3 purinergic receptors—an effect blocked by a P2X3 antagonist. These findings uncover a neuronal activation pathway co-opted by certain Mtb strains to enhance transmission via cough and suggest inhibition of purinergic signaling as a potential strategy to block airborne spread of Mtb.

## INTRODUCTION

The respiratory epithelium is innervated by sensory nociceptive neurons that respond to chemical, inflammatory, and mechanical irritants and protect the airways by initiating the cough reflex^1,2^. While cough has evolved among mammalian species as a protective reflex, it is also a hallmark sign of respiratory infection and a driver of aerosol transmission^3–5^. Despite the role of cough as a pulmonary response to respiratory infection, the molecular mechanisms by which microbially encoded compounds lead to cough remain largely unknown.

Pulmonary tuberculosis (TB), caused by *Mycobacterium tuberculosis* (Mtb) infection, is a highly inflammatory disease with persistent, bloody cough as a primary sign and route of infectious transmission^6–8^. We previously identified the Mtb cell wall glycolipid, sulfolipid-1 (SL-1), as a nociceptive neuron activating molecule that provokes coughing in naïve and Mtb-infected guinea pigs^9^. By testing a variety of Mtb isolates, we also observed differential neuronal activation due to genetic variation in SL-1 production. Beyond SL-1 production, circulating Mtb lineages also differ in synthesis of other cell wall glycolipids and some lineages are reported to have enhanced virulence and transmissibility^10^. In particular, the lineage 2/Beijing strains of Mtb have a higher propensity for bacterial growth, drug resistance, and frequency of transmission^10–12^. HN878, a representative lineage 2 strain of Mtb, is hypervirulent during mouse infection due to its ability to induce anti-inflammatory innate immune signaling characteristic of Th2 immunity^13,14^. This phenotypic characteristic is genetically linked to a polyketide synthase gene, *pks1-15,* and production of the *pks1-15* encoded lipid, phenolic glycolipid (PGL)^13^. In contrast, community and laboratory Mtb lineage 4 strains such as Mtb Erdman, H37Rv, and CDC1551 do not produce PGL due to a naturally occurring frameshift mutation in *pks1-15*^14,15^. Given the propensity of lineage 2 strains for enhanced virulence and transmission, and known changes to their cell wall lipid repertoire^13,16–18^, we hypothesized that these strains may encode additional nociceptive neuron activating molecules to trigger the cough reflex.

Here we identify PGL as a nociceptive neuron agonist produced by lineage 2 Mtb strains. Exposing nociceptive neurons to purified PGL elicited increased intracellular calcium and both nociceptive molecules, SL-1 and PGL, cooperatively activated neurons. PGL triggered activation of primary mouse dorsal root ganglia (DRG) and nodose ganglia neurons and primary human DRG neurons, and depolarized primary human nociceptors. Through structure-activity relationship studies of PGL analogs, obtained through organic synthesis, we identified that the saccharide chain of PGL was sufficient to trigger neuronal activation while the full lipid chain was dispensable for activity. Additionally, exposure of naïve guinea pigs to purified PGL was sufficient to induce the cough reflex. Finally, we identified that PGL activated neuronal signaling by stimulating release of extracellular ATP (eATP) through pannexin channels, and this signaling pathway was abrogated by chemical inhibition of cell surface purinergic receptors. Taken together, we identify a mechanism by which Mtb PGL triggers the cough reflex, with implications for understanding and preventing transmissibility of circulating Mtb lineages.

## RESULTS

### Mtb strains differ in neuron activation due to variable production of phenolic glycolipid

To determine if Mtb strains considered to be hyper-transmissible^19^ produce additional nociceptive compounds beyond SL-1, we extracted lipids from two distinct strains of Mtb, Erdman and HN878. We assessed neuronal activation by these lipid extracts in vitro using live-cell calcium imaging of the nociceptive neuronal cell-line, MED17.11. Although both strains produce SL-1 (Supplemental Figure 1A, related to Figure 1), we found that an organic extract from the lineage 2 strain HN878 elicited higher neuronal activation compared to the lineage 4 Erdman strain (Figure 1A). A major genetic distinction between these two strains lies in the polyketide synthase gene, *pks1-15,* which is naturally mutated in Mtb Erdman but in frame in HN878^14,15^. The polyketide synthase pks15/1 is part of the first committed step in the biosynthesis of the mycobacterial cell wall lipid PGL^20,21^. Beginning with the precursor molecule *p-*hydroxy benzoic acid, a series of enzymes including other polyketide synthases, ligases, and methyltransferases extend the lipid core to form *p-*hydroxyphenol phthiocerol dimycocerosate (PDIM)^22^. Finally, glycosyl and methyltransferases attach and elongate the glycan chain of PGL^23^. The resultant structure includes a lipid chain (PDIM) linked to an aromatic ring (phenolic group), and an oligosaccharide (Figure 1B).

**Figure 1:**
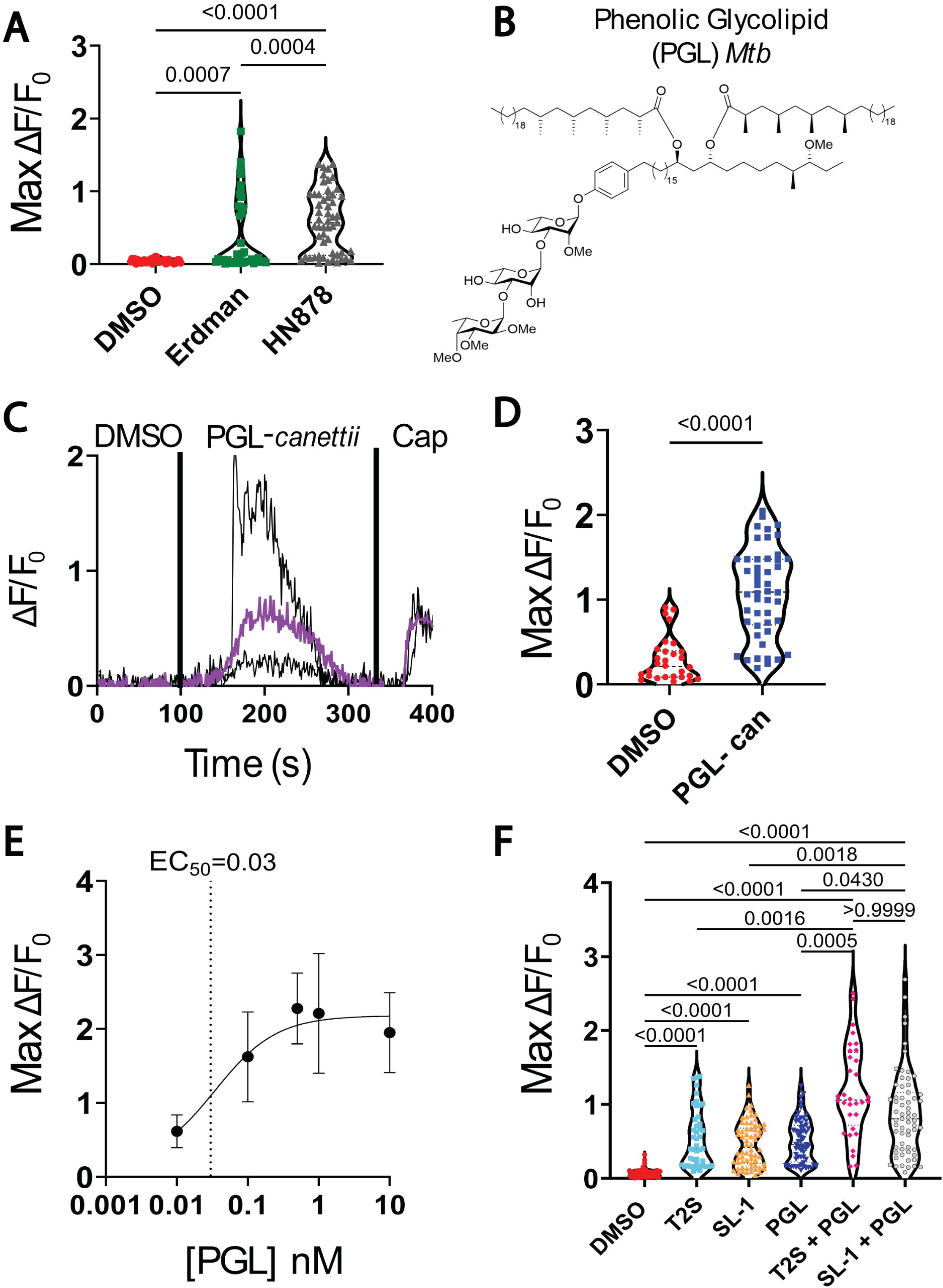
Phenolic Glycolipid (PGL) Activates Nociceptive neurons. (A) Intracellular Ca^2+^ levels of MED17.11 nociceptive neurons treated with lipid extract from Mtb strains Erdman and HN878. (B) Mtb PGL structure containing trisaccharide chain. (C) Intracellular Ca^2+^ of MED17.11 neurons activated with purified *Mycobacterium canettii* 1nM PGL and capsaicin (cap), average response of 57 cells in purple. (D) Maximum calcium change in MED17.11 stimulated with 1nM PGL *canettii* compared to vehicle DMSO. (E) EC_50_ of purified PGL *M. canettii* for activation of nociceptive neurons. (F) Activation of nociceptive neurons by trehalose 2-sulfate (T2S, 100nM), sulfolipid-1 (SL-1, 100nM), PGL (100 nM), and 1:1 equimolar concentration of PGL and T2S or SL-1. *P-* values calculated by Kruskal-Wallis. EC_50_ calculated by nonlinear regression analysis.

Because PGL is produced by Mtb HN878 but not Mtb Erdman, we hypothesized the PGL could serve as an additional nociceptive molecule when comparing both extracts. To test this hypothesis, we treated MED17.11 neurons in vitro with PGL purified from *M. canetti.* When the cells were treated with PGL alone, intracellular calcium increased significantly compared to the response to the vehicle DMSO and was comparable to the levels elicited by the positive control capsaicin, a TRPV1 agonist^24^ (Figure 1C). Capsaicin-responsive nociceptive neurons exhibited higher maximum calcium levels when treated with PGL alone compared to those that were only treated with the vehicle DMSO (Figure 1D). Having established the nociceptive phenotype of PGL, we next determined the potency of PGL in a dose response assay of neuronal activation as assessed by increased levels of intracellular calcium and quantified an EC_50_ of 30 pM (Figure 1E). To ensure that the activity detected from the natural product *M. canetti* PGL was due to PGL alone and not an alternative contaminating molecule, we generated PGL-Mtb through organic synthesis. Three different synthetic Mtb-PGLs, differing only in the location of the methyl ethers on the trisaccharide structure, recapitulated the activation phenotype of nociceptive neurons, further confirming PGL as the active compound (Supplemental Figure 2A-B, related to Figure 1). Subsequent experiments were performed with the synthetic PGL-1 (Supplemental Figure 2A, related to Figure 1), henceforth referred to as PGL.

While both Mtb Erdman^25^ and HN878^9^ produce the established nociceptive glycolipid SL-1, only Mtb HN878 produces both SL-1 and PGL^13^. To test how the combination of SL-1 and PGL triggers neuron activation, we treated MED17.11 cells with equimolar concentrations of both compounds. As previously established, trehalose-2-sulfate (T2S) is the minimal active component of SL-1^9^ and significantly activated neurons in vitro (Figure 1F). Similarly, full length SL-1 and PGL alone triggered neuron activation (Figure 1F). When neurons were exposed to both compounds simultaneously at an equimolar ratio (100 nM each), mimicking production of both molecules by Mtb HN878, we found significantly higher neuron activation compared to treatment of neurons with each compound alone and the vehicle control DMSO (Figure 1F). We observed this increased response when combining PGL with either the minimal compound T2S or SL-1. Taken together these results identify PGL, produced by the HN878 strain, as a nociceptive neuron activating molecule in vitro and establish that the combination of Mtb SL-1 and PGL triggers significantly increased intracellular calcium compared to either molecule alone.

### PGL directly activates lung-innervating nociceptive neurons

Having established that PGL is a nociceptive molecule produced by Mtb, we next determined if PGL could interact with lung-innervating nociceptive neurons. The airways are innervated by neurons from distinct sensory afferent subsets. The larger airways are innervated by neurons stemming from the dorsal root ganglia (DRG) while vagal neurons from the nodose and jugular ganglia innervate conducting airways and project to the alveolar region^26^. We cultured primary mouse nociceptive neurons from both the DRG and nodose ganglia and tested for neuronal activation in response to PGL by live cell calcium imaging. Synthetic PGL alone activated not only the nociceptive cell line MED17.11 (Figure 2A), but also primary mouse DRG and nodose neurons (Figure 2D and G) as demonstrated by large spikes in intracellular calcium. Among the cells stimulated with PGL, a large percentage (>40%) were responsive to both PGL and capsaicin for all cell types tested. Additionally, most of the cells identified as capsaicin-responsive nociceptors were also responsive to PGL, ranging from 60-90% (Figure 2B, E, and H). Compared to cells treated only with the vehicle (DMSO), those treated with PGL alone resulted in significantly higher maximum calcium elevation (Figure 2C, F, and I). Importantly, primary DRG and nodose neurons exhibited pronounced responsiveness to PGL, with the Max ΔF/F_0_ reaching between 1-3. These data further establish PGL as a nociceptive molecule and its ability to activate lung-innervating neurons that orchestrate the cough reflex.

**Figure 2:**
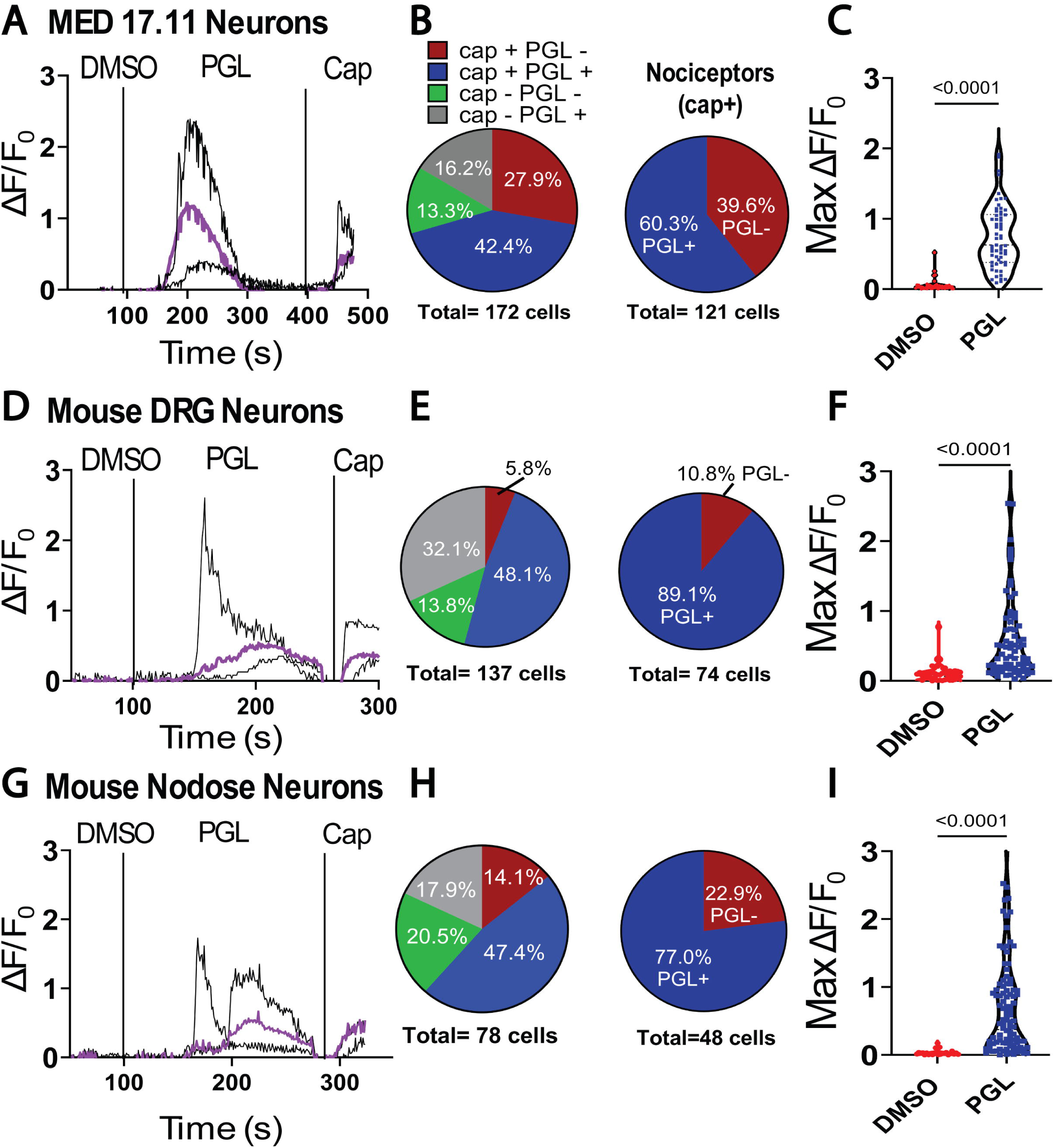
Mtb PGL Activates Primary Lung Innervating Nociceptive Neurons. (A) Intracellular Ca2+ levels of MED17.11 neurons activated with organically synthesized Mtb PGL and capsaicin (cap) over time (seconds). Average response of 64 cells in purple. (B) Pie chart depicting percentage of MED17.11 cells responsive to cap, PGL, or both compounds. Percent of nociceptors (cap+) responsive to PGL depicted as a pie chart. (C) Maximum fluorescent calcium signal following treatment with vehicle (DMSO) or PGL (1nM) of MED cells. (D) Intracellular Ca2+ change of primary murine dorsal root ganglion (DRG) neurons activated with PGL and cap with average response of 23 cells in purple. (E) Pie chart of DRG neurons responses to cap and/or PGL and percentage of nociceptors responsive to PGL. (F) Maximum fluorescent calcium signal to vehicle (DMSO) or PGL of DRG neurons. (G) Intracellular Ca2+ levels of primary nodose ganglion neurons activated with PGL and cap. Average response of 20 cells in purple. (H) Percentage of nodose neurons responsive to cap and/or PGL and associated nociceptor responsiveness to PGL. (I) Maximum fluorescent calcium signal to vehicle (DMSO) or PGL of nodose neurons. *P-* values calculated by Mann-Whitney

### Human neurons are activated by PGL

Due to the human tropism of Mtb infection and to further establish the nociceptive function of PGL, we assessed PGL responses in primary human DRG neurons. We treated DRG derived neurons from two human donors with PGL and observed a significant increase in intracellular calcium compared to the vehicle DMSO (Figure 3A and D). Like mouse neurons, most of the human cells (>50%) were both PGL and capsaicin responsive (Figure 3B and E) and the maximum change in fluorescence due to PGL treatment was significantly higher than for DMSO (Figure 3C and F).

**Figure 3:**
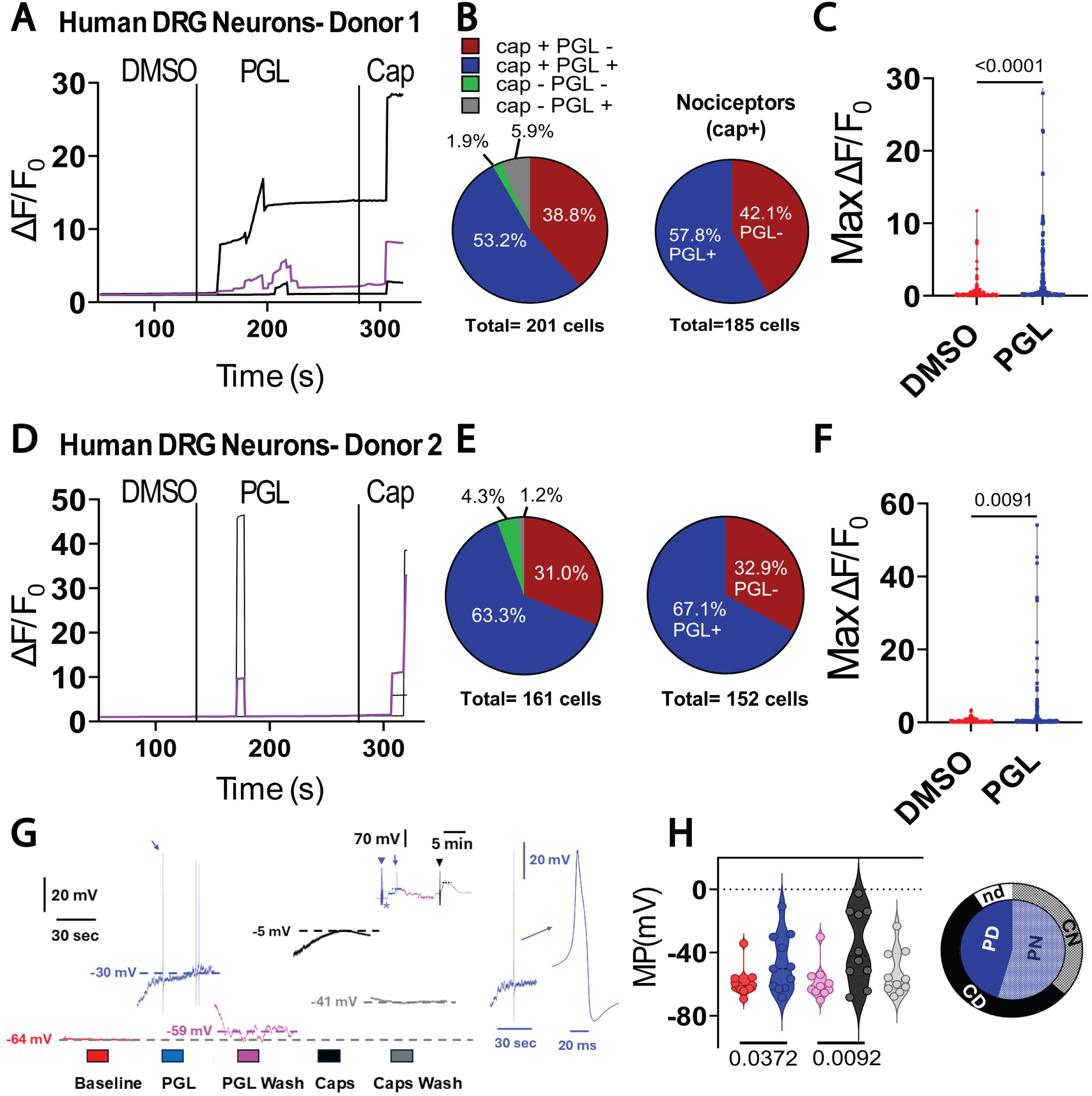
PGL Exposure Activates and Depolarizes Human DRG Neurons. (A) Human donor 1 DRG neuron calcium levels upon exposure to vehicle (DMSO), PGL (1nM), and capsaicin (cap), average of 17 cells response in purple. (B) Pie chart depicting percentage of cells responsive to cap, PGL, or both compounds and response of nociceptors (cap+) to PGL from donor 1. (C) Maximum fluorescent calcium signal following treatment with DMSO or PGL. (D) Calcium change in human donor 2 DRG neurons upon exposure to DMSO, PGL, and cap. Average of 29 cells in purple. (E) Pie chart depicting percentage of cells responsive to cap, PGL, or both compounds and response of nociceptors (cap+) to PGL from donor 2. (F) Maximum calcium fluorescence following treatment with DMSO or PGL. (G) Non continuous 1-min membrane potential traces before/after PGL and before/after capsaicin (caps) exposure. Blue inset shows details of the first spontaneous action potential. Black inset details a 22 min trace showing time flags of the addition of PGL (blue arrowhead), and capsaicin (black arrowhead). (H) Violin plot of the effects of PGL and caps on the membrane potential. Pie chart depicting cells that depolarized with PGL (PD) and to caps (CD). The non-responsive groups are labeled PN (for PGL) and CN (for caps). Cell not tested with caps (nd). *P-* values calculated by Mann-Whitney

To further analyze the mechanisms of PGL excitation of human DRG nociceptors, we conducted patch-clamp experiments of individual neurons. We tested the impact of PGL at rest and during evoked firing of eleven human nociceptive neurons. To test the impact of PGL on membrane potential (MP), some cells were recorded at their resting membrane potential (RMP, no holding current applied; n=5), and other cells were held at −60mV to test the impact of PGL on firing properties (n=6). Dissociated human nociceptors are mostly silent (no spontaneous firing) during rest or between stimulations^27,28^, therefore we tested if perfusion of PGL would induce action potential (AP) firing. While PGL did not induce regular AP firing, two cells did fire one or more APs during perfusion (Figure 3G, inset asterisk). Additionally, five of the eleven cells depolarized five or more mV in the presence of PGL. Figure 3G shows a neuron recorded at the resting membrane potential and where PGL reversibly depolarized 34 mV and fired three APs (blue trace). On average, PGL at 1 nM induced a depolarization of 2.1 mV while 100-300nM capsaicin depolarized 5 mV or more in seven of the ten cells (Figure 3H). We note that the EC_50_ for PGL-induced intracellular calcium elevation was determined to be 30 pM (Figure 1E), while it is between 100-500 nM for capsciacin^24^. The pie chart (Figure 3H) demonstrates that, except for one cell not tested with capsaicin (nd), the remaining cells that depolarized with PGL (PD) were also capsaicin-responsive (CD). These data not only validate our findings in mouse primary neurons but also extend to primary human neurons the activity of PGL as a neuronal agonist triggering increased intracellular calcium and neuron depolarization.

### The saccharide chain structure of PGLs impacts nociceptive activity

PGL is produced not only by Mtb but also by other slow growing mycobacteria including *M. bovis, M. leprae,* and *M. kansasii*, with the glycan moiety distinguishing PGLs produced by each organism. PGL produced by *M. bovis*, mycoside B, shares the common lipid chain (PDIM) with Mtb but contains only the monosaccharide rhamnose^29^.

*M. kansasii* PGL includes a more decorated tetrasaccharide chain, with a 2,6-dideoxysugar as the terminal residue ^30,31^. Both PGLs from Mtb and *M. leprae* contain a trisaccharide chain; however, the terminal sugars are fucose and glucose, respectively^32,33^. Additionally, the carbohydrate chain structure differs between Mtb and *M. leprae* PGLs: where PGL-leprae contains a unique β1-4 linkage of the terminal glucose compared to the α1-3 linkage of the terminal fucose of PGL-Mtb^34^. To determine if structural differences amongst the PGLs produced by pathogenic mycobacteria differ in their ability to activate neurons, we tested various synthetic PGLs reflecting each species described above^35,36^. We treated MED17.11 cells with synthetic PGLs from *M. bovis, M. leprae,* Mtb, and *M. kansasii* (Figure 4A) at the same concentration (1 nM) and measured neuronal activation by live-cell calcium imaging. While PGL from *M. bovis* and *M. leprae* were able to induce modest elevation of intracellular calcium, we observed significantly higher calcium levels in neurons exposed to PGLs from Mtb and *M. kansasii* (Figure 4B). This phenotype was consistent when we tested the same PGL compounds on primary mouse DRG neurons (Figure 4C). We further characterized the nociceptive activities of mycobacterial PGLs by performing dose response studies for each PGL molecule and calculating their EC_50_. The EC_50_ for neuron activation by monosaccharide (*M. bovis)* and disaccharide forms of PGL were significantly higher (nM to µM range) than the EC_50_ for the trisaccharide (Mtb) and tetrasaccharide PGLs (pM to nM range) (Supplemental Table 1, related to Figure 4). Additionally, PGL from *Mycobacterium haemophilum*, which also contains a trisaccharide form but with rhamnose as the terminal sugar, compared to fucose for Mtb PGL, significantly activated neuronal calcium elevation (Supplemental Figure 3A-B, related to Figure 4). These data demonstrate that the length and type of sugar linked to the phenol and PDIM lipid are vital to the nociceptive activity of mycobacterial PGLs.

**Figure 4:**
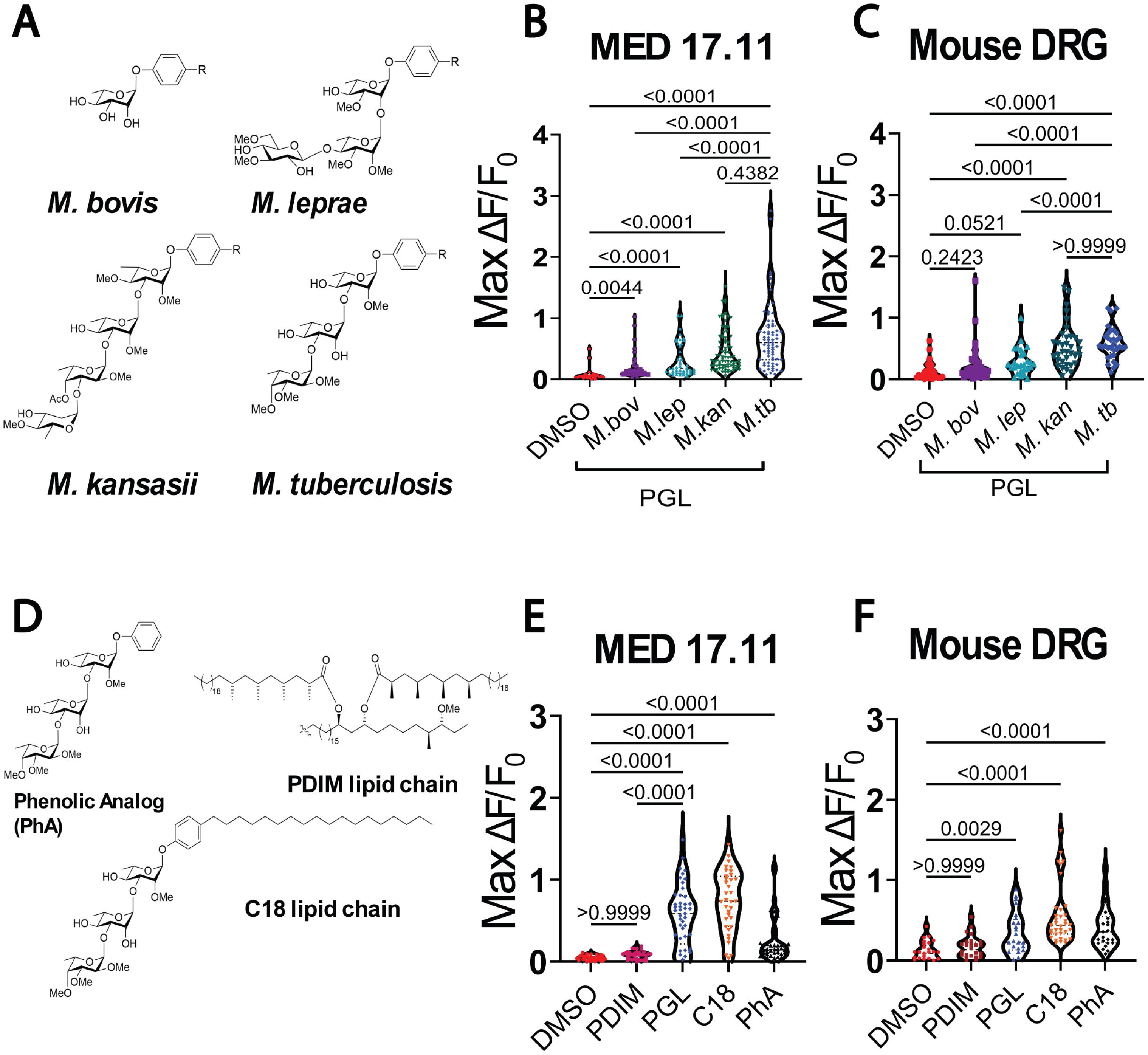
PGL Saccharide Chain Influences Structure Activity Relationship. (A) Saccharide chain structure of PGLs from mycobacterial species *bovis, leprae, kansasii,* and *tuberculosis.* R= PDIM lipid chain. (B) Activation of MED17.11 nociceptive neurons and (C) DRG neurons by mycobacterial PGLs at 1nM concentration. (D) Structure of PGL analogs and PDIM lipid chain. (E) Activation of MED17.11 cells and (F) DRG neurons by PGL analogs at 1nM concentration. *P*-values calculated by Kruskal-Wallis

To further evaluate the structure activity relationship of PGL, we next tested PGL analogs, modified with respect to the lipid chain. We synthesized PGL analogs containing the full lipid chain (Figure 1B), a simplified C18 lipid chain, or no lipid chain (phenolic analog, PhA) and compared these to the common full lipid chain, PDIM, alone (Figure 4D). We tested each analog for neuron activation on MED17.11 cells at the same concentration (1 nM). We observed that only Mtb PGLs containing the sugar chain (PGL-Mtb, C18, and PhA) were able to activate neurons, while PDIM alone resulted in no significant activation phenotype (Figure 4E). This phenotype was also conserved when the same compounds were tested using primary mouse DRG neurons (Figure 4F). Furthermore, the phenolic analogs lacking the PDIM moiety of the species specific PGLs exhibited the same phenotype as their full-length forms (Supplemental Figure 3 C-D, related to Figure 4) further highlighting the saccharide chain as the functional component of PGL. Taken together, these data demonstrate that while multiple forms of mycobacterial PGLs activate nociceptive neurons, the activity of PGL is highly dependent on the presence, length, and structure of the attached glycan chain while the full-length lipid core is largely dispensable for the observed phenotype.

### Purified PGL triggers a cough response in guinea pigs

Given the in vitro nociceptive function of PGL, we hypothesized that PGL could serve as a cough inducing molecule produced by Mtb. To test this, we employed an established methodology for measuring cough in naïve guinea pigs using whole body plethysmography^9,37–40^. Healthy unanesthetized guinea pigs were placed inside the plethysmography chamber followed by continuous pressure recording over 10 minutes during compound nebulization and 10 minutes of acclimation, 20 minutes total (Figure 5A).

**Figure 5:**
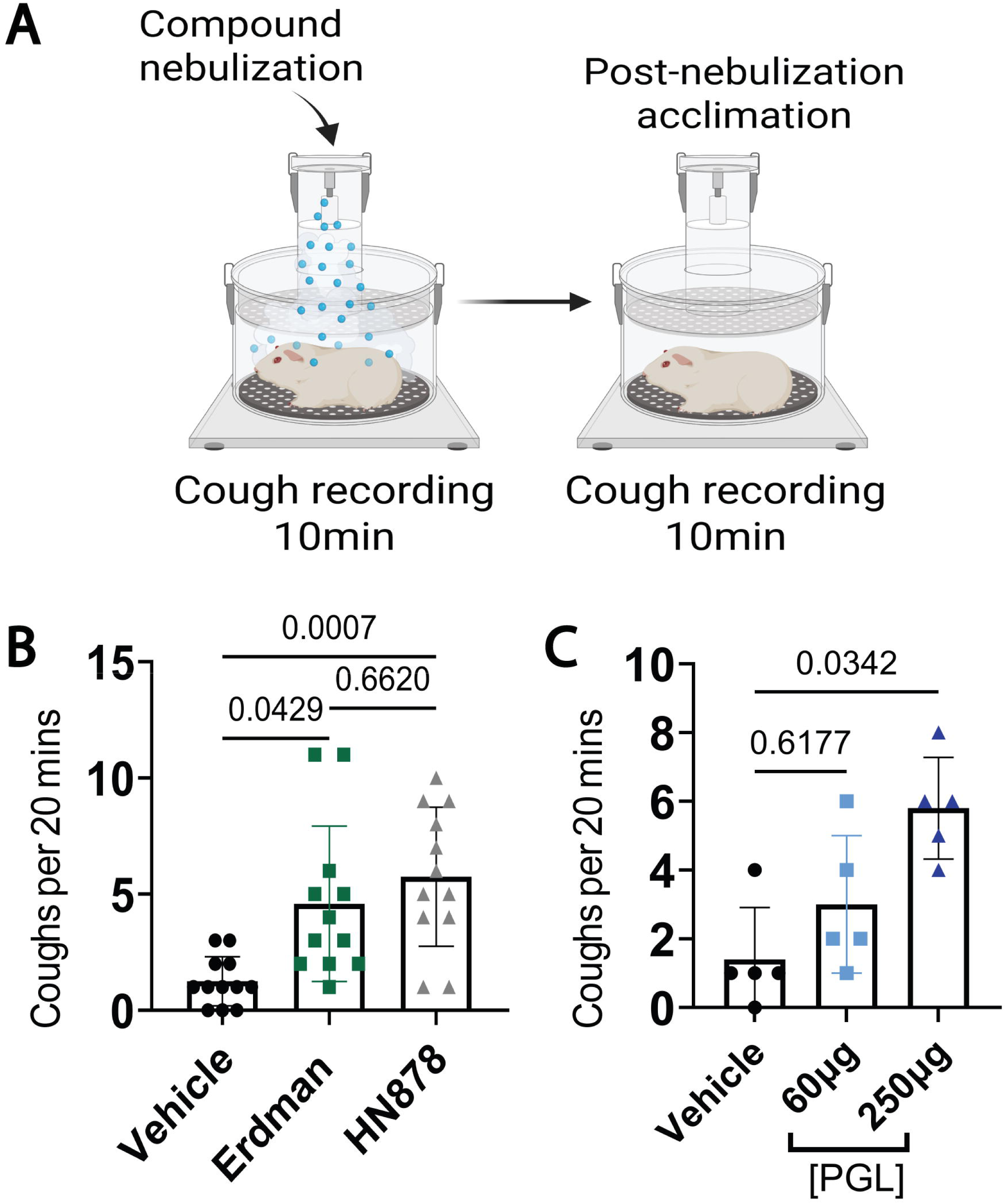
PGL Triggers Cough in Healthy Guinea Pigs. (A) Whole body plethysmography-based cough recording of naïve unanesthetized guinea pigs. (B) Coughs per 20 mins recorded from guinea pigs nebulized with vehicle (10% methanol: PBS) or lipid extract from Erdman and HN878 strains of *Mtb.* (C) Coughs induced by nebulization of organically synthesized PGL *Mtb.* P-value calculated by Friedman test

To determine if PGL triggers the cough reflex in guinea pigs, we first nebulized lipid extract from the Mtb Erdman and HN878 strains. Guinea pigs were first exposed to the vehicle (10% methanol: PBS) then the same animals were treated with 20 mg/mL of lipid extract with 2-3 days of rest between cough recordings. Prior to nebulization, the lipid extract was confirmed for the presence of SL-1 and PGL by mass spectrometry and TLC, respectively (Supplemental Figure 1 A-B), and guinea pigs were treated with a positive control cough agonist, citric acid, at the end of study. While we did observe significantly more coughs in the Mtb lipid extract treated groups compared to the vehicle, there was no statistically significant difference between the Mtb Erdman and HN878 extracts (Figure 5B). This could be due to the bioavailability of nociceptive compounds, SL-1 and PGL, during the nebulization period or variation in the amount of SL-1 and PGL in the lipid extracts. We next tested if PGL alone could trigger cough in healthy guinea pigs. A separate cohort of guinea pigs was placed inside the plethysmography chamber and exposed to the vehicle control (10% methanol: PBS) or two different doses of PGL (60 and 250 µg/mL). When treated with the lower dose of PGL, guinea pigs exhibited an elevated cough response; however, this did not reach statistical significance. The higher concentration (250 µg/mL), however, triggered significantly more coughs in the same animals compared to the vehicle control (Figure 5C). All guinea pigs included in the cough studies exhibited significant cough responses due to citric acid treatment, confirming their neurophysiological ability to cough (Supplemental Figure 4 A-B, related to Figure 5). Based on these results, PGL not only activates nociceptive neurons in vitro but is also sufficient to trigger the cough reflex.

### Extracellular ATP serves as a secondary messenger for neuronal activation by PGL

Having established the nociceptive function of PGL, we next sought to determine the mechanistic basis for PGL-dependent neuronal activation. While PGL is a potent activator of nociceptors, we observed a 50-to 100-second delay in activation across the neuronal cells tested, which contrasted with the rapid increase observed by capsaicin mediated opening of TRPV1 channels^24^ (Figure 2). This delayed response suggests that exposure of neurons to PGL may trigger release of a secondary messenger to activate neurons. Indeed, prior work has demonstrated the importance of secondary messengers in mediating nociceptive functions like pain and cough^41,42^. Among the known secondary messengers released by neurons, extracellular ATP (eATP) has been implicated as an alarmin that mediates inflammatory and neuropathic pain^43^ as well as chronic cough^44,45^. Upon stimulation by exogenous signals^46^, sensory neurons and surrounding accessory cells can release eATP through pannexin channels^47,48^ and further activate purinergic receptors^49^ to activate neuronal signaling (Figure 6A). Thus, we hypothesized that PGL may leverage this signaling pathway through release of the secondary messenger eATP. To test this hypothesis, we first stimulated MED17.11 cells with PGL and measured extracellular ATP released into the supernatant after exposure. Cells stimulated with PGL (100 nM), or the positive control potassium chloride (50 mM), released an average of 1.5-fold more ATP compared to the vehicle control DMSO (Figure 6B) suggesting that ATP may be involved in the broader neuronal response to PGL.

**Figure 6:**
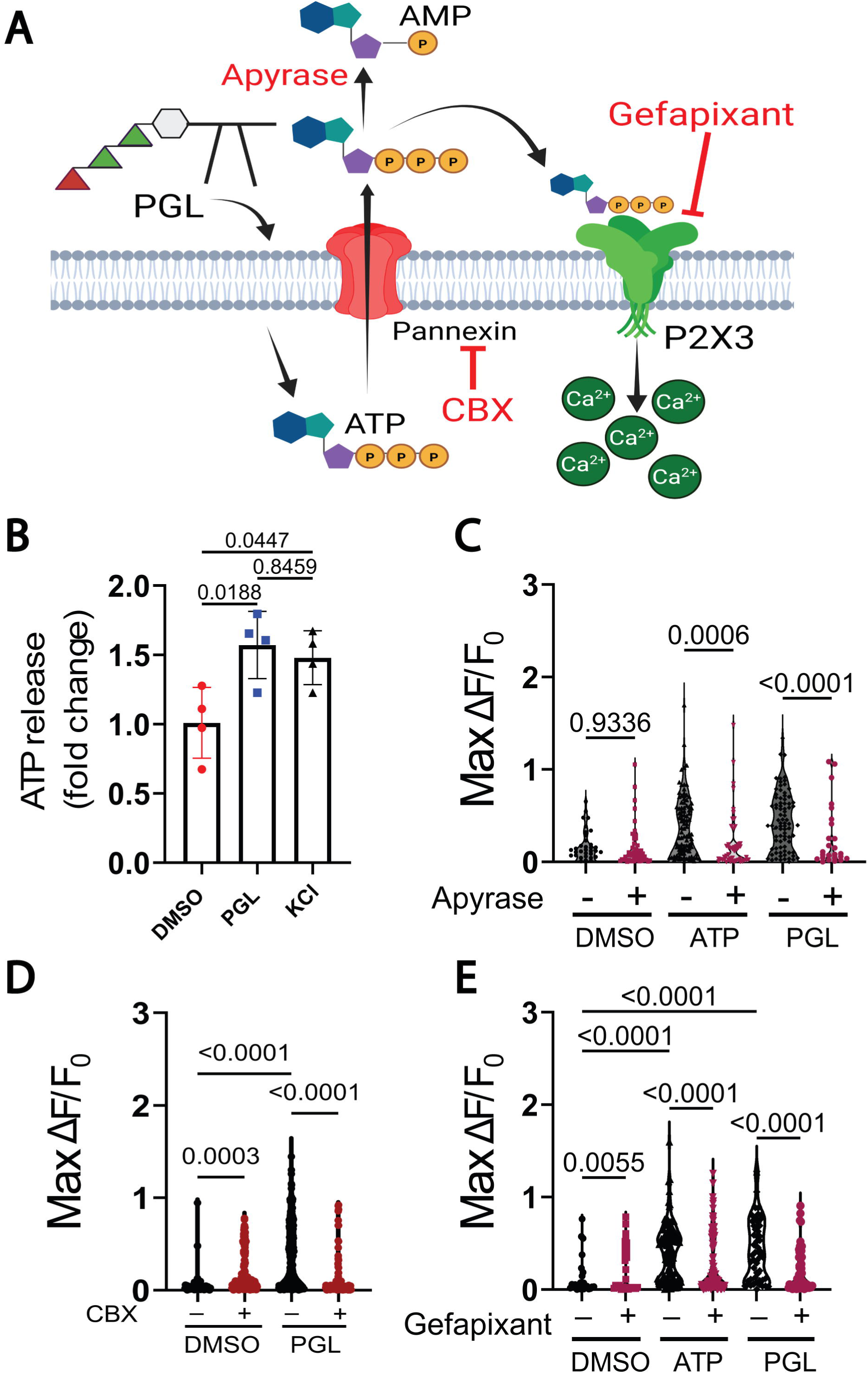
PGL Activates Neurons Through the Secondary Messenger ATP. (A) Signaling pathway for ATP release and purinergic receptor antagonism by chemical inhibitors. (B) fold change of extracellular ATP release by nociceptive neurons stimulated with DMSO vehicle, PGL (100nM) or KCl (50mM) (C) Intracellular calcium levels of MED17.11 cells with or without apyrase (5U/mL) treatment prior to stimulation with positive control ATP (2µM) or PGL (100nM). (D) Calcium levels from MED17.11 cells treated with the pannexin channel inhibitor carbenoxolone (CBX, 10 µM) and (E) P2X3 inhibitor, gefapixant (10µM), or vehicle control (DMSO) prior to stimulation with ATP or PGL. *P*-values calculated by Kruskal-Wallis.

To further assess the role of eATP in PGL dependent neuron activation, we stimulated MED17.11 neurons with PGL in the presence of the enzyme apyrase, which catalyzes the hydrolysis of eATP (Figure 6A). As a positive control for apyrase activity, and to test the ability of eATP to activate MED17.11 cells, we first compared eATP or eATP plus apyrase in live cell calcium responses. Exposure of MED17.11 neurons to eATP led to a significant increase in intracellular calcium, which was inhibited by concomitant treatment with apyrase (Figure 6C). Likewise, when we co-incubated neurons with PGL and apyrase, PGL-dependent neuron activation was markedly reduced (Figure 6C).

ATP can be released through pannexin channels^50–53^, and such channels are involved in a variety of biological processes, including neuropathic pain^42^. Furthermore, the *PANX1* gene, encoding pannexin 1 protein, is highly expressed in human and mouse nociceptors in the DRG^54^ and the protein is detected in hDRG using unbiased proteomic methods^55^. To determine if pannexin channels are involved in the PGL-dependent release of eATP, we pretreated MED17.11 cells with the pannexin1 inhibitor carbenoxolone (CBX)^56^. We observed a modest increase in intracellular calcium in cells treated with both vehicle (DMSO) and CBX compared to vehicle alone; however, the average responses (CBX+DMSO average 0.17 and DMSO alone average 0.07) were markedly lower than that observed for PGL (Figure 6D). CBX treatment prior to exposure to PGL, significantly reduced neuronal activation compared to cells without CBX treatment (Figure 6D). These results demonstrate that pannexin 1 channels contribute to accumulation of eATP following PGL exposure.

Once released into the extracellular environment, eATP can signal through purinergic receptors (P2X and P2Y) to activate nociceptive pathways in sensory neurons of the DRG and nodose ganglia^5,57,58^. In addition, P2X3 receptor activity has been implicated in chronic cough^59,60^, and two P2X3 receptor antagonists have reached phase 3 human studies for efficacy in refractory chronic cough^61,62^. Thus, we tested the impact of blocking P2X3 receptors on PGL-dependent neuron activation. MED17.11 cells were pretreated with the P2X3 inhibitor gefapixant^63,64^ or vehicle control prior to calcium imaging. As expected, eATP-induced MED17.11 neuron activation was inhibited by gefapixant (Figure 6E). Likewise, gefapixant pretreatment significantly inhibited PGL-mediated neuronal activation (Figure 6E). Taken together these data suggest that eATP serves as a secondary messenger for the nociceptive function of PGL and provides a mechanistic target for therapeutic intervention.

## DISCUSSION

In this study, we identify PGL, produced by a hypervirulent and hypertransmissible strain of Mtb, as a potent activator of nociceptive neurons and inducer of cough. Purified PGL triggers intracellular calcium flux in mouse dorsal root ganglia and nodose ganglia neurons, and human DRG neurons. Using synthetic PGL analogs, we show that nociceptive activity is conferred by the saccharide moiety, while the PDIM lipid is dispensable. Importantly, PGL variants with longer saccharide chains such as those from Mtb exhibited enhanced neuronal activation compared to PGLs from other non-tuberculous mycobacteria. In vivo, aerosolized PGL alone is sufficient to induce dose-dependent coughing in naïve guinea pigs. Mechanistically, PGL stimulates the release of extracellular ATP (eATP), which acts through the P2X3 receptor to activate sensory neurons.

These findings expand our understanding of Mtb-host interactions by linking a specific bacterial lipid to direct neuronal activation and cough induction - a process with major implications for airborne transmission. Although cough is widely regarded as a major driver of Mtb aerosolization, other respiratory actions such as singing^65^, speaking and breathing ^66^ can contribute to transmission. Nevertheless, the ability of the HN878 strain to produce PGL and trigger enhanced neuron activation adds to its known immunomodulatory activities, including dampening of pro-inflammatory responses^13,67^ and skewing toward Th2 immunity^13^. Notably, the lineage 2/Beijing strains of Mtb are among the most widespread globally due to enhanced frequency of transmission and poor treatment outcome^68–70^. While lipid extracts from HN878 and Erdman strains did not differ significantly in their capacity to induce cough, purified PGL alone triggered robust coughing, suggesting lipid organization and delivery dynamics may influence in vivo activity. Though aerosol delivery of cough agonists to conscious, unrestrained animals is the most physiologically relevant approach to measure cough induction^37–40^, we are unable to determine the precise amount of specific lipid species (i.e. SL-1 and PGL) inhaled by each animal during nebulization or whether complex micelles are formed, which may prevent lipids from reaching the neurons innervating distal alveoli. These limitations may confound our lipid extract cough results. Future work is needed to quantify individual lipid species during aerosolization and infection and assess their precise contributions to transmission.

PGL production varies across Mtb lineage 2 strains^13^. While many retain the intact *pks1-15* locus required for PGL biosynthesis, some subgroups lack functional PGL despite retaining the gene^18^, suggesting alternative regulatory or mutational constraints. Intriguingly, outbreaks have been associated with both PGL-positive and PGL-deficient strains^71^, implying that while PGL may enhance transmission via cough, it is not the sole determinant. Other lipids, such as SL-1, may compensate in certain genetic backgrounds, a hypothesis that merits further investigation. Additionally, Mtb strains devoid of PGL synthesize structurally related *p-*hydroxybenzoic acid derivatives (*p-* HBADs) containing the oligosaccharide moiety^21,72^. Given the activity of the phenolic analog, it is possible that release of *p-*HBADs may also contribute to cough induction in strains unable to synthesize the full PGL molecule.

Beyond Mtb, PGLs are produced by diverse pathogenic mycobacteria, each with distinct immunomodulatory roles. *M. leprae* PGL mediates bacterial entry into phagocytes ^73^ leading to macrophage activation and nerve damage^74–76^. Similar to PGL produced by Mtb, PGLs from *M. kansasii* and *M. leprae* suppress pro-inflammatory cytokines or modulate host immunity^77,78^ while PGL from *M. marinum*, structurally distinct from PGL-Mtb, induces myeloid cell recruitment and chemokine production by infected macrophages^79,80^. Across species, the saccharide moiety consistently emerges as a critical determinant of host interaction, influencing receptor engagement and immune evasion^81^. Our data extend this paradigm by linking glycan structure to neuronal activation and cough induction, particularly among respiratory pathogens like Mtb and *M. kansasii*^82^.

Bacterial pathogens including *Staphylococcus aureus, Streptococcus pyogenes, Klebsiella pneumonia* and *E. coli* directly interact with nociceptive neurons to cause pain^83,84^, itch^85^, and inflammation^86–88^. While we have yet to identify a direct PGL receptor on nociceptive neurons, the saccharide chain likely governs receptor binding affinity or specificity. Similarly, trehalose-2-sulfate, a sulfated disaccharide precursor in the SL-1 synthesis pathway lacking acyl chains, is also sufficient to activate neurons in vitro^9^, highlighting that a shared feature of nociceptive neuron activation by mycobacterial glycolipids is recognition and response to bacterial saccharides. Indirect pathways may also contribute. For instance, *Candida albicans* activates sensory neurons via keratinocyte-derived eATP in response to β-glucan^89^. Similarly, we show that PGL-driven neuron activation is mediated by eATP signaling and is blocked by eATP depletion and purinergic receptor antagonism. These insights connect microbial lipids to a broader neuroimmune signaling axis.

P2X3 receptor antagonists, such as gefapixant and camlipixant, are already in clinical development for chronic cough^61,90,91^. Our findings suggest they may have utility in treating eATP-triggered infectious cough as well. Such host-directed therapeutics, used alongside antibiotics, could reduce transmission during peak bacterial shedding both for Mtb^7,92–95^ and other pandemic pathogens^96^. Future studies should assess whether purinergic blockade can reduce cough and limit aerosolized bacterial particles during infection.

In summary, we describe a previously unrecognized function for Mtb PGL in triggering cough via nociceptive neuron activation and eATP signaling. These findings highlight a novel mechanism of pathogen-host communication that promotes Mtb transmission and opens therapeutic opportunities to disrupt cough-mediated spread.

## MATERIALS AND METHODS

### Bacterial culture conditions

Mtb Erdman (laboratory of J. Cox, UC Berkeley) and HN878 (BEI Resources, #NR-13647), were grown in Middlebrook 7H9 medium (Sigma, #M0178) supplemented with 10% oleic acid-albumin-dextrose-catalase, 0.5% glycerol and 0.05% freshly prepared tween-80. Bacteria were grown using a roller apparatus and maintained in roller bottles at 37°C until preparation for lipid extractions.

### Extraction of mycobacterial lipids

Mtb cultures were grown to an optical density_600_ of 0.8-1.0 in 7H9 medium and bacterial pellets from 400mL of culture were collected by centrifugation at 3,500 RPM (Thermo Sorvall Legend XTR) for 10 mins. Bacterial pellets were resuspended in 2mL of PBS and transferred to 45 mL 2:1 chloroform: methanol in a glass conical tube and incubated for 16 hours at 58°C. Samples were then centrifuged at 2000 RPM (Allegra X-14R) for 5 mins and the aqueous (upper) phase and cellular debris were discarded. To the remaining organic phase, 8mL of water was added and the samples were centrifuged again for phase separation^97^. After removal of the aqueous phase, the organic layer interface was washed with 2mL 1:1 methanol:water to remove any debris. The remaining lipids were concentrated at 65°C under nitrogen gas (Techne Dri-Block and Sample Concentrator).

For detection of SL-1, lipid extracts were resuspended in 1:1 MeOH: IPA 16mM NH_4_F to a final concentration of 4mg/mL. The samples were then infused on a quadrupole TOF TripleTOF 6600+ (SCIEX) mass spectrometer using previously established^9^ source parameters. PGL was detected by thin layer chromatography (TLC) of lipid extracts. Lipids were resuspended in chloroform to a final concentration of 20mg/mL and 10uL were spotted onto a silica gel 60 TLC plate (Sigma, #1057500001). The TLC plate was placed in a latch-lock chamber containing 95:5 chloroform:methanol solvent. After development, the plates were sprayed with a solution of 0.2% anthrone (Sigma, #319899) in H_2_SO_4_ and dried with a heat gun.

### PGL and SL-1 of bacterial origin

For initial experiments, PGL extracted from *Mycobacterium canettii* was obtained from BEI Resources (#NR-36510). Likewise, SL-1 extracted from *M. tuberculosis* was from BEI Resources (#NR-14845). All biologic molecules were stored at −20°C until use, at which time they were resuspended in DMSO.

### Organic synthesis of PGLs and PGL aglycon analogues

The synthesis of the phenolic glycolipids followed a modular strategy^35,98^ in which the required iodoglycan^36,99^, carrying protecting groups on the different hydroxyl functions that can be removed through a single hydrogenation step, was coupled with a phthiocerol alkyne in a Sonogashira cross-coupling. Next, the two mycocerosic acids were introduced through a Steglich esterification reaction. Global deprotection by hydrogenation over Pd/C then furnished the desired PGLs.

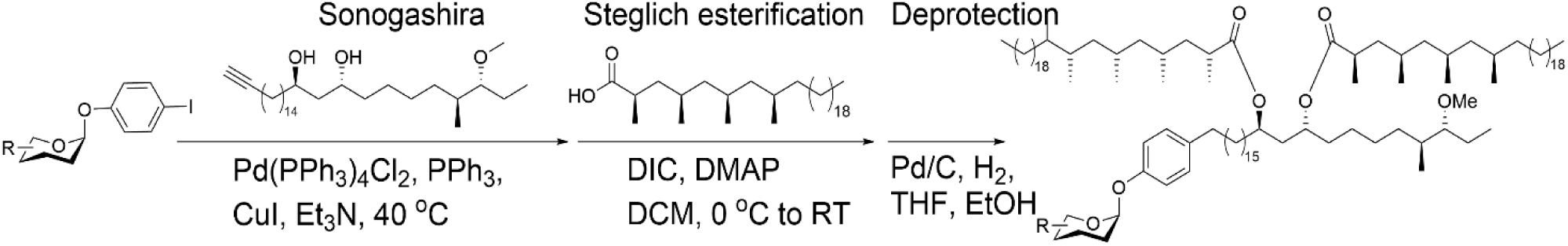

### General procedure Sonogashira cross coupling

Iodoaryl glycoside (1.0 eq) was dissolved in freshly distilled NEt_3_ (0.05 M) together with alkyne (1.2 – 3 eq). A mixture of Pd(PPh_3_)_2_Cl_2_, PPh_3_ and CuI (ratio 1:1:2) was dissolved in freshly distilled NEt_3_ and was stirred for 15 minutes at 40°C. Of this cocktail, enough was added to the sugar/alkyne mixture to amount to 0.05 eq Pd(PPh_3_)_2_Cl_2_, 0.05 eq PPh_3_ and 0.1 eq CuI. The reaction was allowed to stir at 40°C until the starting material was completely consumed, as indicated by TLC (2-16 h). The solvent was then removed under a stream of N_2_. The crude material was then transferred to a silica column in toluene and the column was flushed with toluene. Thereafter the product was purified by means of column chromatography.

### General esterification procedure with mycocerosic acid

Starting material (1.0 eq) was dissolved in dry DCM (0.05 M) together with mycocerosic acid (3.0 eq) and DMAP (9 eq). The resulting mixture was cooled to 0 °C after which DIC (6 eq) was added. The reaction was allowed to stir for 16 hours while warming to room temperature, after which it was warmed to 40°C and stirred for a further 5 hours. The reaction mixture was then diluted with Et_2_O and the organic layer was washed 1 M HCl, sat. aq. NaHCO_3_ and brine, dried with MgSO_4_ and concentrated *in vacuo*. Thereafter the product was purified by means of column chromatography.

### General hydrogenation procedure

Starting material (1.0 eq) was dissolved in a mixture of THF and EtOH (1:1, 0.007 M) and the solution was purged with N_2_. Pd/C (10%, 1.0 eq) was then added to the solution and the resulting mixture was purged with H_2_. The reaction was left to stir under H_2_ atmosphere until TLC revealed complete conversion of the starting material and reaction intermediates to a single low running spot (DCM-MeOH 19:1). The reaction mixture was then purged with N_2_ and filtered over celite and rinsed with acetone. Thereafter the product was purified by means of column chromatography. Extended methods and validation of synthesized PGL analogs are included in supplemental dataset 1.

### Neuronal cell line

The mouse nociceptive neuronal cell line, MED17.11^100^, was generously provided by M. Nassar (University of Sheffield). Cells were propagated in DMEM F12/Glutamax (Gibco, #10565018) supplemented with 10% fetal bovine serum (FBS) (Gibco, #16000044), 0.5% penicillin/streptomycin (Gibco, #15140122), 5 ng/mL recombinant mouse interferon gamma (R&D systems, #485-MI), and 0.5% chick embryo extract (US Biological, #NC1202490). Undifferentiated MED17.11 cells were cultured at 33°C and 5% CO_2_ until differentiation. For differentiation, cells were plated onto 35mm glass-bottom dishes (MatTek, #P35G-1.5-14-C) at a density of 2×10^4^ cells/dish in supplemented differentiation media at 37°C and 5% CO_2_ for 7 days with media change every 2 days. MED17.11 differentiation media consisted of DMEM F12/ Glutamax (Gibco, #10565018), 10% FBS (Gibco, #16000044), 0.5% penicillin/streptomycin (Gibco, #15140122), 10 ng/mL FGF basic (R&D systems, #3139FB025), 10 ng/mL GDNF (Sigma, #SRP3200), 100 ng/mL β-NGF (R&D systems, #1156-NG-100), 25 µM forskolin (R&D systems, #10-995-0) and 5 µg/mL Y-27632 (R&D systems, #12-545-0). For the first 2 days of differentiation, 0.5 mM of dibutyryl cAMP sodium salt (Sigma, #D0627) was added to the supplemented differentiation media.

### Animal Studies

Protocols for neuronal isolation from mice and guinea pig cough studies were reviewed and approved by the Institutional Animal Care and Use Committee at the University of Texas at Dallas and the University of Texas Southwestern Medical Center.

### Primary mouse neuron isolation and culture Dorsal Root Ganglia

Primary mouse dorsal root ganglia (DRG) neurons were dissected and cultured as described ^101^. In brief, C57BL/6J mice (JAX) were sacrificed and DRG were dissected from the intervertebral foramina and placed in media containing DMEM (Gibco, #10565018), 10% FBS (Gibco, #16000044), and 0.5% penicillin/streptomycin (Gibco, #15140122), on ice. Ganglia were then enzymatically digested in 1.25mg/mL collagenase A (Sigma, #10103578001) and 2.5 mg/mL dispase II (Sigma, #D4693) for 40 mins at 37°C with shaking. Dissociated tissue was then centrifuged at 1000 rpm for 5 mins and washed with DMEM media. The cell pellet was carefully resuspended in supplemented neurobasal-A media (Gibco, #10888022) containing 1x B-27 supplement (Thermo, 17504044), 1x GlutaMax (Thermo, #35050061), 1% penicillin/streptomycin, 50 ng/mL β-NGF (R&D systems, #1156-NG-100), 2 ng/mL GDNF (Sigma, #SRP3200), and 10 µM cytosine beta-D-arabinofuranoside hydrochloride (Sigma, #C6645). The DRGs were dissociated by aspirating and ejecting the solution several times through a 1mL syringe attached to an 18-gauge needle followed by 25- and 27-gauge needles until a single-cell suspension was achieved. Cells were then plated on 35mm glass-bottom dishes (MatTek, #P35G-1.5-14-C) that were pre-coated with 10 µg/mL laminin (Sigma, #L2020) and incubated at 37°C and 5% CO_2_ for 1-2 days. DRGs from 3-4 mice were combined to generate approximately 5-6 dishes of cells per experiment.

### Nodose Ganglia

Nodose tissues for culture were dissected from adult male and female ICR CD-1 mice and placed in Hanks’ balanced salt solution without calcium or magnesium (Sigma, #H6648). Nodose ganglia from 16 mice (equal numbers of male and female animals) were pooled together to create cultures for downstream processing. The nodose ganglia were then enzymatically digested using collagenase A and collagenase D (each 1 mg/ml, Roche, 45-11088858001) with papain (30 U/m, Roche, 45-10108014001) for 20 min at 37°C. Following digestion, the ganglia were triturated in 1 ml of Hanks’ balanced salt solution. The solution was then passed through a 70-μm cell strainer to remove debris. The isolated cells were resuspended in Dulbecco’s modified Eagle’s medium/F12/GlutaMAX (Gibco, #10565018) culture media supplemented with 10% fetal bovine serum (Hyclone, H30088.03) and 1% penicillin/streptomycin (Gibco, 15070-063). Cells were plated on precoated poly-d-lysine dishes (MatTek, P35GC-1.5-10-C) that were coated with laminin (Sigma-Aldrich, L2020). The plated cells were then incubated for 2 hours to allow for cell adhesion. The cells were then supplemented with the same culture media as described above but with the addition of NGF (10 ng/ml; Sigma, 01-125) and 5-fluoro-2’-deoxyuridine (3 μg/ml) + uridine (7 μg/ml) (FRD + U; Sigma-Aldrich, 856657) and incubated at 37°C with 5% CO_2_ for 48 hours before use.

### Live-cell neuronal calcium imaging

MED17.11 and primary mouse DRG neurons plated on 35mm dishes were loaded with 5 µg/mL of Fura-2 AM calcium indicator dye (Thermo, #F1221) in loading buffer containing Hank’s Balanced Salt Solution (HBSS) with phenol red (Thermo, #14170120), 2.5 mg/mL bovine serum albumin (Thermo, #BP1600), and 2 mM calcium chloride (Sigma, #10043-52-4). Cells were incubated with Fura-2 for 20 minutes and then placed in a solution of HBSS (no phenol red) with 2% HEPES buffer (Cytiva, #SH3023701) for 10 minutes prior to live-cell imaging. Nodose cultures were loaded with Fura-2 AM (1 μg/ml; Thermo, #F1221) for 1 hour then transferred to a normal bath containing 135 mM NaCl, 5 mM KCl, 10 mM Hepes, 1 mM CaCl_2_, 1 mM MgCl_2_, and 20 mM glucose, adjusted to pH 7.4 and osmolarity 300 ± 5 mOsm with N-methyl-glucamine (NMDG). Calcium imaging was performed using an Olympus IX73 microscope with MetaFluor Fluorescence Ratio imaging software. Cells were imaged every second at 340/380 nm excitation wavelengths and 510 nm emission wavelength and 340/380 ratiometric data were exported for downstream analysis.

For each cell dish, a 50 second baseline was first recorded following the addition of a vehicle control (1% DMSO) for 50-100 seconds. Next, test compounds or vehicle (DMSO) were added with continuous recording for a minimum of 100 seconds or until response reached baseline. Finally, a positive control of 200-400 nM capsaicin (Sigma, #404-86-4) was added for 30-100 seconds. Ratio data was analyzed through an R script to determine the maximum change in fluorescence of our compound of interest compared to the vehicle over the baseline fluorescence intensity (Max ΔF/F_0_). All data points shown are from individual cells with a capsaicin response >40% (MED17.11) or >10% (primary DRG and nodose).

### Inhibitor treatment and ATP release

To measure release of extracellular ATP, MED17.11 neurons were cultured in a 96 well plate (10,000 cells/well) and allowed to differentiate for 7 days. Following differentiation, cells were washed with HBSS and 100µL of HBSS was added to each well. Designated wells were stimulated with DMSO, PGL (100 nM) or KCl (50 mM) for 3 minutes. Supernatants were immediately collected after stimulation and ATP was measured using a luminescent ATP determination kit (Thermo, #A22066) based on manufacturer instructions.

For inhibitor treatment, MED17.11 neurons were loaded with Fura-2 calcium indicator dye as described above. Prior to calcium imaging, cells were treated with vehicle (DMSO) or inhibitor 5 U/mL apyrase (Sigma, #A6535), 10 µM carbenoxolone (Sigma, #C4790), or 10 µM gefapixant (Selleckchem, #S6664) for 10 or 20 minutes. The cells were then used for calcium imaging with the experimental design described for live-cell neuronal calcium imaging. Exposure to ATP (Sigma, #A7699) (2 µM) was used as a positive control for inhibitor treatment.

### Human Dorsal Root Ganglia

Human DRG (hDRGs) tissue was recovered as described previously^102^. All human tissue procurement procedures and ethical regulations were approved by the Institutional Review Board at the University of Texas at Dallas under protocol Legacy-MR-15-237. DRGs were procured from organ donors through a collaboration with the Southwest Transplant Alliance, an organ procurement organization (OPO) in Texas. The Southwest Transplant Alliance obtain informed consent for research tissue donation from first-person consent (driver’s license or legally binding document) or from the donor’s legal next of kin. Ethical oversight of OPOs, including the Southwest Transplant Alliance, is maintained by several federal agencies, such as the Health Resources and Services Administration (HRSA), Centers for Medicare and Medicaid Services (CMS), and the United Network for Organ Sharing (UNOS). These agencies ensure that donation practices comply with ethical guidelines, including informed consent and donor autonomy. Following isolation in the operating room at the organ recovery site, hDRGs were maintained in artificial cerebrospinal fluid (ACSF) on ice consisting of: 95 mM NMDG, 2.5mM KCL, 1.25mM NaH_2_PO_4_, 30mM NaHCO_3_, 20mM HEPES, 25mM

Glucose, 5mM Ascorbic acid, 2mM Thiourea, 3mM Sodium pyruvate, 10mM MgSO_4_, 0.5mM CaCl_2_, 12mM N-acetylcysteine. Upon returning to the lab, hDRG tissue was transferred to fresh ACSF solution on ice and carefully trimmed and cut into small pieces to enable better digestion. The tissue pieces were then transferred to a tube containing 1mL of 10mg/ml Stemxyme 1, Collagenase/Neutral Protease (Dispase, Worthington Biochemical, #LS004107) and placed in a sideways-shaking water bath. The tissues were digested for 3-4 hours with regular trituration with glass pipettes every hour. When DRG tissue was sufficiently digested, it was passed through a 100-µm cell strainer. The resulting suspension was carefully transferred to a 10% BSA gradient using a pipette and centrifuged at 900xg (9 acceleration, 7 deceleration) for 5 minutes. The supernatant was carefully discarded and the pellet resuspended in media containing BrainPhys® media (Stemcell technologies, #05790), 1% penicillin/streptomycin (Thermo, #15070063), 1% GlutaMAX® (United States Biological, #235242), 2% NeuroCult™ SM1 (Stemcell technologies, #05711), 1% N-2 Supplement (Thermo, #17502048), 2% HyClone™ Fetal Bovine Serum (Thermo, #SH3008803IR), 1: 1000 FrdU, and 10 ng/mL human β NGF. A 5 μL aliquot of the cell suspension was plated onto a dish, and the cells were counted to determine the appropriate media volume required to achieve the desired cell density for calcium imaging. The cells were plated on PDL-coated coverslips at a density of 150-200 cells per coverslip and incubated at 37°C (5% CO_2_) for 3 hours before flooding with media. The cells were incubated for 5 days with media change on alternate days prior to imaging. Specific donor details are provided in Supplementary Table 2.

### Human DRG calcium imaging

Calcium imaging of hDRGs was performed using Fluo-4 dye. Cells were loaded with a suspension of 1 mL HBSS containing 5 mL Fluo-4 resuspended in 82 µl of HBSS and 9.12 µl of pluronic acid. Cells were incubated with Fluo-4 for 1 hour. Following incubation, coverslips were transferred to a perfusion slide and external bath containing 125 mM NaCl, 4.2 mM KCl, 1.1 mM CaCl_2_, 29 mM NaHCO_3_, 20 mM Glucose, 2 mM MgSO_4_, 1 mM CaCl_2_, 2.5 mM KCl, 1.25 mM NaH_2_PO_4_, pH 7.4 with NMDG and osmolality of 300-305 mOSm. The cells were then treated, imaged, and analyzed in the same way as described for mouse nodose neurons except ratiometric measurements were not captured for Fluo-4.

### Electrophysiology

Recordings were done 3 or more days after plating. The external solution contained: 147 mM NaCl, 3 mM KCl, 1.25 mM NaH_2_PO_4_, 2 mM CaCl_2_,1. 25 mM MgCl_2_, 10 mM dextrose, and 10 mM HEPES (pH 7.4, 310 mOsm). Whole-cell patch clamp was performed using borosilicate capillaries pulled with a P-97 flaming-brown micropipette puller (Sutter Instruments). The pipettes had a resistance of 1.2-3.5 MΩ, when using an internal solution containing: 120 mM K-Glutamate, 2 mM KCl, 8 mM NaCl, 0.2 mM EGTA, 14 mM Na2-Phoshphocreatine, 2 mM Mg-ATP, 0.3 mM Na-GTP and 10 mM HEPES-K (pH 7.3, 295 mOsm). Recordings were obtained using an Axopatch 200B amplifier (Molecular Devices). Neurons were visualized using a Nikon Eclipse Ti inverted microscope equipped with Nikon Advanced Modulation Contrast. Acquisition was done at 10-20 kHz and data was filtered at 5 kHz and analyzed offline using Clampfit Analysis Suite 11 (Molecular Devices), GraphPad Prism 11, and Microsoft Excel 2015. In whole-cell configuration, cells were held in voltage clamp mode at near resting potential for at least 5 min to allow for dialysis of the pipette internal solution. Afterwards, when needed, cells were held at −60 mV (H-60) by injecting current of the appropriate sign and amplitude. The acute effect of PGL (1 nM) and of capsaicin (100-300 nM) on membrane potential and on evoked firing was evaluated using both step and ramp protocols separated by 10 seconds within the same sweep and repeated at 0.033 Hz and where the first 10 sweeps were recorded as baseline before adding PGL. In a different set of experiments, cells were recorded while at resting membrane potential (RMP), that is, no current was injected to hold them at a predetermined (holding) potential. After 1 min recording at RMP, 1 nM PGL was perfused and changes in RMP or spontaneous firing activity were monitored for 8-9 additional minutes. Afterwards PGL was washed out and a similar sequence was recorded for capsaicin.

### Guinea Pig Cough Studies

Male Hartley guinea pigs weighing >200g were purchased from Charles River Laboratories. Upon arrival, animals were housed in standard cages with alpha-dri bedding (Shepherd Specialty Papers) with irradiated pressed timothy hay cubes (Bioserv) and provided food and water *ad libitum.* All guinea pigs were given a 3-4 day acclimation period before any experimental procedures were performed.

For cough recording, naïve guinea pigs were placed inside specialized whole body plethysmograph chambers (Data Sciences International) fitted with a pressure transducer and nebulizer for aerosol delivery of compounds into the chamber. Unrestrained animals were first allowed to acclimate inside the chambers for 5 minutes, followed by 10-minute nebulization of vehicle (10% methanol in PBS), 1mL of Mtb lipid extract (20mg/mL), or purified PGL (60 or 250 µg/mL). All animals were treated with a positive control cough agonist 0.4-0.8M citric acid (Thermo, #036665.36). Following the nebulization period, animals were acclimated inside the chambers for another 10 minutes with continuous pressure recording for a total cough recording time of 20 minutes. Cough recordings were performed on the same animals over the experimental course with a 2-day waiting period between treatments to prevent tachyphylaxis. Using the companion software for analysis (Buxco FinePointe), primary cough data was recorded as the bias flow of the system, the slope of the bias flow, and the transition from the compressive to expulsive phase of a cough (delta half peak crossing). Individual coughs were counted for each animal both by automated software and manual counts of blinded data.

### Statistical Analysis

Statistical comparisons between groups were performed using GraphPad Prism 10.4.0 software. Comparisons between multiple groups were assessed by Kruskal-Wallis test while two groups were compared by Mann-Whitney. For guinea pig cough experiments, paired data sets were analyzed by Friedman test. EC_50_ of PGL compounds were calculated by nonlinear regression analysis. For all statistical tests, differences were considered significant at p<0.05.

## ACKNOWLEDGMENTS

The authors would like to thank all members of the Shiloh lab for their support and constructive feedback on the manuscript. We also thank members of the Animal Resources Center at UT Southwestern for providing support with all animal welfare and husbandry. We thank the Southwest Transplant Alliance for recovery of tissues from organ donors and we thank the organ donors and their families for their gift. This work was supported by NIH T32 AI007520 (K.F.N), R01 AI158688 (M.U.S), P01 AI159402 (M.U.S) and U19NS130608 (T.J.P). Financial support was also provided by the Burroughs Wellcome Fund 1017894 (M.U.S.), Welch Foundation I-1964-20240404 (M.U.S.), the Nederlandse Organisatie voor Wetenschappelijk Onderzoek (NWO), Toppunt 15.002 (J.D.C.C.) and VICIVI.C.182.020 (J.D.C.C.). Michael Shiloh would also like to acknowledge support from the Disease Oriented Clinical Scholar program at UT Southwestern.

## AUTHOR CONTRIBUTIONS

Conceptualization, K.F.N, L.L.W. and M.U.S. Formal Analysis, K.F.N, L.L.W. and M.U.S. Funding Acquisition, J.D.C.C., T.J.P and M.U.S. Investigation, K.F.N, L.L.W., D.K.N, F.dJ.E-B, I.M.O and K.C.R. Project Administration, K.F.N and M.U.S. Resources, J.H.M.vD, V.V.F.V.. J.D.C.C, and T.J.P. Validation, K.F.N, J.H.M.vD, V.V.F.V, K.C.R and J.D.C.C. Software, C.P.X. Supervision, J.D.C.C., T.J.P. and M.U.S. Writing-original draft, K.F.N and M.U.S. Writing-Review and Editing, all authors.

---

**Supplemental Figure 1: Detection of Nociceptive Lipids from Mtb Extracts. (**A) Sulfolipid-1 detected by mass spectrometry from lipid extracts of HN878 and Erdman strains. (B) PGL detection by TLC (95:5 chloroform:methanol) and 0.2% anthrone in H_2_SO_4_ stain. Arrow= PGL

**Supplemental Figure 2: Activation of Neurons by Organically Synthesized PGL Analogs.** (A) Synthetic PGL analogs with modifications made to methyl groups on the saccharide chains. R=PDIM. (B) Neuronal activation of all three PGL analogs (1nM) compared to DMSO vehicle control. *P-*values calculated by Kruskal-Wallis.

**Supplementary Figure 3: Variations in PGL Saccharide Chain Impact Activity.** (A) Structure of organically synthesized PGL from *Mycobacterium haemophilum*. (B) Level of activation following MED17.11 stimulation with PGL *haemophilum* (1nM). (C) Organically synthesized analogs of mycobacterial PGLs containing phenol ring and corresponding saccharide chain. (D) Neuronal activation of phenolic analogs at 1nM concentration on MED17.11 cells. *P-*values calculated by Kruskal-Wallis and Mann-Whitney

**Supplemental Figure 4: Cough Analysis by Positive Control Agonist Citric Acid.** (A) Coughs recorded from cohort 1 (Figure 5B) guinea pigs upon exposure to 0.8M citric acid. (B) Coughs from cohort 2 (Figure 5C) of guinea pigs exposed to 0.4M citric acid. *P-*values calculated by Friedman test

