## Supplementary figures and images for "Mycobacterial Phenolic Glycolipid Triggers ATP-Mediated Neuronal P2X3 Signaling and Cough"

### Supplemental Figure 1

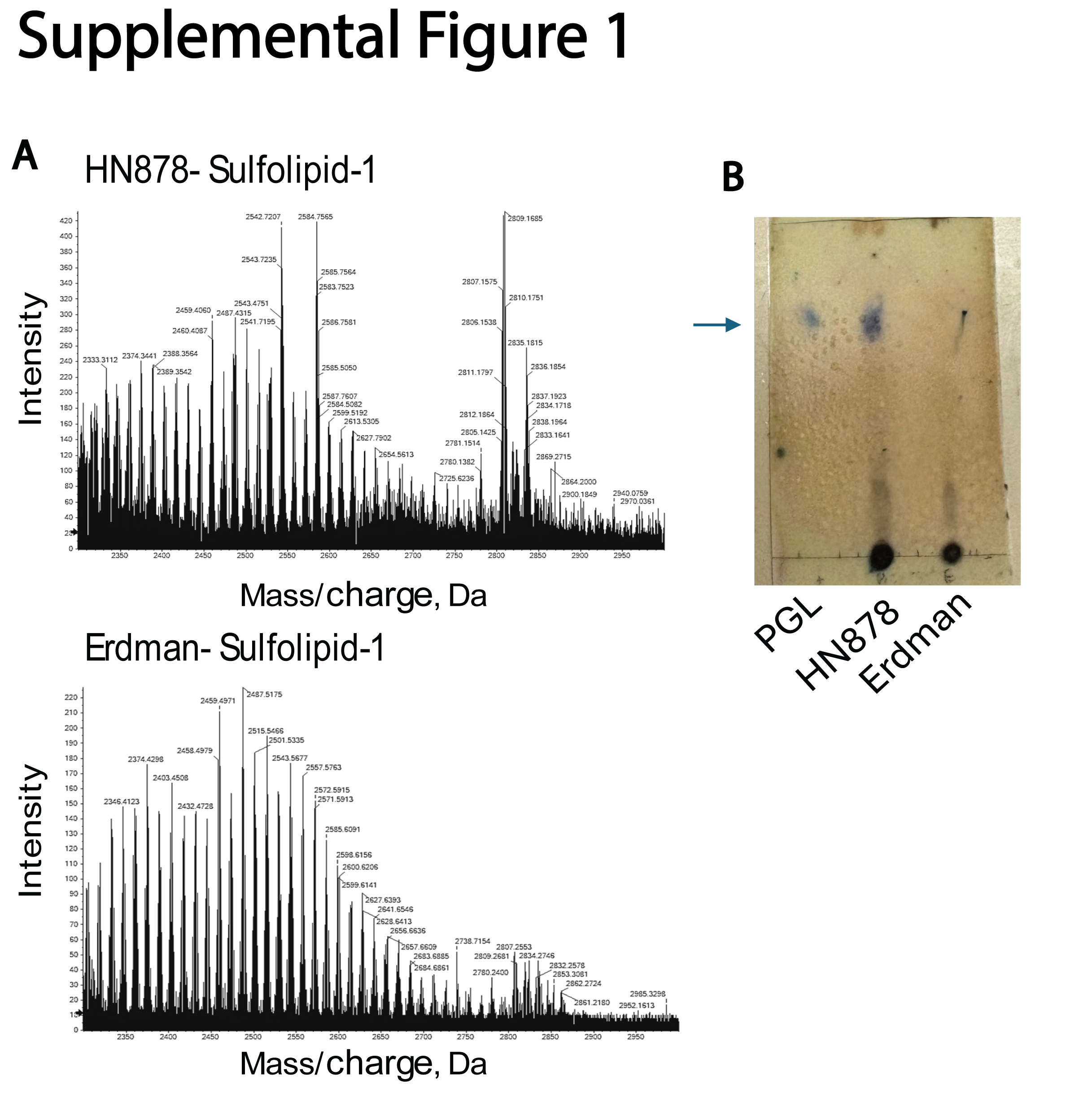

### Supplemental Figure 2

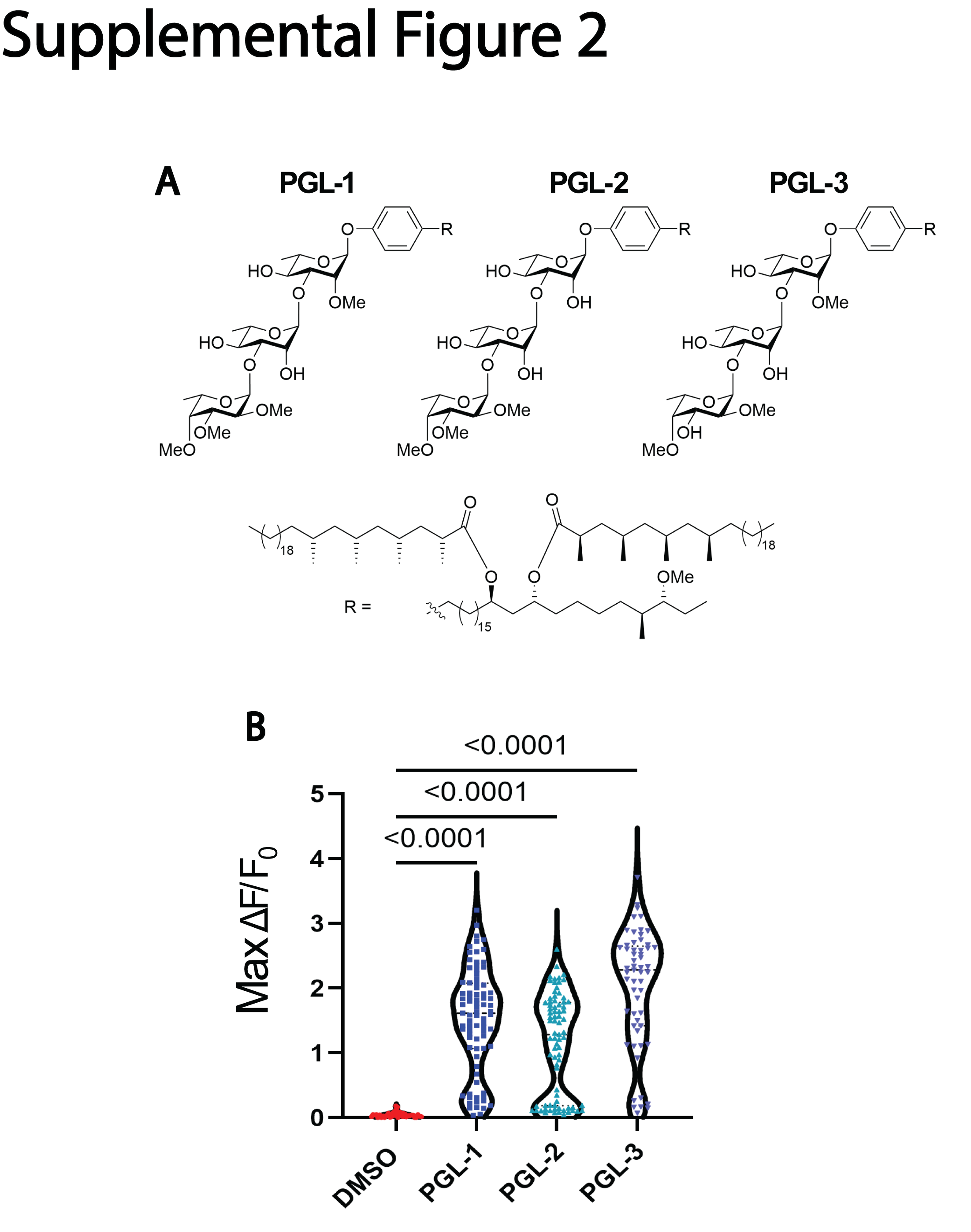

### Supplemental Figure 3

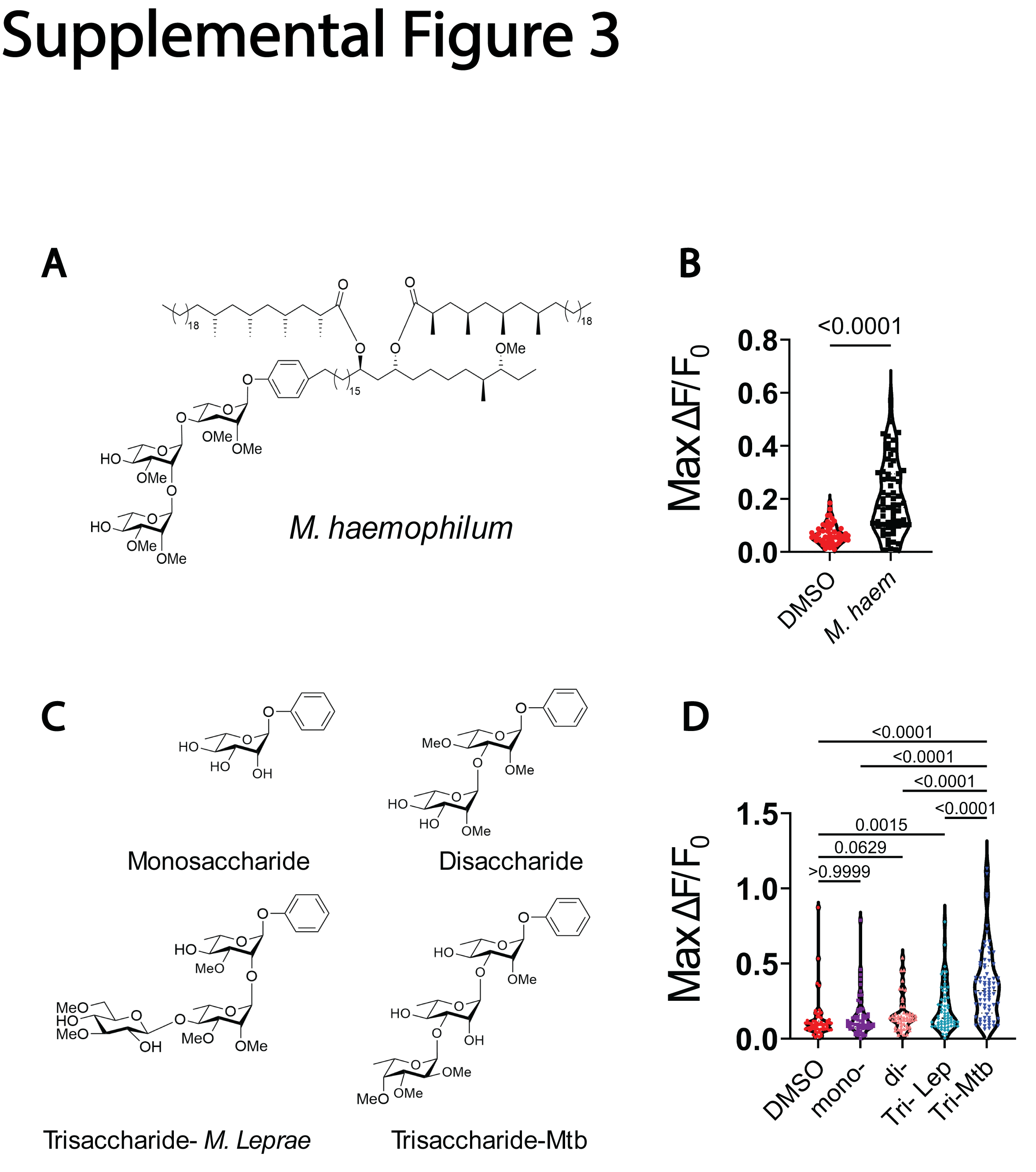

### Supplemental Figure 4

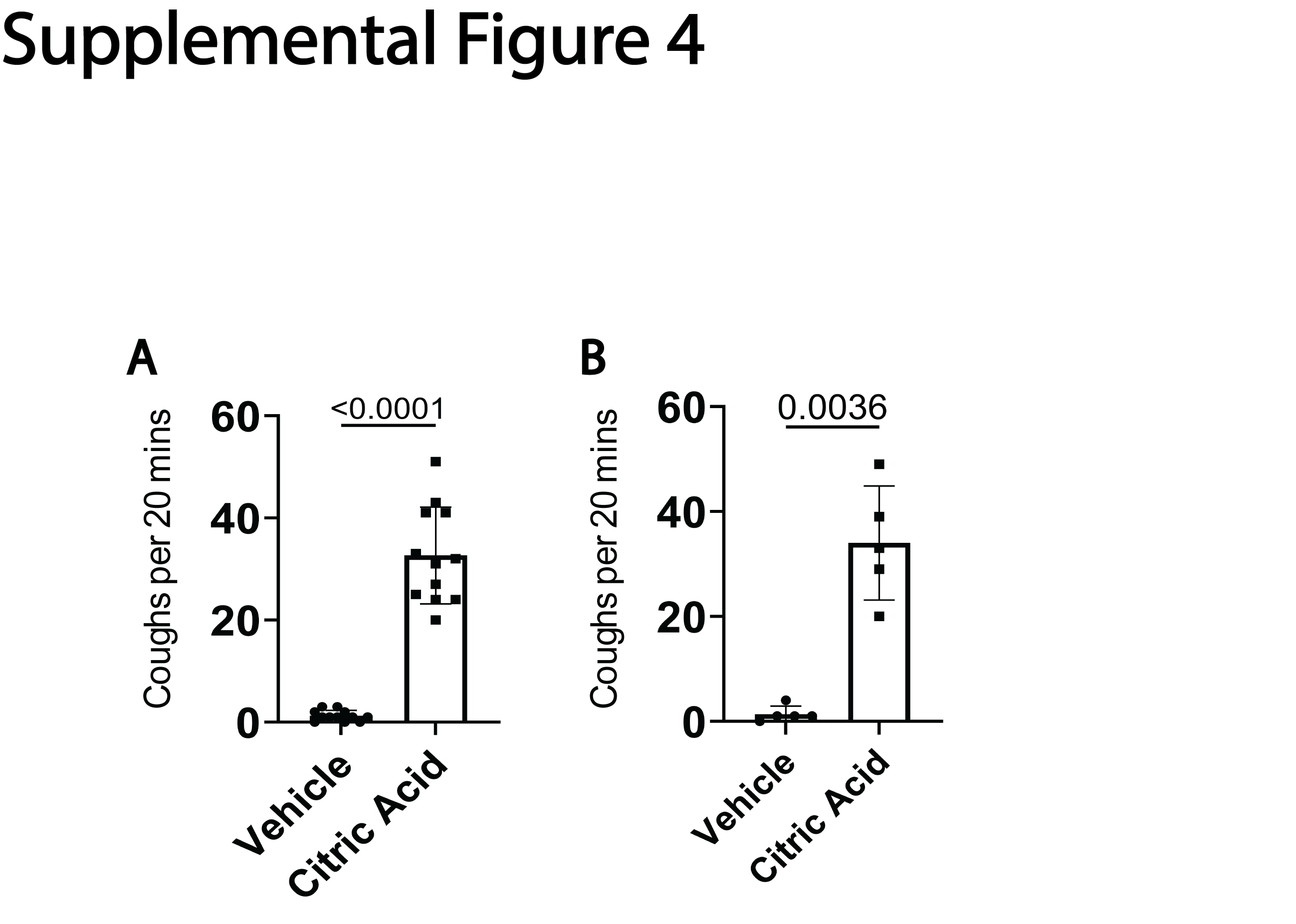

### Supplemental Table 1

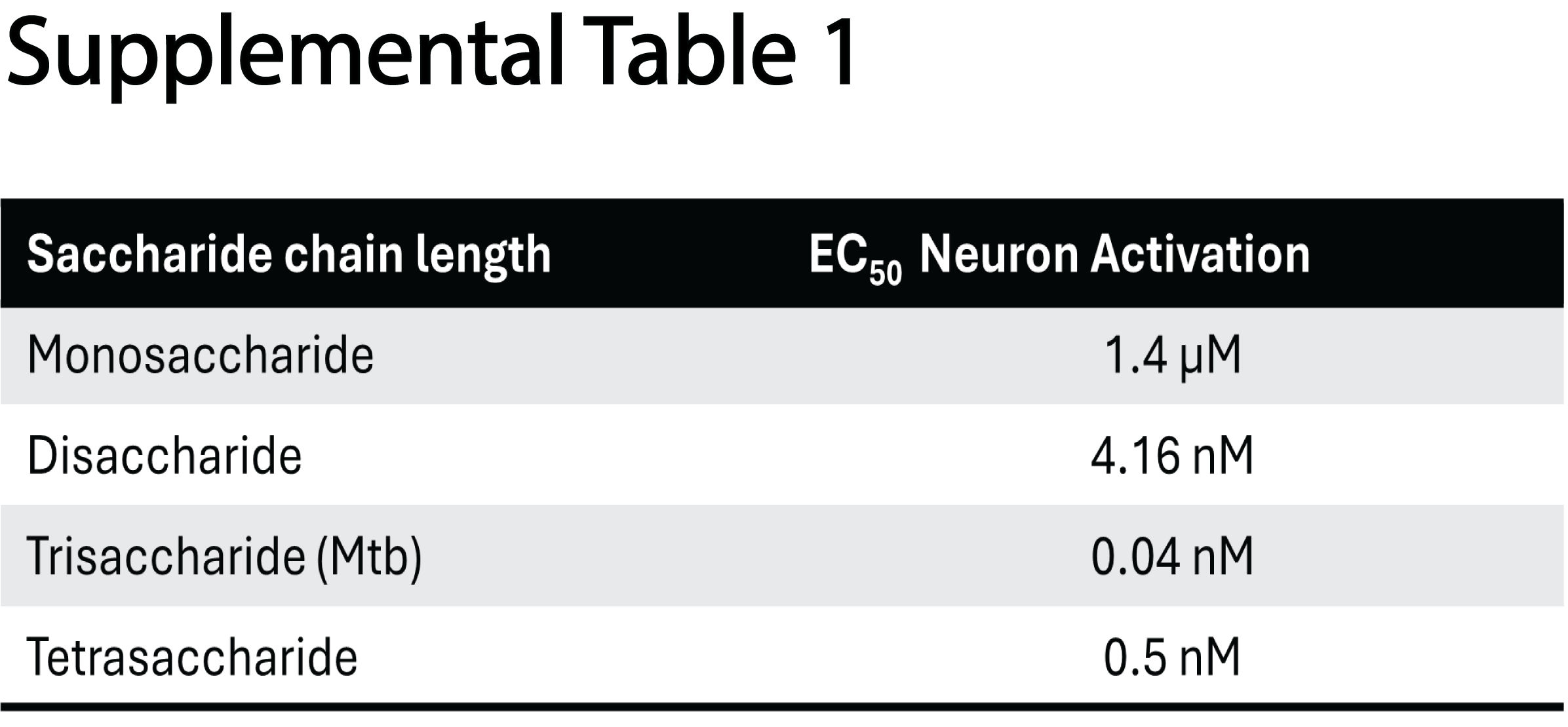

### Supplemental Table 2

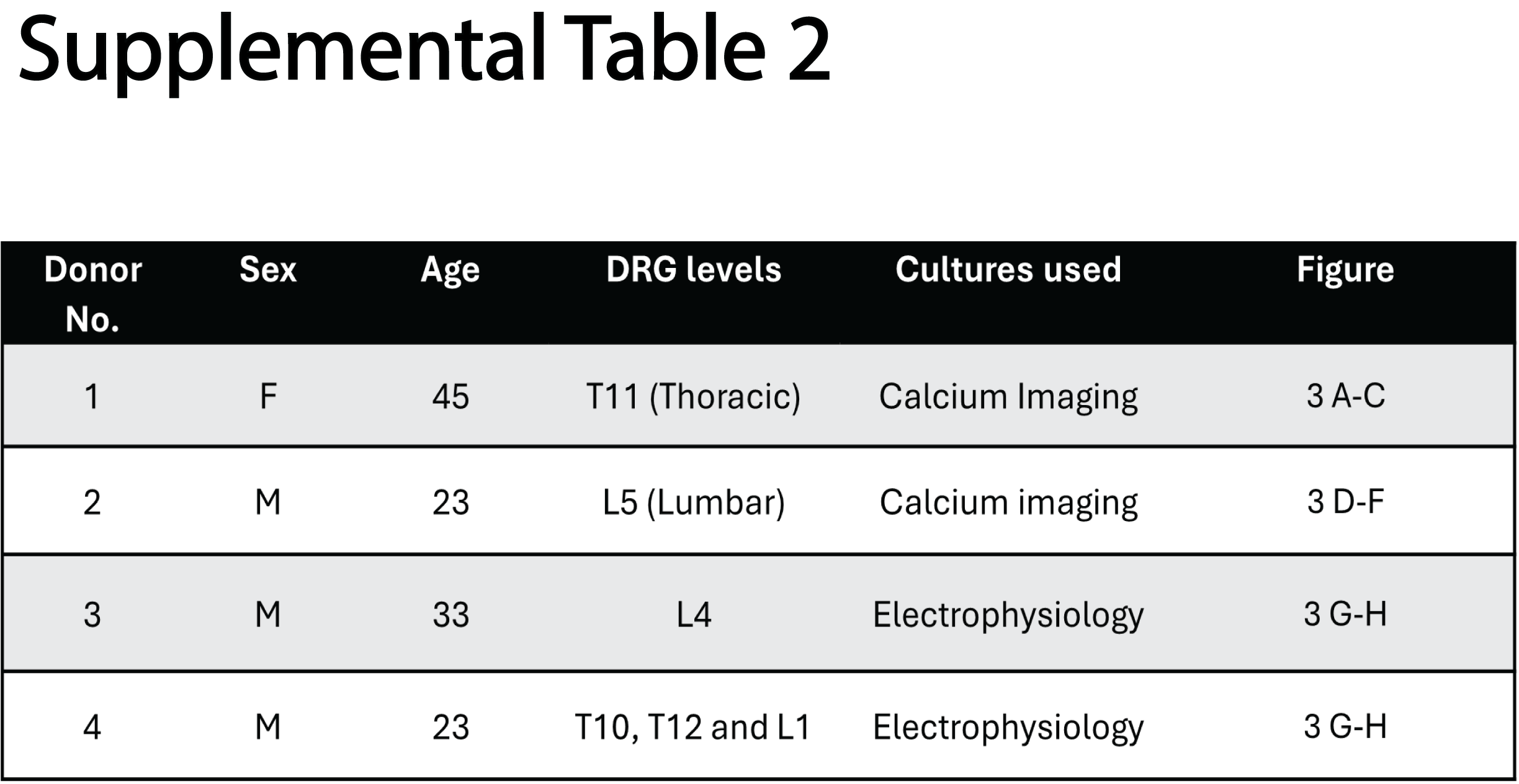
