## Supplemental dataset for "Mycobacterial Phenolic Glycolipid Triggers ATP-Mediated Neuronal P2X3 Signaling and Cough"

#### Synthesis of PGLs and aglycon analogues

##### General two-step deiodination and hydrogenation procedure

Iodoaryl glycoside (1.0 eq) and $NH_4OAc$ (3.3 eq) were dissolved in EtOH (0.01 M). The solution was sparged with Ar for 30 min. Pd/C (10 wt%, catalytic amount) was added and the resulting mixture was sparged with Ar for 30 min followed by sparging with $H_2$ for 30 min. The reaction mixture was stirred for 3 days under a $H_2$ atmosphere. Progress of the reaction was monitored with TLC (9:1 DCM/MeOH). After completion, the reaction mixture was sparged with Ar for 30 min and filtered over a Whatman filter. The resulting mixture was extracted with DCM (3x). The combined organic layers were washed with sat. aq. $Na_2S_2O_3$, dried over $MgSO_4$, filtered and concentrated *in vacuo*. The crude was redissolved in 1:1 THF/EtOH (0.007 M). The solution was sparged with Ar for 30 min. Pd/C (10 wt%, 1 eq) was added and the resulting mixture was sparged with Ar for 30 min followed by sparging with $H_2$ for 30 min. The reaction mixture was stirred for 2 h under a $H_2$ atmosphere. Progress of the reaction was monitored with TLC (9:1 DCM/MeOH). After completion, the reaction mixture was sparged with Ar for 30 min and filtered over a Whatman filter. The resulting mixture was concentrated *in vacuo*. The product was purified by means of flash column chromatography.

### Characterization data

#### MTb PGL-1

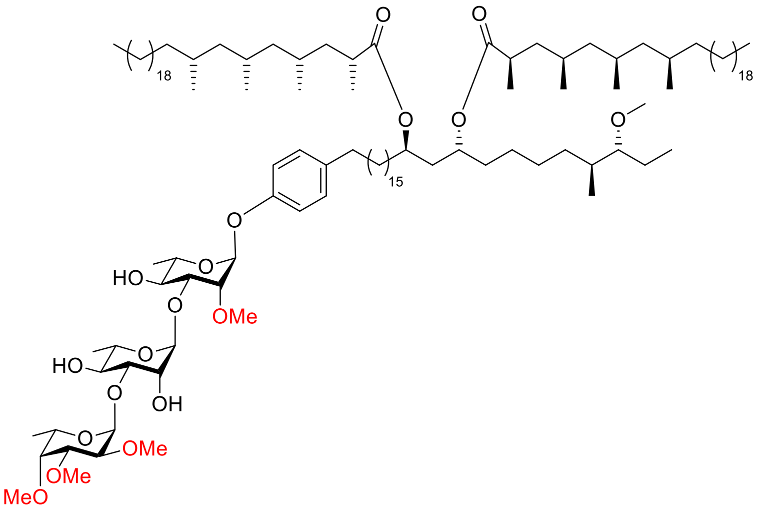


$[\alpha]_D^{25}$ = -48.4 (c = 0.5, $CHCl_3$). $^1$H-NMR (400 MHz) δ: 7.10 (d, 2H, *J* = 8.4 Hz, $CH_{arom}$); 6.99 (d, 2H, *J* = 8.8 Hz, $CH_{arom}$); 5.51 (d, 1H, *J* = 1.6 Hz, H-1); 5.15-5.14 (m, 2H, H-1', H-1"); 4.84 (quint, 2H, *J* = 6.4 Hz, $CH_{Phth}$); 4.11 (s, 1H, H-2'); 4.07-4.03 (m, 2H, H-3, H-5"); 3.98-3.91 (m, 1H, H-5'); 3.82-3.74 (m, 3H, H-2, H-3', H-5); 3.70-3.64 (m, 4H, H-2", H-3", H-4, H-4'); 3.61 (s, 3H, $OCH_3$); 3.58 (s, 3H, $OCH_3$); 3.52 (s, 3H, $OCH_3$); 3.49 (s, 3H, $OCH_3$); 3.48 (d, 1H, *J* = 1.6 Hz, H-4"); 3.33 (s, 3H, $OCH_3$); 2.88-2.83 (m, 1H, $CH_{Phth}$); 2.58-2.48 (m, 4H, $CH_{2,Phth}$, $CH_{Myc}$); 2.27 (bs, 1H, *OH*); 2.16 (bs, 1H, *OH*); 1.77-0.81 (m, 203H, $CH_{Phth}$, $CH_{2,Phth}$, $CH_{3,Phth}$, $CH_{Myc,}$ $CH_{2,Myc}$, $CH_{3,Myc}$, H-6, H-6', H-6"). $^{13}$C-APT NMR (101 MHz) δ: 176.2, 176.1 ($CO_{Myc}$); 154.7, 137.0 ($C_{q,arom}$); 129.5, 116.3 ($CH_{arom}$); 102.3 (C-1"); 100.9 (C-1'); 95.0 (C-1); 86.8 ($CH_{Phth}$); 83.3 (C-3'); 81.1 (C-3"); 80.2 (C-2); 80.1 (C-3); 79.1 (C-4"); 78.9 (C-2"); 71.9 (C-4'); 71.8 (C-4); 71.3 (C-2'); 70.4 ($CH_{Phth}$); 69.2 (C-5); 68.8 (C-5'); 67.6 (C-5"); 62.1, 60.4, 58.7, 57.9, 57.5 ($OCH_3$); 45.6, 45.4 ($CH_{2,Myc}$); 41.1, 38.6 ($CH_{2,Phth}$); 37.9, 37.9 ($CH_{Myc}$); 36.8 ($CH_{2,Myc}$); 34.9 ($CH_{Phth}$); 34.8, 32.8 ($CH_{2,Phth}$); 32.1 ($CH_{2,Myc}$); 31.9, 30.2 ($CH_{2,Phth}$); 30.1 ($CH_{Myc}$); 29.9, 29.9, 29.8, 29.7, 29.5 ($CH_2$); 28.2 ($CH_{Myc}$); 27.6 ($CH_{2,Phth}$); 27.3 ($CH_{Myc}$); 27.1 ($CH_{2,Myc}$); 25.7, 25.3 ($CH_{2,Phth}$); 22.8 ($CH_{2,Myc}$); 22.5 ($CH_{2,Phth}$); 20.9, 20.6, 20.6 ($CH_{3,Myc}$); 18.6 ($CH_{3,Myc}$); 18.0, 17.9 (C-6 and C-6'); 16.8 (C-6"); 14.8 ($CH_{3,Phth}$); 14.3 ($CH_{3,Myc}$); 10.3 ($CH_{3,Phth}$). IR (thin film, $cm^{-1}$): 1020, 1043, 1095, 1229, 1259, 1379, 1460, 1508, 1731, 1736, 2853, 2923, 3420. HRMS calculated for $C_{121}H_{227}O_{18}$ 1969.68761 $[M+H]^+$; found 1969.68884.

#### MTb PGL-2

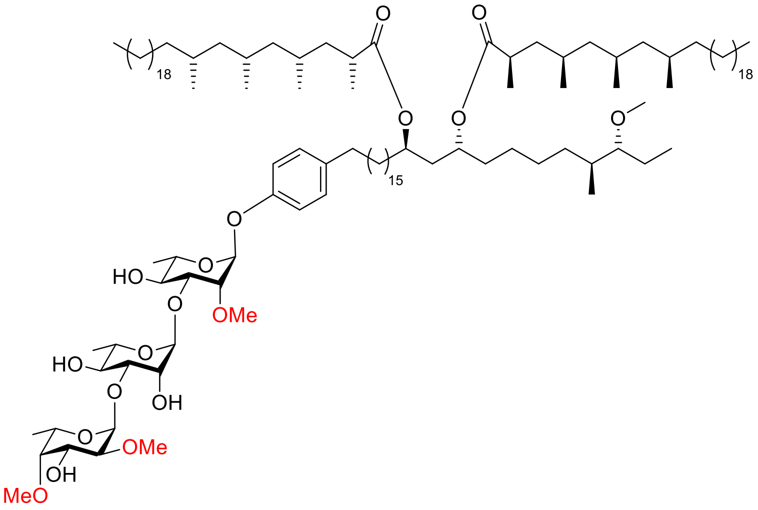

$[\alpha]_D^{25}$ = -44.7 (c = 1.0, $CHCl_3$). $^1$H-NMR (400 MHz) δ: 7.10 (d, 2H, *J* = 8.4 Hz, $CH_{arom}$); 6.99 (d, 2H, *J* = 8.8 Hz, $CH_{arom}$); 5.51 (d, 1H, *J* = 1.6 Hz, H-1); 5.18 (d, 1H, *J* = 3.6 Hz, H-1”); 5.13 (d, 1H, *J* = 1.2 Hz, H-1’); 4.84 (quint, 2H, *J* = 6.4 Hz, $CH_{Phth}$); 4.16-4.11 (m, 2H, H-2’, H-5”); 4.04 (dd, 2H, *J* = 3.0, 9.4 Hz, H-3, H-3”); 3.98-3.91 (m, 1H, H-5’); 3.81-3.74 (m, 3H, H-2, H-3’, H-5); 3.70-3.61 (m, 5H, H-4, H-4’, $OCH_3$); 3.58 (s, 3H, $OCH_3$); 3.50-3.47 (m, 4H, H-2”, $OCH_3$); 3.40 (d, 1H, *J* = 2.4 Hz, H-4”); 3.33 (s, 3H, $OCH_3$); 2.88-2.83 (m, 1H, $CH_{Phth}$); 2.58-2.48 (m, 4H, $CH_{2,Phth}$, $CH_{Myc}$); 1.77-0.81 (m, 191H, H-6, H-6’, H-6”, $CH_{Phth}$, $CH_{2,Phth}$, $CH_{3,Phth}$, $CH_{Myc,}$ $CH_{2,Myc}$, $CH_{3,Myc}$). $^{13}$C-APT NMR (101 MHz) δ: 176.2, 176.1 ($CO_{Myc}$); 154.7, 137.0 ($C_{q,arom}$); 129.5, 116.3 ($CH_{arom}$); 102.3 (C-1”); 99.9 (C-1’); 95.0 (C-1); 86.8 ($CH_{Phth}$); 83.0 (C-3’); 82.5 (C-4”); 80.2 (C-3); 80.1 (C-2”); 80.1 (C-2); 71.9 (C-4); 71.6 (C-4’); 71.2 (C-2’); 70.6 (C-3”); 70.4 ($CH_{Phth}$); 69.2 (C-5); 68.8 (C-5’); 67.6 (C-5”); 62.6, 59.8, 58.7, 57.7 ($OCH_3$); 45.6, 45.4 ($CH_{2,Myc}$); 41.1, 38.6 ($CH_{2,Phth}$); 37.9 ($CH_{Myc}$); 36.7, 35.3 ($CH_{2,Myc}$); 34.9 ($CH_{Phth}$); 34.8, 32.8 ($CH_{2,Phth}$); 32.1 ($CH_{2,Myc}$); 31.9, 30.2 ($CH_{2,Phth}$); 30.1 ($CH_{Myc}$); 29.9, 29.9, 29.8, 29.7, 29.5 ($CH_2$); 28.2 ($CH_{Myc}$); 27.6 ($CH_{2,Phth}$); 27.3 ($CH_{Myc}$); 27.1 ($CH_{2,Myc}$); 25.7, 25.3 ($CH_{2,Phth}$); 22.8 ($CH_{2,Myc}$); 22.5 ($CH_{2,Phth}$); 20.9, 20.6, 20.5, 20.5, 18.6 ($CH_{3,Myc}$); 18.0 (C-6); 17.9 (C-6’); 16.9 (C-6”); 14.8 ($CH_{3,Phth}$); 14.3 ($CH_{3,Myc}$); 10.2 ($CH_{3,Phth}$). IR (thin film, $cm^{-1}$): 1043, 1129, 1150, 1173, 1261, 1378, 1461, 1510, 1734, 2853, 2923, 3414. HRMS calculated for $C_{120}H_{225}O_{18}$ 1955.67196 $[M+H]^+$; found 1955.67295.

#### MTb PGL-3

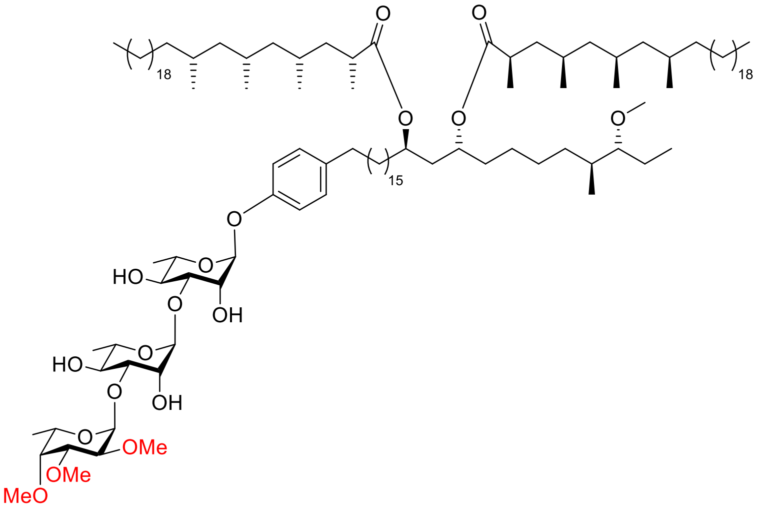


$[\alpha]_D^{25}$ = -47.8 (c = 1.0, $CHCl_3$). $^1$H-NMR (400 MHz) δ: 7.09 (d, 2H, *J* = 8.8 Hz, $CH_{arom}$); 6.97 (d, 2H, *J* = 8.4 Hz, $CH_{arom}$); 5.45 (d, 1H, *J* = 1.6 Hz, H-1); 5.20 (d, 1H, *J* = 1.2 Hz, H-1'); 5.16 (d, 1H, *J* = 3.2 Hz, H-1"); 4.84 (quint, 2H, *J* = 6.2 Hz, $CH_{Phth}$); 4.18 (dd, 1H, *J* = 2.0, 3.2 Hz, H-2); 4.12 (dd, 1H, *J* = 1.6, 3.2 Hz, H-2'); 4.10-4.02 (m, 2H, H-3, H-5"); 3.92-3.76 (m, 3H, H-3', H-5, H-5'); 3.71-3.64 (m, 4H, H-2", H-3", H-4, H-4'); 3.61 (s, 3H, $OCH_3$); 3.59 (s, 3H, $OCH_3$); 3.52 (s, 3H, $OCH_3$); 3.48 (d, 1H, *J* = 1.6 Hz, H-4"); 3.33 (s, 3H, $OCH_3$); 2.88-2.83 (m, 1H, $CH_{Phth}$); 2.58-2.48 (m, 4H, $CH_{2,Phth}$, $CH_{Myc}$); 1.77-0.81 (m, 192H, H-6, H-6', H-6", $CH_{Phth}$, $CH_{2,Phth}$, $CH_{3,Phth}$, $CH_{Myc}$, $CH_{2,Myc}$, $CH_{3,Myc}$). $^{13}$C-APT NMR (101 MHz) δ: 176.2 ($CO_{Myc}$); 154.3, 137.0 ($C_{q,arom}$); 129.4, 116.3 ($CH_{arom}$); 101.8 (C-1"); 101.0 (C-1'); 97.9 (C-1); 86.8 ($CH_{Phth}$); 83.2 (C-3'); 81.1 (C-3"); 79.7 (C-3); 79.0 (C-4"); 78.8 (C-2"); 72.2 (C-4); 71.6 (C-4'); 71.2 (C-2'); 70.8 (C-2); 70.4 ($CH_{Myc}$); 69.1 (C-5'); 68.8 (C-5); 67.7 (C-5"); 62.1, 60.4, 57.9, 57.5 ($OCH_3$); 45.6, 45.4 ($CH_{2,Myc}$); 41.1, 38.6 ($CH_{2,Phth}$); 37.9 ($CH_{Myc}$); 36.8, 35.3 ($CH_{2,Myc}$); 34.9 ($CH_{Phth}$); 34.8, 32.8 ($CH_{2,Phth}$); 32.1 ($CH_{2,Myc}$); 31.8, 30.2 ($CH_{2,Phth}$); 30.1 ($CH_{Myc}$); 29.9, 29.9, 29.8, 29.7, 29.5 ($CH_2$); 28.2 ($CH_{Myc}$); 27.6 ($CH_{2,Phth}$); 27.3 ($CH_{Myc}$); 27.1 ($CH_{2,Myc}$); 25.7, 25.3 ($CH_{2,Phth}$); 22.8 ($CH_{2,Myc}$); 22.5 ($CH_{2,Phth}$); 20.9, 20.6, 20.6, 18.6 ($CH_{3,Myc}$); 17.8 (C-6) 17.8 (C-6'); 16.8 (C-6"); 14.8 ($CH_{3,Phth}$); 14.3 ($CH_{3,Myc}$); 10.3 ($CH_{3,Phth}$). IR (thin film, $cm^{-1}$): 1045, 1090, 1228, 1172, 1229, 1261, 1378, 1457, 1511, 1734, 2853, 2922, 3441. HRMS calculated for $C_{120}H_{225}O_{18}$ 1955.67196 $[M+H]^+$; found 1955.67222.

#### ***M. leprae*** **PGL-I**

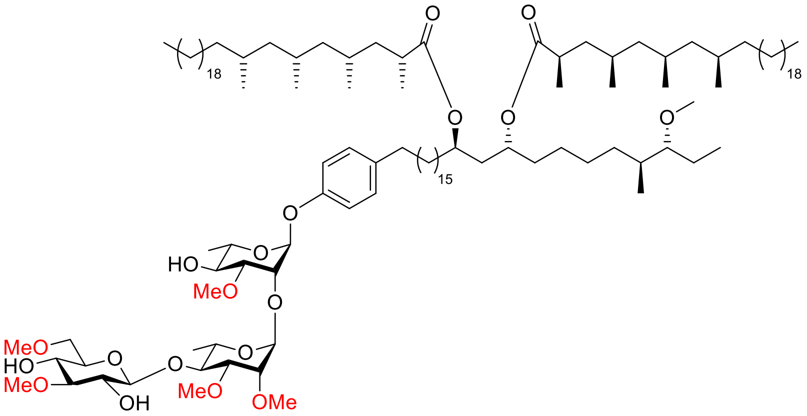


$[\alpha]_D^{25}$ = -25.2 (c = 1.0, $CHCl_3$). $^{1}$<u>H-NMR</u> (850 MHz) δ: 7.10 (d, 2H, *J* = 9.4 hz, $CH_{arom}$); 6.94 (d, 2H, *J* = 8.5 Hz, $CH_{arom}$); 5.43 (d, 1H, *J* = 1.7 Hz, H-1); 5.10 (d, 1H, *J* = 1.7 Hz, H-1'); 4.84 (quint, 2H, *J* = 6.4 Hz, $CH_{Phth}$); 4.41 (d, 1H, *J* = 7.7 Hz, H-1"); 4.22 (dd, 1H, *J* = 2.4, 2.8 Hz, H-2); 3.89 (s, 1H, 2"-O*H*); 3.77-3.74 (m, 3H, H-2', H-5, H-5'); 3.69-3.66 (m, 4H, H-3', $OCH_3$); 3.65-3.61 (m, 4H, H-3, H-4', H-6"); 3.58 (dt, 1H, *J* = 1.7, 9.4 Hz, H-4); 3.55-3.52 (m, 4H, H-4", $OCH_3$); 3.50 (s, 3H, $OCH_3$); 3.48 (s, 3H, $OCH_3$); 3.17 (t, 1H, *J* = 9.4 Hz, H-3"); ); 2.87-2.84 (m, 1H, $CH_{Phth}$); 2.80 (bs, 1H, 4"-O*H*); 2.56-2.51(m, 4H, $CH_{2,Phth}$, $CH_{Myc}$); 2.30 (bs, 1H, 4-O*H*); 1.77-0.81 (m, 190H, $CH_{Phth}$, $CH_{2,Phth}$, $CH_{3,Phth}$, $CH_{Myc,}$ $CH_{2,Myc}$, $CH_{3,Myc}$, H-6, H-6'). $^{13}$<u>C-APT NMR</u> (214 MHz) δ: 176.2, 176.1 ($CO_{Myc}$); 154.3, 137.0 ($C_{q,arom}$); 129.5, 116.1 ($CH_{arom}$); 105.6 (C-1"); 98.5 (C-1'); 97.4 (C-1); 86.8 ($CH_{Phth}$); 85.6 (C-3"); 81.8 (C-4'); 81.5 (C-3); 80.3 (C-3'); 75.6 (C-2'); 75.1 (C-2"); 74.1 (C-5"); 73.0 (C-6"); 72.2 (C-2); 71.9 (C-4); 71.4 (C-4"); 70.4, 70.4 ($CH_{Phth}$); 69.1 (C-5); 68.4 9C-5'); 60.7, 59.7, 59.2, 57.8, 57.5, 56.7 ($OCH_3$); 45.6, 45.6, 45.4 45.4 ($CH_{2,Myc}$); 41.1, 41.1, 38.5 ($CH_{2,Phth}$); 37.9, 37.9 ($CH_{Myc}$); 36.7, 35.3 ($CH_{2,Myc}$); 34.9 ($CH_{Phth}$); 34.8, 34.8, 32.8 ($CH_{2,Phth}$); 32.1 ($CH_{2,Myc}$); 31.9, 30.2 ($CH_{2,Phth}$); 30.0 ($CH_{Myc}$); 29.9, 29.9, 29.9, 29.9, 29.8, 29.8, 29.7, 29.7, 29.6, 29.5 ($CH_2$); 28.1 ($CH_{Myc}$); 27.6 ($CH_{2,Phth}$); 27.3 ($CH_{Myc}$); 27.1 ($CH_{2,Myc}$); 25.7, 25.3 ($CH_{2,Phth}$); 22.8 ($CH_{2,Myc}$); 22.4 ($CH_{2,Phth}$); 20.9, 20.6, 20.5, 20.5, 18.6 ($CH_{3,Myc}$); 18.6 ($CH_{3,Phth}$); 17.9 (C-6); 17.7 (C-6'); 14.8 ($CH_{3,Phth}$); 14.3 ($CH_{3,Myc}$); 10.3 ($CH_{3,Phth}$). <u>IR</u> (thin film, $cm^{-1}$): 1010, 1070, 1123, 1175, 1229, 1261, 1378, 1457, 1464, 1511, 1734, 2853, 2922, 3434. <u>HRMS</u> calculated for $C_{122}H_{229}O_{19}$ 1998.69476 $[M+H]^+$; found 1998.69683.

##### *M. kansasii* PGL K-I

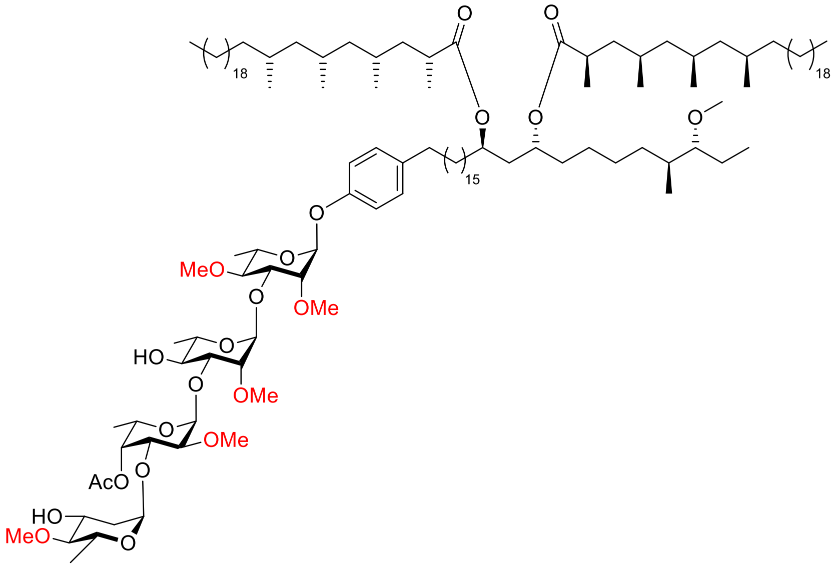


$[\alpha]_D^{25}$ = -38.8 (c = 1.0, $CHCl_3$). $^1$H-NMR (500 MHz) δ: 7.09 (d, 2H, *J* = 8.5 Hz, $CH_{arom}$); 6.97 (d, 2H, *J* = 8.5 Hz, $CH_{arom}$); 5.47 (d, 1H, *J* = 2.0 Hz, H-1); 5.18-5.17 (m, 2H, H-1’, H-4’’); 5.09-5.08 (m, 2H, H-1’’, H-1’’’); 4.84 (quint, 2H, *J* = 6.5 Hz, $CH_{Phth}$); 4.29 (q, 1H, *J* = 6.8 Hz, H-5’’); 4.16-4.11 (m, 2H, H-3, H-3’’); 3.88-3.82 (m, 2H, H-3’’’, H-5’); 3.79-3.74 (m, 3H, H-3’, H-5’’’); 3.71-3.68 (m, 2H, H-2, H-5); 3.63-3.62 (m, 2H, H-2’, H-4’); 3.57-3.47 (m, 16H, H-2’’, $OCH_3$); 3.33 (s, 3H, $OCH_3$); 3.23 (t, 1H, *J* = 9.5 Hz, H-4); 2.87-2.84 (m, 1H, $CH_{Phth}$); 2.70 (t, 1H, *J* = 9.0 Hz, H-4’’’); 2.55-2.50 (m, 4H, $CH_{Myc}$, $CH_{2,Phth}$); 2.16-2.12 (m, 4H, H-2’’’, $CH_{3,Ac}$); 1.76-0.81 (m, 197H, H-2’’’, H-6, H-6’, H-6’’, H-6’’’, $CH_{Phth}$, $CH_{2,Phth}$, $CH_{3,Phth}$, $CH_{Myc}$, $CH_{2,Myc}$, $CH_{3,Myc}$). $^{13}$C-APT NMR (125 MHz) δ: 176.2, 176.1 ($CO_{Myc}$); 170.7 ($CO_{Ac}$); 154.6, 137.0 ($C_{q,arom}$); 129.5, 116.3 ($CH_{arom}$); 101.0 (C-1’’); 99.5 (C-1’); 99.4 (C-1’’’); 95.2 (C-1); 88.1 (C-4’’’); 86.8 ($CH_{Phth}$); 83.6 (C-3’); 82.5 (C-4); 80.7 (C-2’); 80.5 (C-2); 79.7 (C-2’’); 79.0 (C-3); 74.1 (C-3’’); 73.7 (C-4’’); 71.7 (C-4’); 70.4 ($CH_{Phth}$); 69.1 (C-5’); 68.9 (C-5); 68.0 (C-3’’’); 67.7 (C-5’’’); 66.1 (C-5’’); 61.2, 60.4, 60.0, 59.1, 58.8, 57.5 ($OCH_3$); 45.6, 45.4 ($CH_{2,Myc}$); 41.1, 38.6 ($CH_{2,Phth}$); 37.9, 37.9 ($CH_{Myc}$); 37.7, 36.8, 35.3 ($CH_{2,Myc}$); 34.9 ($CH_{Phth}$); 34.8, 32.8 ($CH_{2,Phth}$); 32.1 ($CH_{2,Myc}$); 31.9, 30.5 ($CH_{2,Phth}$); 30.2 ($CH_{Myc}$); 30.1, 29.9, 29.9, 29.8, 29.7, 29.7, 29.5 ($CH_2$); 28.2 ($CH_{Myc}$); 27.6 ($CH_{2,Phth}$); 27.3 ($CH_{Myc}$); 27.1 ($CH_{2,Myc}$); 25.7, 25.3 ($CH_{2,Phth}$); 22.8 ($CH_{2,Myc}$); 22.5 ($CH_{2,Phth}$); 21.0 ($CH_{3,Ac}$); 20.9, 20.6, 20.6, 20.5, 18.6 ($CH_{3,Myc}$); 18.3 (C-6’’’); 18.0 (C-6) 18.0 (C-6’); 16.5 (C-6’’); 14.9 ($CH_{3,Phth}$); 14.3 ($CH_{3,Myc}$); 10.3 ($CH_{3,Phth}$). IR (thin film, $cm^{-1}$): 1132, 1173, 1233, 1378, 1464, 1510, 1734, 2853, 2923, 3440. HRMS calculated for $C_{130}H_{240}O_{22}Na$ 2177.75877 $[M+Na]^+$; found 2177.75981.

##### *M. kansasii* PGL K-7

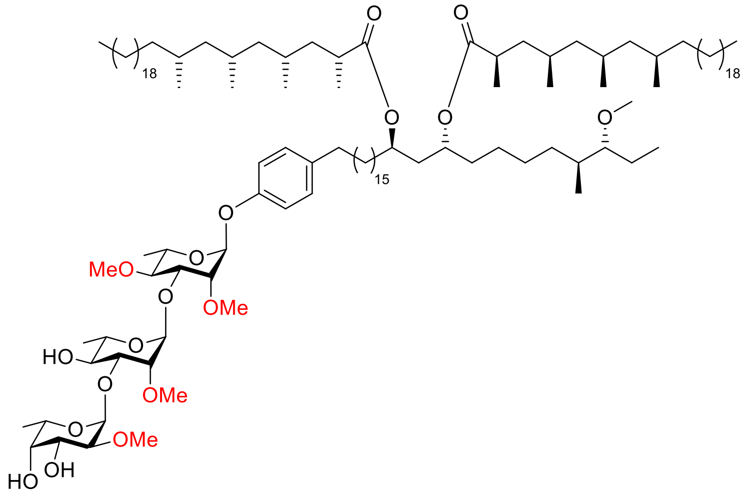

$[\alpha]_D^{25}$ = -55.6 (c = 1.0, $CHCl_3$). [1]H-NMR (400 MHz) δ: 7.09 (d, 2H, *J* = 8.4 Hz, C*H*$_{arom}$); 6.97 (d, 2H, *J* = 8.8 Hz, C*H*$_{arom}$); 5.48 (d, 1H, *J* = 1.2 Hz, H-1); 5.19 (d, 1H, *J* = 1.2 Hz, H-1'); 5.14 (d, 1H, *J* = 4.0 Hz, H-1''); 4.84 (quint, 2H, *J* = 6.4 Hz, C*H*$_{Phth}$); 4.25 (q, 1H, *J* = 6.4 Hz, H-5''); 4.12 (dd, 1H, *J*= 3.4, 9.8 Hz, H-3); 4.06 (dd, 1H, *J* = 3.2, 10.0 Hz, H-3''); 3.91-3.86 (m, 1H, H-5'); 3.83 (d, 1H, *J* = 2.8 Hz, H-4''); 3.78 (dd, 1H, *J* = 3.2, 9.6 Hz, H-3'); 3.73-3.61 (m, 3H, H-2, H-2', H-4', H-5); 3.56-3.45 (m, 13H, H-2'', OC*H*$_3$); 3.33 (s, 3H, OC*H*$_3$); 3.23 (t, 1H, *J* = 9.6 Hz, H-4); 2.88-2.84 (m, 1H, C*H*$_{Phth}$); 2.72 (bs, 1H, O*H*); 2.56-2.48 (m, 4H, C*H*$_{2,Phth}$, C*H*$_{Myc}$); 2.41 (bs, 1H, O*H*); 1.77-0.81 (m, 221H, H-6, H-6', H-6'', C*H*$_{Phth}$, C*H*$_{2,Phth}$, C*H*$_{3,Phth}$, C*H*$_{Myc}$, C*H*$_{2,Myc}$, C*H*$_{3,Myc}$). [13]C-APT NMR (101 MHz) δ: 176.1 (*C*O$_{Myc}$); 154.6, 137.0 (C$_{q,arom}$); 129.4, 116.2 (*C*H$_{arom}$); 99.8 (C-1''); 99.4 (C-1'); 95.1 (C-1); 86.8 (*C*H$_{Phth}$); 83.0 (C-3'); 82.4 (C-4); 80.6, 80.5 (C-2 and C-2'); 79.5 (C-2''); 79.3 (C-3) 72.0 (C-4''); 71.7 (C-4'); 70.4 (*C*H$_{Phth}$); 69.9 (C-3''); 69.1 (C-5); 68.9 (C-5'); 66.5 (C-5''); 61.2, 59.4, 58.7, 58.7, 57.5 (O*C*H$_3$); 45.6, 45.4 (*C*H$_{2,Myc}$); 41.1, 38.6 (*C*H$_{2,Phth}$); 37.9 (*C*H$_{Myc}$); 36.8, 35.3 (*C*H$_{2,Myc}$); 34.9 (*C*H$_{Phth}$); 34.8, 32.8 (*C*H$_{2,Phth}$); 32.1 (*C*H$_{2,Myc}$); 31.9, 30.2 (*C*H$_{2,Phth}$); 30.1 (*C*H$_{Myc}$); 29.9, 29.9, 29.8, 29.7, 29.5 (*C*H$_2$); 28.2 (*C*H$_{Myc}$); 27.6 (*C*H$_{2,Phth}$); 27.3 (*C*H$_{Myc}$); 27.1 (*C*H$_{2,Myc}$); 25.7, 25.3 (*C*H$_{2,Phth}$); 22.8 (*C*H$_{2,Myc}$); 22.5 (*C*H$_{2,Phth}$); 20.9, 20.6, 20.5, 20.5, 18.6 (*C*H$_{3,Myc}$); 18.0 (C-6) 18.0 (C-6'); 16.5 (C-6"); 14.8 (*C*H$_{3,Phth}$); 14.3 (*C*H$_{3,Myc}$); 10.3 (*C*H$_{3,Phth}$). IR (thin film, $cm^{-1}$): 1100, 1132, 1173, 1229, 1378, 1461, 1510, 1734, 2853, 2923, 3421. HRMS calculated for $C_{121}H_{227}O_{18}Na$ 1969.68761 $[M+H]^+$; found 1969.68743.

##### ***M. Haemophilum* PGL**

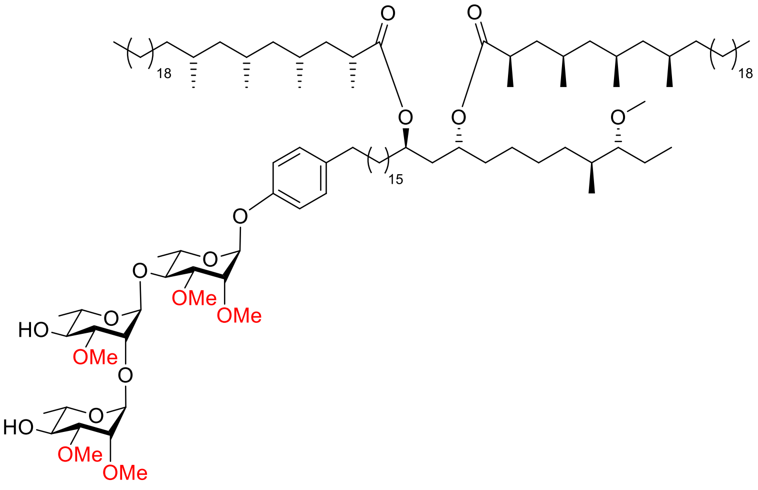

$[\alpha]_D^{25}$ = -36.1 (c = 1.0, $CHCl_3$). <u>$^1$H NMR</u> (400 MHz) δ: 7.10 (d, 2H, *J* = 8.4 Hz, C$H_{arom}$); 6.96 (dd, 2H, *J* = 2.0, 6.4 Hz, C$H_{arom}$); 5.50 (d, 1H, *J* = 1.6 Hz, H-1); 5.26 (d, 1H, *J* = 1.6 Hz, H-1'); 5.15 (d, 1H, *J* = 1.6 Hz, H-1"); 4.84 (quint, 2H, *J* = 6.4 Hz, C$H_{Phth}$); 4.16 (dd, 1H, *J* = 2.0, 2.4 Hz, H-2'); 3.79-3.70 (m, 6H, H-2, H-3, H-4, H-5, H-5', H-5''); 3.68 (dd, 1H, *J* = 1.8, 3.0 Hz, H-2"); 3.58-3.41 (m, 18H, H-3", H-4', H-4", OC$H_3$); 3.37 (dd, 1H, *J* = 2.6, 9.4 Hz, H-3'); 3.33 (s, 3H, OC$H_3$); 2.88-2.83 (m, 1H, C$H_{Phth}$); 2.57-2.52 (m, 4H, C$H_{2,Phth}$, C$H_{Myc}$); 2.29 (bs, 2H, 4'-O*H,* 4"-OH); 1.77-0.81 (m, 218H, C$H_{Phth}$, C$H_{2,Phth}$, C$H_{3,Phth}$, C$H_{Myc,}$ C$H_{2,Myc}$, C$H_{3,Myc}$, H-6, H-6', H-6"). <u>$^{13}$C-APT NMR</u> (100 MHz) δ: 176.2, 176.1 (*C*$O_{Myc}$); 154.6, 137.0 ($C_{q,arom}$); 129.5, 116.3 (*C*$H_{arom}$); 101.0 (C-1'); 98.3 (C-1"); 96.1 (C-1); 86.8 (*C*$H_{Phth}$); 82.1 (C-3 and C-3'); 80.8 (C-3"); 77.9 (C-4); 76.3 (C-2); 76.1 (C-2''); 71.8, 71.7 (C-4'and C-4"); 71.2 (C-2') 70.4 (*C*$H_{Phth}$); 69.2 (C-5'); 68.8 (C-5"); 68.0 (C-5); 59.6, 59.2, 57.5, 57.5, 57.1, 57.1 (O*C*$H_3$); 45.6, 45.4 (*C*$H_{2,Myc}$); 41.1, 38.6 (*C*$H_{2,Phth}$); 37.9, 37.9 (*C*$H_{Myc}$); 36.8, 35.3 (*C*$H_{2,Myc}$); 34.9 (*C*$H_{Phth}$); 34.8, 32.8 (*C*$H_{2,Phth}$); 32.1 (*C*$H_{2,Myc}$); 31.9, 30.2 (*C*$H_{2,Phth}$); 30.1 (*C*$H_{Myc}$); 29.9, 29.9, 29.8, 29.7, 29.5 (*C*$H_2$); 28.2 (*C*$H_{Myc}$); 27.6 (*C*$H_{2,Phth}$); 27.3 (*C*$H_{Myc}$); 27.1 (*C*$H_{2,Myc}$); 25.7, 25.3 (*C*$H_{2,Phth}$); 22.8 (*C*$H_{2,Myc}$); 22.5 (*C*$H_{2,Phth}$); 20.9, 20.6, 20.6, 20.5 (*C*$H_{3,Myc}$); 18.6 (*C*$H_{3,Myc}$); 18.5, (C-6'); 17.9, 17.7 (C-6 and C-6''); 14.8 (*C*$H_{3,Phth}$); 14.3 (*C*$H_{3,Myc}$); 10.3 (*C*$H_{3,Phth}$). <u>IR</u> (thin film, $cm^{-1}$): 1009, 1075, 1082, 1093, 1106, 1122, 1262, 1378, 1457, 1464, 1510, 1736, 2853, 2923, 3430. <u>HRMS</u> calculated for $C_{122}H_{229}O_{18}$ 1983.70326 $[M+H]^+$; found 1983.70441.

##### ***M. bovis*** **PGL (Mycoside B)**

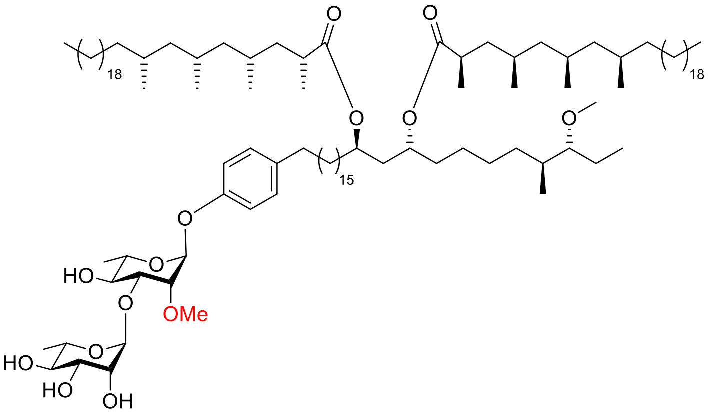


$[\alpha]_D^{25}$ = -23.0 (c = 0.5, $CHCl_3$). $^1$<u>H-NMR</u> (400 MHz) δ: 7.10 (d, 2H, *J* = 8.8 Hz, $CH_{arom}$); 6.98 (d, 2H, *J* = 8.4 Hz, $CH_{arom}$); 5.49 (s, 1H, H-1); 5.10 (s, 1H, H-1'); 4.84 (quint, 2H, *J* = 6.4 Hz, $CH_{Myc}$); 4.06-4.01 (m, 2H, H-2', H-3); 3.86-3.79 (m, 2H, H-3', H-5'); 3.76-3.72 (m, 2H, H-2, H-5); 3.59 (t, 1H, *J* = 9.6 Hz, H-4); 3.51-3.42 (m, 4H, H-4', $OCH_3$); 3.33 (s, 3H, $OCH_3$); 2.89-2.85 (m, 1H, $CH_{Phth}$); 2.56-2.50 (m, 4H, $CH_{2,Phth}$, $CH_{Myc}$); 2.27 (bs, 1H, O*H*); 1.77-0.81 (m, 195H, H-6, H-6', $CH_{Phth}$, $CH_{2,Phth}$, $CH_{3,Phth}$, $CH_{Myc,}$ $CH_{2,Myc}$, $CH_{3,Myc}$). $^{13}$<u>C-APT NMR</u> (101 MHz) δ: 176.3 ($CO_{Myc}$); 154.6, 137.0 ($C_{q,arom}$); 129.4, 116.3 ($CH_{arom}$); 102.5 (C-1'); 95.2 (C-1); 86.8 ($CH_{Phth}$); 80.3 (C-2); 79.2 (C-3); 73.0 (C-4'); 71.9 (C-4); 71.4 (C-3'); 70.7 (C-2'); 70.5 ($CH_{Phth}$); 69.3 (C-5); 68.9 (C-5'); 58.9, 57.5 ($OCH_3$); 45.6, 45.4 ($CH_{2,Myc}$); 41.1, 38.5 ($CH_{2,Phth}$); 37.9 ($CH_{Myc}$); 36.7, 35.3 ($CH_{2,Myc}$); 34.9 ($CH_{Phth}$); 34.8, 32.7 ($CH_{2,Phth}$); 32.0 ($CH_{2,Myc}$); 31.8, 30.2 ($CH_{2,Phth}$); 30.0 ($CH_{Myc}$); 29.9, 29.8, 29.8, 29.7, 29.5 ($CH_2$); 28.1 ($CH_{Myc}$); 27.6 ($CH_{2,Phth}$); 27.3 ($CH_{Myc}$); 27.1 ($CH_{2,Myc}$); 25.7, 25.3 ($CH_{2,Phth}$); 22.8 ($CH_{2,Myc}$); 22.4 ($CH_{2,Phth}$); 20.8, 20.6, 20.5, 18.5 ($CH_{3,Myc}$); 17.8 (C-6 and C-6'); 14.7 ($CH_{3,Phth}$); 14.2 ($CH_{3,Myc}$); 10.2 ($CH_{3,Phth}$). <u>IR</u> (thin film, $cm^{-1}$): 1098, 1130, 1173, 1229, 1261, 1378, 1457, 1511, 1736, 2853, 2923, 3396. <u>HRMS</u> calculated for $C_{112}H_{211}O_{14}$ 1781.58275 $[M+H]^+$; found 1781.58272.

#### Synthesis of PGL analogues

The MTb PGL-I C18 was assembled using the general synthesis strategy, using octadec-1-yne in the Sonogashira cross-coupling reaction.

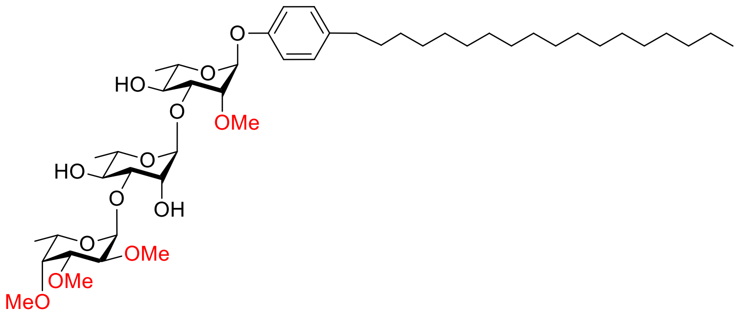


$[\alpha]_D^{25}$ = -91.8 (c = 1.0, $CHCl_3$). $^1$H-NMR (400 MHz) δ: 7.10 (d, 2H, *J* = 8.4 Hz, $CH_{arom}$); 6.99 (dd, 2H, *J* = 2.2, 6.6 Hz, $CH_{arom}$); 5.51 (d, 1H, *J* = 1.6 Hz, H-1); 5.15-5.14 (m, 2H, H-1', H-1"); 4.11 (s, 1H, H-2'); 4.09-4.03 (m, 2H, H-3, H-5"); 3.97-3.90 (m, 1H, H-5'); 3.81-3.74 (m, 3H, H-2, H-3', H-5); 3.70-3.57 (m, 10H, H-2", H-3", H-4, H-4', $OCH_3$); 3.52 (s, 3H, $OCH_3$); 3.49 (s, 3H, $OCH_3$); 3.48 (d, 1H, *J* = 1.6 Hz, H-4"); 2.55 (t, 2H, *J* = 7.6 Hz, $CH_2$); 2.35 (bs, 1H, O*H*); 2.21 (bs, 1H, O*H*); 1.55 (quint, 2H, *J* = 7.6 Hz, $CH_2$); 1.36 (d, 3H, *J* = 6.4 Hz, H-6'); 1.34-1.25 (m, 38H, H-6, H-6", $CH_2$); 0.88 (t, 3H, *J* = 7.4 Hz, $CH_3$). $^{13}$C-APT NMR (101 MHz) δ: 154.7, 137.0 ($C_{q,arom}$); 129.5, 116.3 ($CH_{arom}$); 102.3 (C-1"); 100.9 (C-1'); 95.0 (C-1); 83.3 (C-3'); 81.1 (C-3"); 80.2 (C-2); 80.0 (C-3); 79.1 (C-4"); 78.9 (C-2"); 71.9 (C-4'); 71.7 (C-4); 71.3 (C-2'); 69.2 (C-5); 68.8 (C-5'); 67.6 (C-5"); 62.1, 60.4, 58.7, 57.9 ($OCH_3$); 35.3, 32.1, 29.8, 29.8, 29.7, 29.5, 29.5, 22.8 ($CH_2$); 18.0, 17.9 (C-6 and C-6'); 16.8 (C-6"); 14.3 ($CH_3$). IR (thin film, $cm^{-1}$): 1043, 1089, 1129, 1195, 1229, 1510, 2852, 2923, 3433. HRMS calculated for $C_{46}H_{84}O_{13}N$ 858.59372 $[M+NH_4]^+$; found 858.59328.

The analogues lacking the phthiocerol dimycocerosate lipids were assembled by hydrogenation of the parent iodoaryl glycans.

**Phenyl rhamnose**

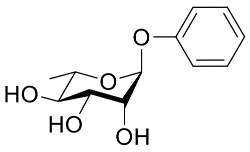


The starting material (100 mg, 0.27 mmol, 1.0 eq) and $NH_4OAc$ (69 mg, 0.89 mmol, 3.3 eq) were dissolved in EtOH (27 mL, 0.01 M). The solution was sparged with Ar for 30 min. Pd/C (10 wt%, catalytic amount) was added and the resulting mixture was sparged with Ar for 30 min followed by sparging with $H_2$ for 30 min. The reaction mixture was stirred for 3 days under a $H_2$ atmosphere. Progress of the reaction was monitored with TLC (9:1 DCM/MeOH). After completion, the reaction mixture was filtered over a Whatman filter, concentrated *in vacuo* and co-evaporated with MeOH (3x). Flash column chromatography (2% → 10% MeOH in EtOAc) afforded the title compound as a slightly yellow oil. The product was further purified through an acetylation/deacetylation sequence. The crude was dissolved in pyridine (2 mL, 0.14 M) under a $N_2$ atmosphere. $Ac_2O$ (0.10 mL, 1.1 mmol, 4.1 eq) was added to the reaction mixture, which was stirred for 16 h. Progress of the reaction was monitored with TLC (9:1 DCM/MeOH). After completion, the reaction mixture was concentrated *in vacuo*. The resulting mixture was redissolved in EtOAc and washed with 1M HCl, sat. aq. $NaHCO_3$, sat. aq. $N_2S_2O_3$ and brine. The organic layer was dried over $MgSO_4$, filtered and concentrated *in vacuo*. Purification of the crude using flash column chromatography (10% → 40% EtOAc in pentane) afforded the corresponding tri-acetate intermediate, which was immediately used in the next step. The tri-acetate intermediate was dissolved in MeOH (2 mL, 0.14 M). NaOMe (25 wt% in MeOH, 0.03 mL, 0.135 mmol, 0.5 eq) was added to the reaction mixture which was stirred for 2 days. Progress of the reaction was monitored with TLC. After completion, the reaction mixture was neutralized with Amberlite $H^+$, filtered and concentrated *in vacuo*. Purification of the crude using flash column chromatography (2% → 10% MeOH in EtOAc) afforded the title compound (28 mg, 0.117 mmol, 43% yield over 3 steps) as a colorless oil.

$[\alpha]_D^{25}$ = -101 (c = 1.0, MeOH). (c = 1.0, MeOH). $^1$H NMR (400 MHz, MeOD): δ 7.27 (dd, *J* = 7.3, 8.7 Hz, 2H, C$H_{arom}$), 7.08 – 7.03 (m, 2H, C$H_{arom}$), 6.99 (tt, *J* = 1.1, 7.4 Hz, 1H, C$H_{arom}$), 5.42 (d, *J* = 1.8 Hz, 1H, H-1), 4.01 (dd, *J* = 1.8, 3.4 Hz, 1H, H-2), 3.86 (dd, *J* = 9.5, 3.4 Hz, 1H, H-3), 3.65 (dq, *J* = 9.5, 6.2 Hz, 1H, H-5), 3.47 (t, *J* = 9.5 Hz, 1H, H-4), 1.22 (d, *J* = 6.2 Hz, 3H, H-6). $^{13}$C NMR (101 MHz, MeOD) δ 157.8 ($C_{q,arom}$), 130.5, 123.2, 117.5 ($CH_{arom}$), 99.8 (C-1), 73.8 (C-4), 72.2 (C-3), 72.1 (C-2), 70.6 (C-5), 18.0 (C-6). HRMS calculated for $C_{12}H_{16}O_5Na$ 263.08899 $[M+Na]^+$; found 263.08872

**Phenyl di-rhamnose**

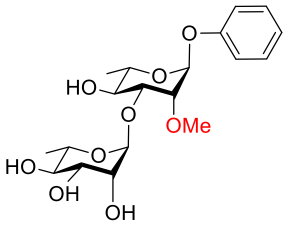


The title compound was prepared according to the general two-step deiodination and hydrogenation procedure. Colorless oil, (24 mg, 55 µmol, 79% yield over 2 steps).

$[\alpha]_D^{25}$ = -93° (c = 1.0, MeOH). $^1$H NMR (400 MHz, MeOD): δ 7.29 (dd, *J* = 8.7, 7.4 Hz, 2H, $CH_{arom}$), 7.07 (dt, *J* = 7.9, 1.0 Hz, 2H, $CH_{arom}$), 7.01 (tt, *J* = 7.4, 1.1 Hz, 1H, $CH_{arom}$), 5.56 (d, *J* = 2.0 Hz, 1H, H-1), 5.15 (d, *J* = 1.7 Hz, 1H, H-1‘), 4.09 (dd, *J* = 9.6, 3.2 Hz, 1H H-3), 3.82 – 3.74 (m, 2H, H-3‘, H-5‘), 3.72 (dd, *J* = 3.3, 1.9 Hz, 1H, H-2), 3.66 – 3.60 (m, 1H, H-5), 3.55 (m, 7H, H-2‘, OMe 2x), 3.49 (s, 3H, OMe), 3.42 – 3.34 (m, 1H, H-4‘), 3.23 (t, *J* = 9.5 Hz, 1H, H-4), 1.30 (d, *J* = 6.2 Hz, 3H, H-6‘), 1.22 (d, *J* = 6.3 Hz, 3H, H-6). $^{13}$C NMR (101 MHz, MeOD): δ 157.7 ($C_{q,arom}$), 130.6, 123.4, 117.6 ($CH_{arom}$), 100.4 (C-1‘), 96.4 (C-1), 83.8 (C-4), 82.6 (C-2‘), 81.5 (C-2), 79.5 (C-3), 74.2 (C-4‘), 72.1 (C-3‘), 70.5 (C-5‘), 70.1 (C-5), 61.5, 59.4, 59.3 (OMe), 18.2, 18.1 (C-6, C-6‘). HRMS calculated for $C_{21}H_{32}O_9Na$ 451.19385 [M+Na]+; found 451.19422.

#### MTb PGL-1 phenyl trisaccharide

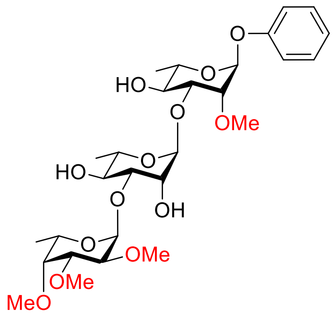


The title compound was prepared according to the general two-step deiodination and hydrogenation procedure. Colorless oil, (21 mg, 36 µmol, 92%).

$[\alpha]_D^{25}$ = -123.4 (c = 1.0, $CHCl_3$). [1]H-NMR (400 MHz) δ: 7.29 (t, 2H, *J* = 7.6 Hz, $CH_{arom}$); 7.08 (dd, 2H, *J* = 0.5, 8.6 Hz, $CH_{arom}$); 7.03 (t, 2H, *J* = 7.8 Hz, $CH_{arom}$); 5.56 (d, 1H, *J* = 1.2 Hz, H-1); 5.15-5.14 (m, 2H, H-1', H-1"); 4.11 (s, 1H, H-2'); 4.09-4.03 (m, 3H, H-3, H-5", O*H*); 3.96-3.90 (m, 1H, H-5'); 3.82-3.71 (m, 3H, H-2, H-3', H-5); 3.69-3.58 (m, 10H, H-2", H-3", H-4, H-4', $OCH_3$); 3.55-3.45 (m, 7H, H-4'', $OCH_3$); 2.45 (d, 1H, *J* = 3.2 Hz, O*H*); 2.29 (d, 1H, *J* = 3.2 Hz, O*H*); 1.37 (d, 3H, *J* = 6.0 Hz, H-6'); 1.30-1.25 (m, 6H, H-6, H-6''). [13]C-APT NMR (101 MHz) δ: 156.6, ($C_{q,arom}$); 129.7, 122.4 116.4 ($CH_{arom}$); 102.3 (C-1"); 100.9 (C-1'); 94.8 (C-1); 83.2 (C-3'); 81.1 (C-3"); 80.1 (C-2); 79.9 (C-3); 79.1 (C-4"); 78.9 (C-2"); 71.9 (C-4'); 71.7 (C-4); 71.3 (C-2'); 69.3 (C-5); 68.8 (C-5'); 67.6 (C-5"); 62.1, 60.4, 58.8, 57.9 ($OCH_3$); 18.0 (C-6') 17.9 (C-6); 16.8 (C-6"). IR (thin film, $cm^{-1}$): 1042, 1089, 1128, 1365, 1495, 2853, 2925, 3419. HRMS calculated for $C_{28}H_{44}O_{13}Na$ 611.26741 $[M+Na]^+$; found 611.26758.

#### M. leprae PGL-I phenyl trisaccharide

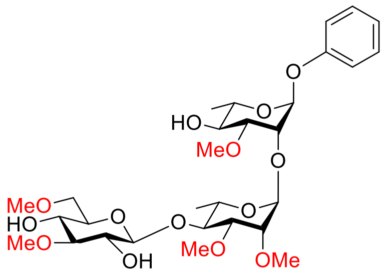


The title compound was prepared according to the general two-step deiodination and hydrogenation procedure. Colorless oil, (28 mg, 45 μmol, 94%).

$[\alpha]_D^{25}$ = -64.3 (c = 1.0, $CHCl_3$). $^1$H-NMR (400 MHz) δ: 7.29-7.27 (m, 2H, $CH_{arom}$); 7.06-7.01 (m, 3H, $CH_{arom}$); 5.48 (d, 1H, *J* = 2.0 Hz, H-1); 5.11 (d, 1H, *J* = 1.6 Hz, H-1'); 4.41 (d, 1H, *J* = 8.0 Hz, H-1"); 4.24 (dd, 1H, *J* = 2.4, 4.8 Hz, H-2); 3.89 (d, 1H, *J* = 1.2 Hz, 2"-O*H*); 3.78-3.71 (m, 3H, H-2', H-5, H-5'); 3.69-3.49 (m, 19H, H-3, H-3', H-4, H-4', H-4'', H-6", $OCH_3$); 3.45-3.38 (m, 5H, H-2", H-5", $OCH_3$); 3.17 (t, 1H, *J* = 9.0 Hz, H-3"); 2.94 (d, *J* = 1.6 Hz, 1H, O*H*); 2.45 (d, *J* = 1.6 Hz, 1H, O*H*); 1.34 (d, 3H, *J* = 6.0 Hz, H-6'); 1.27 (d, 3H, *J* = 6.0 Hz, H-6'). $^{13}$C-APT NMR (101 MHz) δ: 156.2 ($C_{q,arom}$); 129.7, 122.4, 116.3 ($CH_{arom}$); 105.7 (C-1"); 98.6 (C-1'); 97.2 (C-1); 85.6 (C-3''); 81.7 (C-4'); 81.5 (C-3); 80.3 (C-3'); 75.8 (C-2'); 75.1 (C-2"); 74.2 (C-5"); 72.9 (C-6"); 72.2 (C-2); 71.9 (C-4); 71.2 (C-4"); 69.1 (C-5); 68.4 (C-5'); 60.7, 59.7, 59.2, 57.8, 56.7 ($OCH_3$); 17.9 (C-6); 17.7 (C-6'). IR (thin film, $cm^{-1}$): 1009, 1069, 1120, 1228, 1387, 1494, 2854, 2928, 3429. HRMS calculated for $C_{29}H_{46}O_{14}Na$ 641.27798 $[M+H]^+$; found 641.27776.

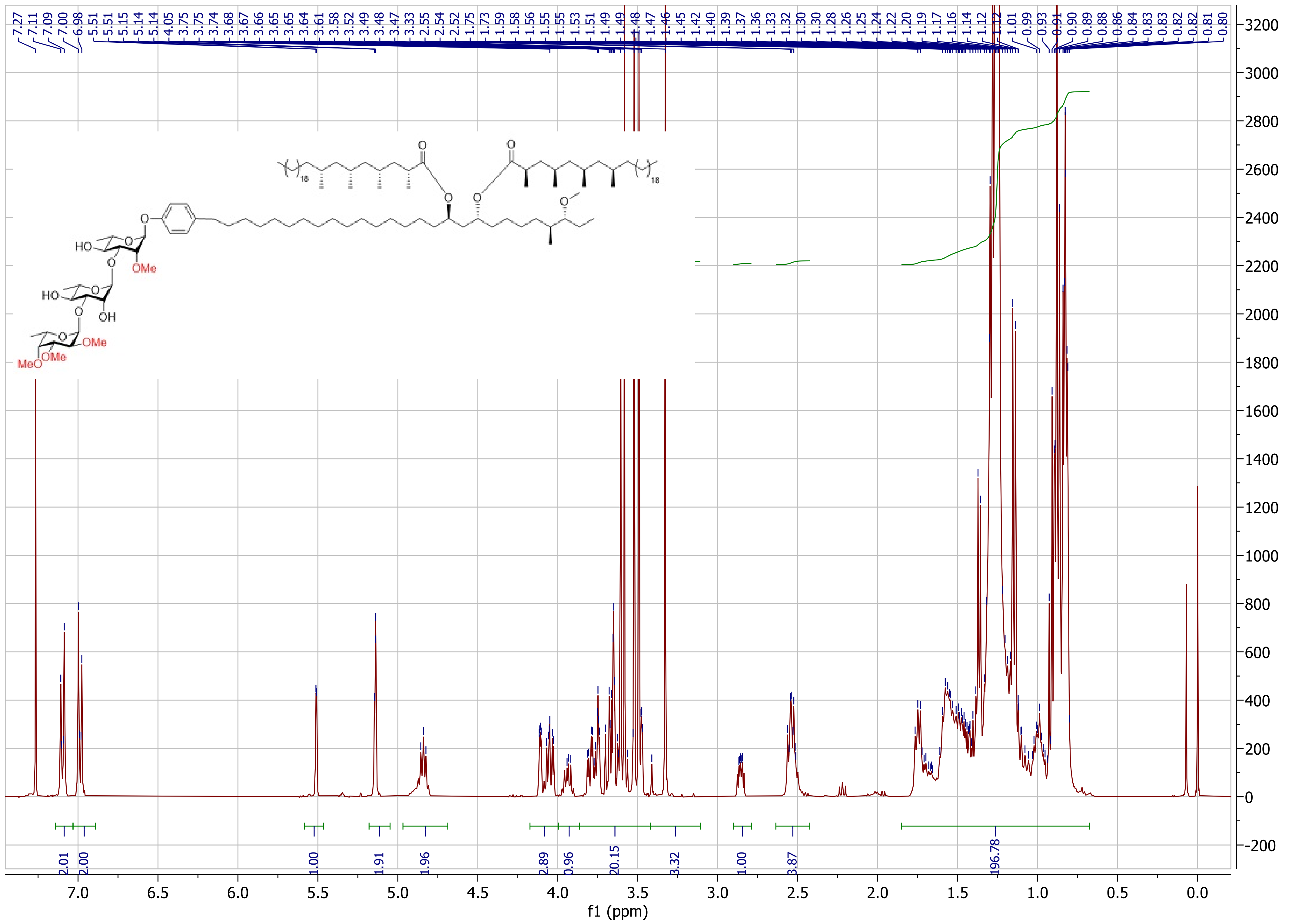

7.27 7.11 7.09 7.00 6.98 5.51 5.51 5.15 5.14 5.14 4.05 3.75 3.75 3.74 3.68 3.67 3.66 3.65 3.65 3.64 3.61 3.58 3.52 3.49 3.48 3.47 3.33 2.55 2.54 2.52 1.75 1.73 1.59 1.58 1.56 1.55 1.55 1.53 1.51 1.49 1.49 1.48 1.47 1.46 1.45 1.42 1.40 1.39 1.37 1.36 1.33 1.32 1.30 1.30 1.28 1.26 1.25 1.24 1.22 1.20 1.19 1.17 1.16 1.14 1.12 1.12 1.01 0.99 0.93 0.91 0.90 0.89 0.88 0.86 0.84 0.83 0.83 0.82 0.82 0.81 0.80
2.01 2.00 1.00 1.91 1.96 2.89 0.96 20.15 3.32 1.00 3.87 196.78
f1 (ppm)
HO
OMe
OH
MeO
18

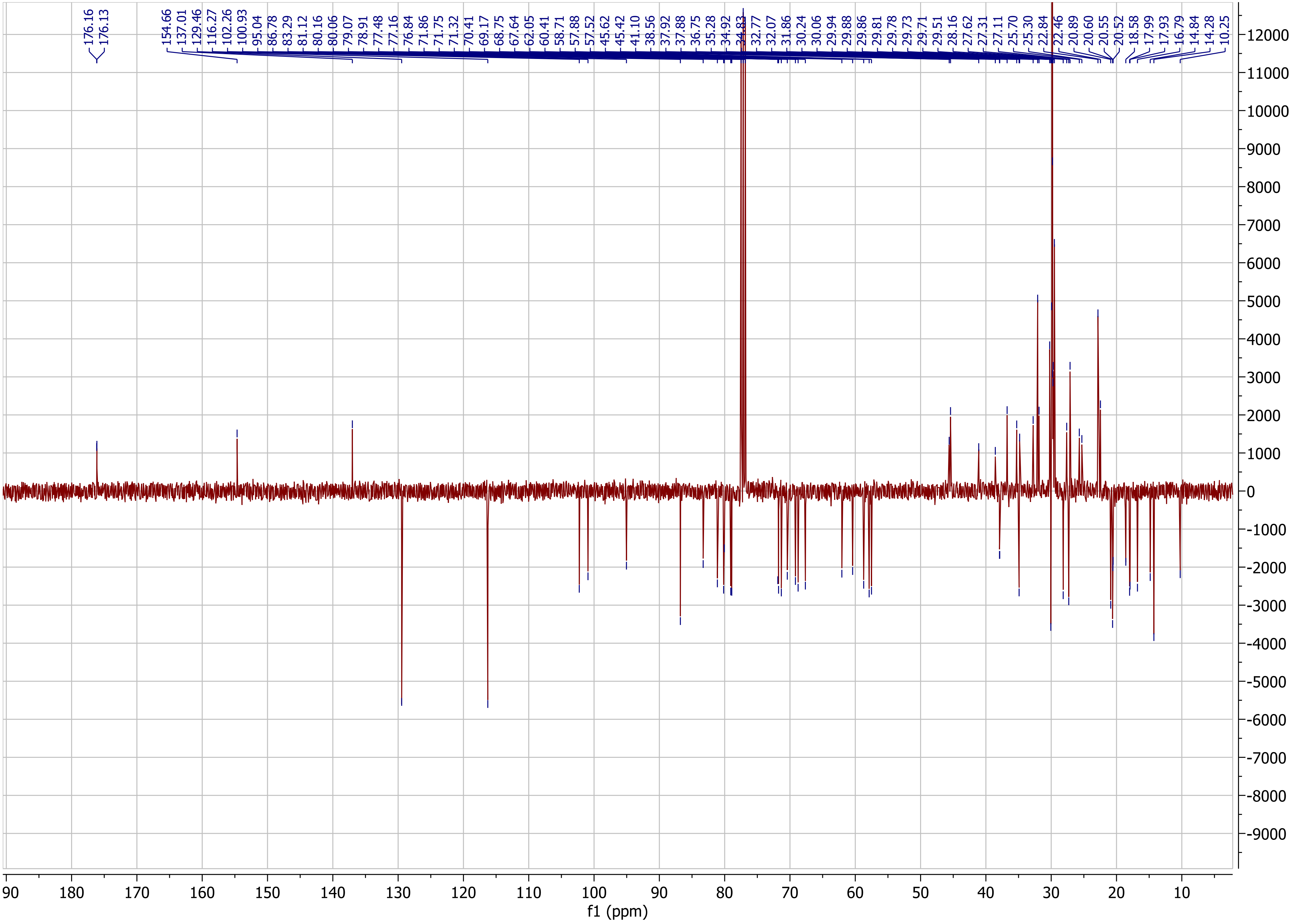

176.16
176.13
154.66
137.01
129.46
116.27
102.26
100.93
95.04
86.78
83.29
81.12
80.16
80.06
79.07
78.91
77.48
77.16
76.84
71.86
71.75
71.32
70.41
69.17
68.75
67.64
62.05
60.41
58.71
57.88
57.52
45.62
45.42
41.10
38.56
37.92
37.88
36.75
35.28
34.92
34.83
32.77
32.07
31.86
30.24
30.06
29.94
29.88
29.86
29.81
29.78
29.73
29.71
29.51
28.16
27.62
27.31
27.11
25.70
25.30
22.84
22.46
20.89
20.60
20.55
20.52
18.58
17.99
17.93
16.79
14.84
14.28
10.25
f1 (ppm)

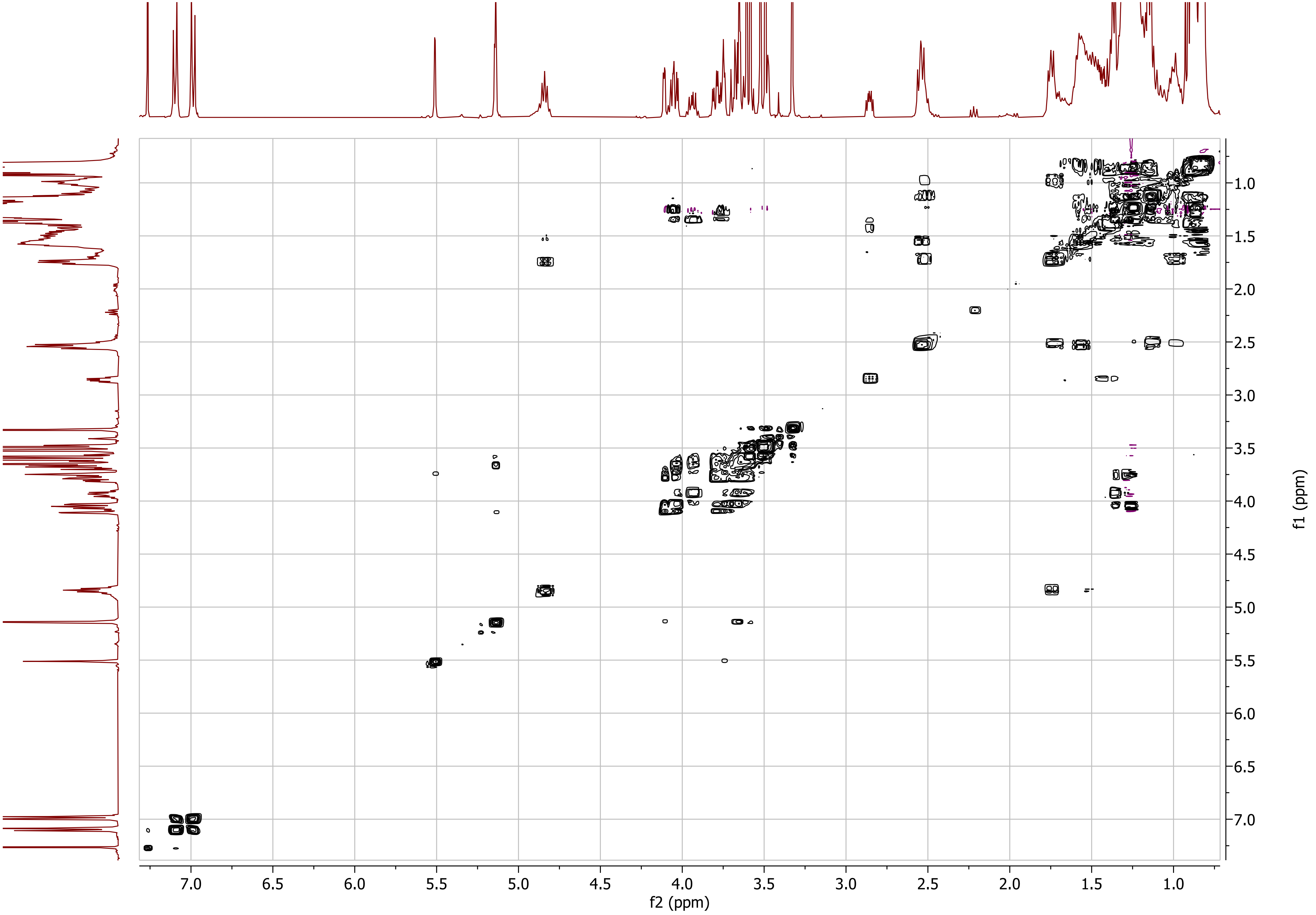

7.0
6.5
6.0
5.5
5.0
4.5
4.0
3.5
3.0
2.5
2.0
1.5
1.0
f2 (ppm)
f1 (ppm)

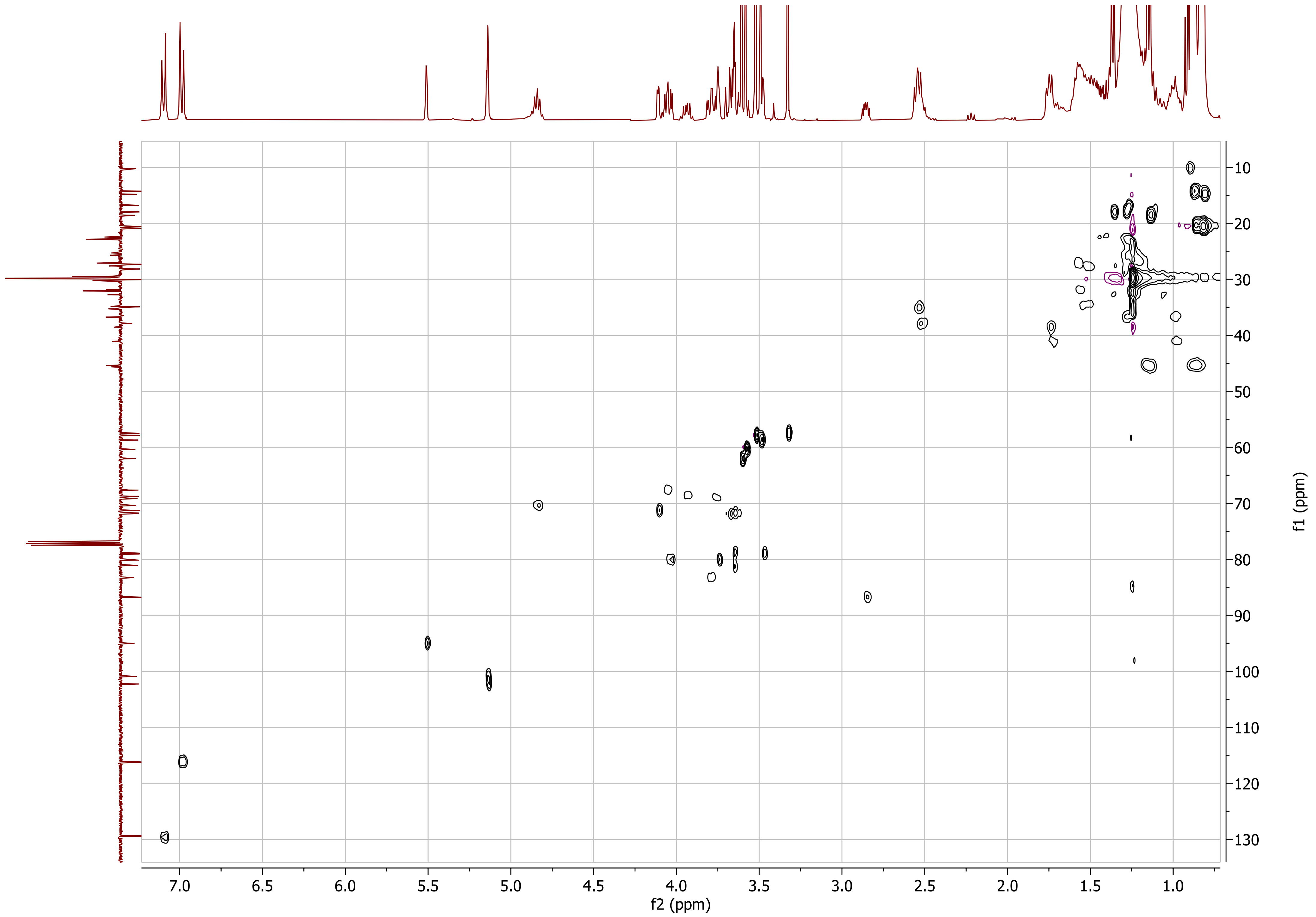

f2 (ppm)
f1 (ppm)

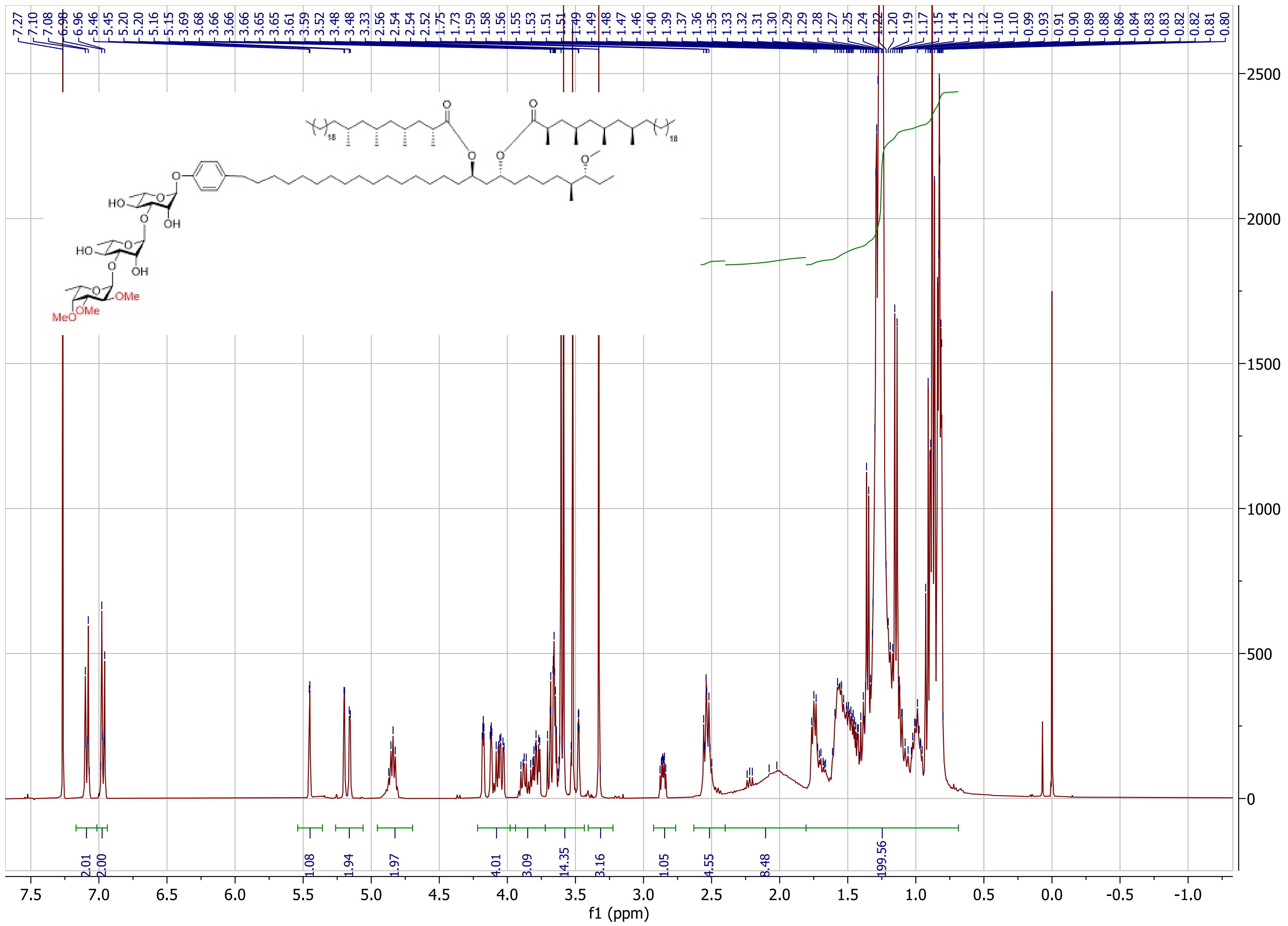
7.27 7.10 7.08 6.98 6.96 5.46 5.45 5.20 5.20 5.16 5.15 3.69 3.68 3.66 3.66 3.66 3.65 3.65 3.61 3.59 3.52 3.48 3.48 3.33 2.56 2.54 2.54 2.52 1.75 1.73 1.59 1.58 1.56 1.55 1.53 1.51 1.51 1.49 1.49 1.48 1.47 1.46 1.40 1.39 1.37 1.36 1.35 1.33 1.32 1.31 1.30 1.29 1.29 1.28 1.27 1.25 1.24 1.22 1.20 1.19 1.17 1.15 1.14 1.12 1.12 1.10 1.10 0.99 0.93 0.91 0.90 0.89 0.88 0.86 0.84 0.83 0.83 0.82 0.82 0.81 0.80
HO
OH
HO
OH
OMe
MeO
OMe
18
18
2.01
2.00
1.08
1.94
1.97
4.01
3.09
14.35
3.16
1.05
4.55
8.48
199.56
2500
2000
1500
1000
500
0
7.5 7.0 6.5 6.0 5.5 5.0 4.5 4.0 3.5 3.0 2.5 2.0 1.5 1.0 0.5 0.0 -0.5 -1.0
f1 (ppm)

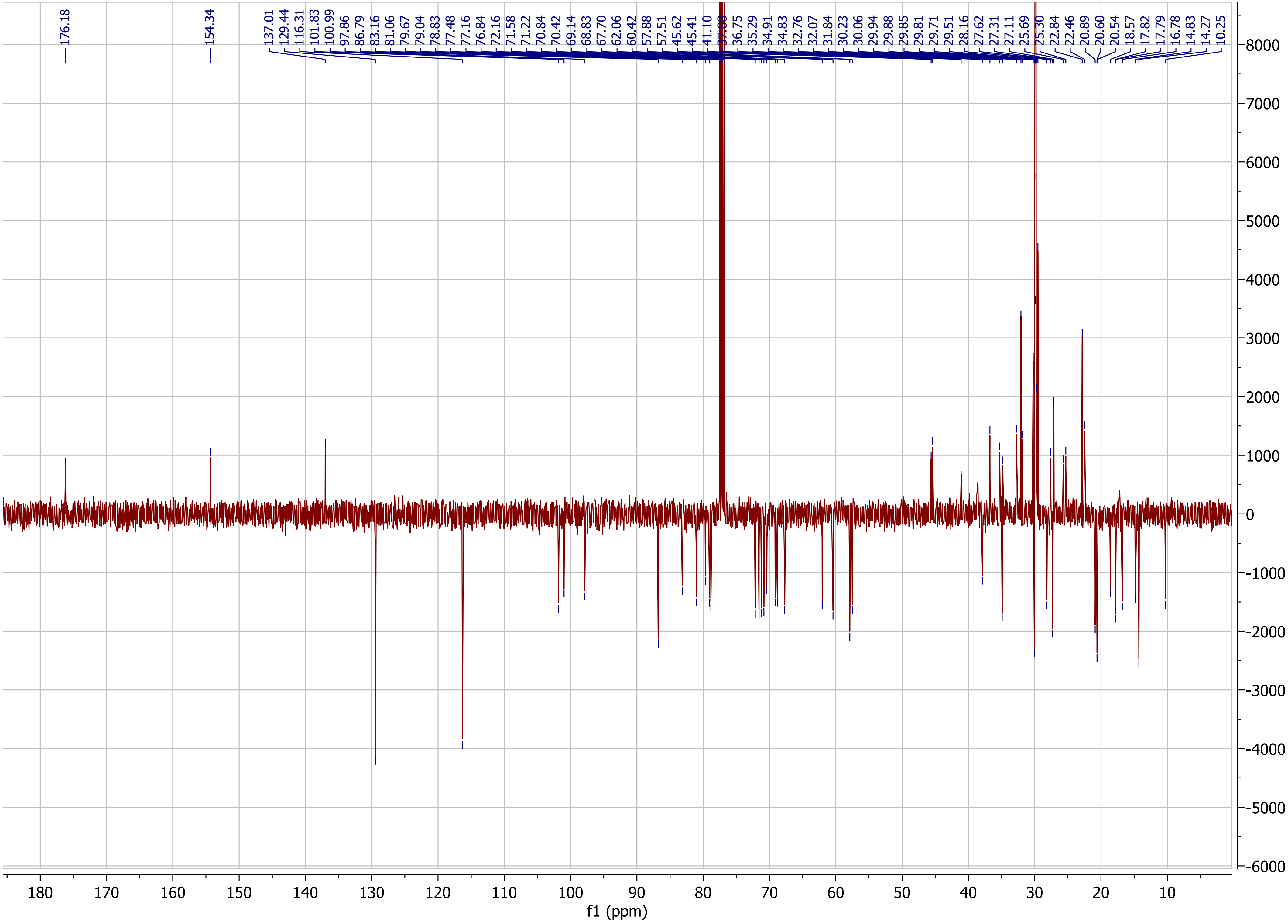
176.18
154.34
137.01
129.44
116.31
101.83
100.99
97.86
86.79
83.16
81.06
79.67
79.04
78.83
77.48
77.16
76.84
72.16
71.58
71.22
70.84
70.42
69.14
68.83
67.70
62.06
60.42
57.88
57.51
45.62
45.41
41.10
37.88
36.75
35.29
34.91
34.83
32.76
32.07
31.84
30.23
30.06
29.94
29.88
29.85
29.81
29.71
29.51
28.16
27.62
27.31
27.11
25.69
25.30
22.84
22.46
20.89
20.60
20.54
18.57
17.82
17.79
16.78
14.83
14.27
10.25
f1 (ppm)

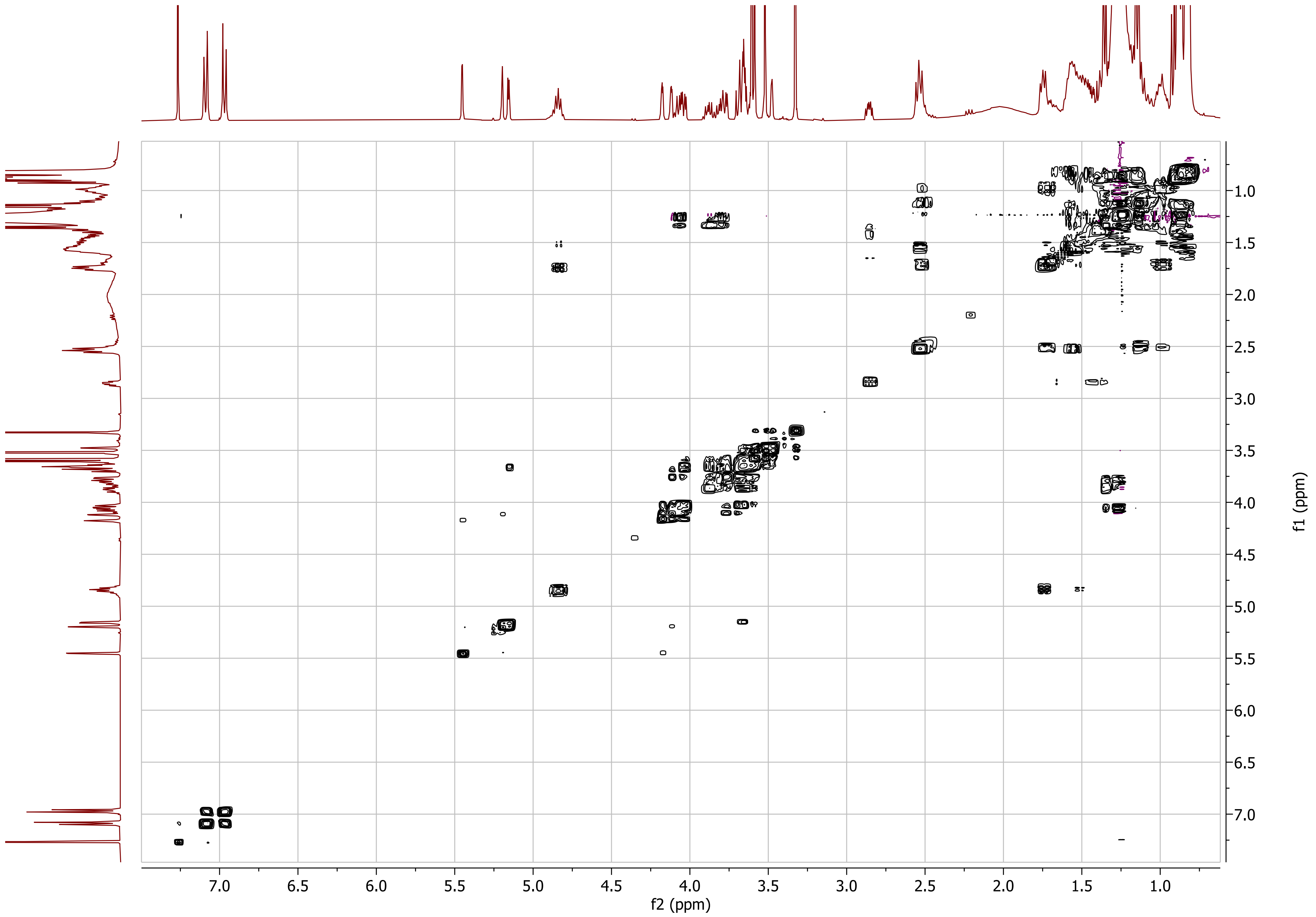

f2 (ppm)
f1 (ppm)
7.0
6.5
6.0
5.5
5.0
4.5
4.0
3.5
3.0
2.5
2.0
1.5
1.0

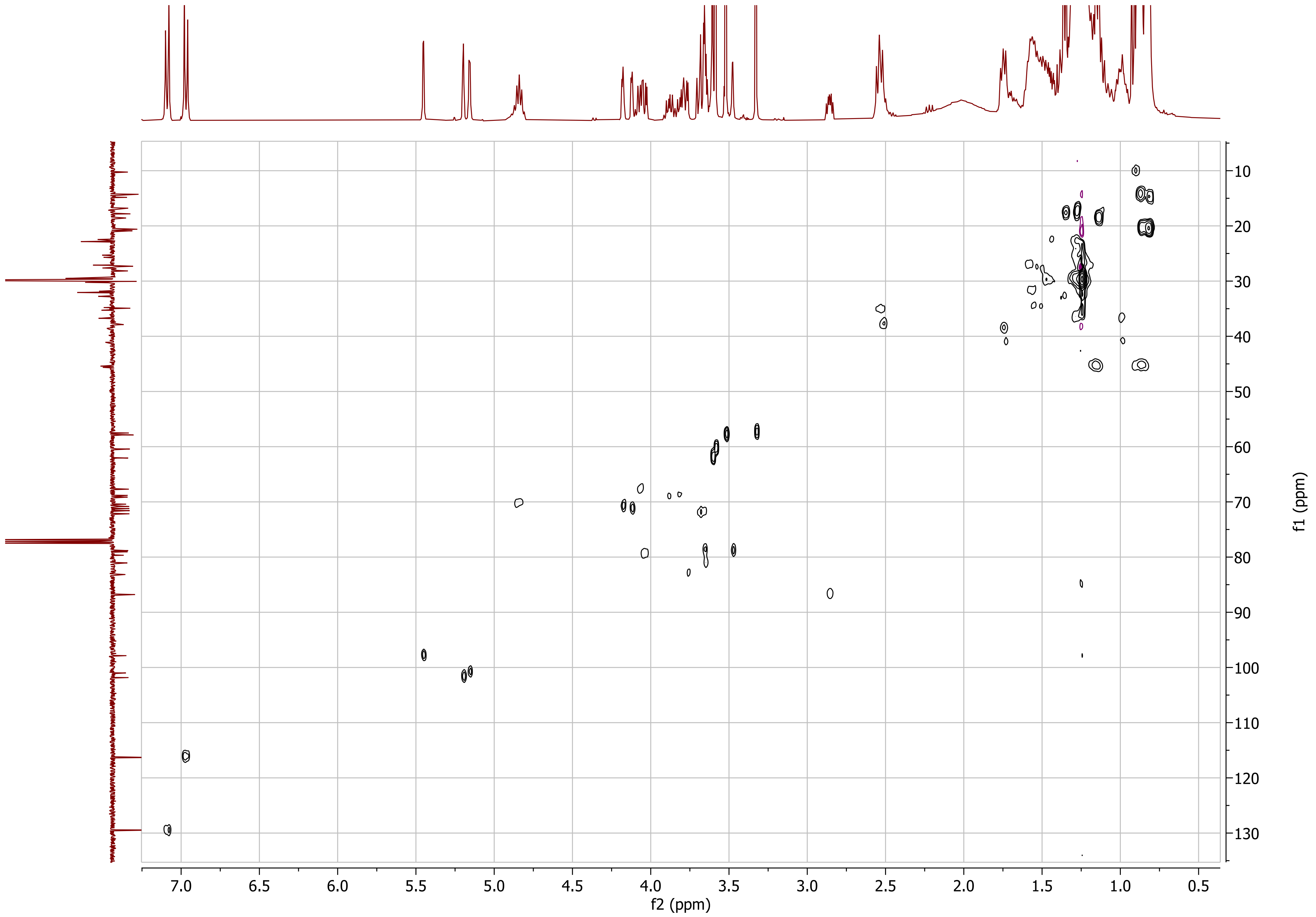

f2 (ppm)
f1 (ppm)

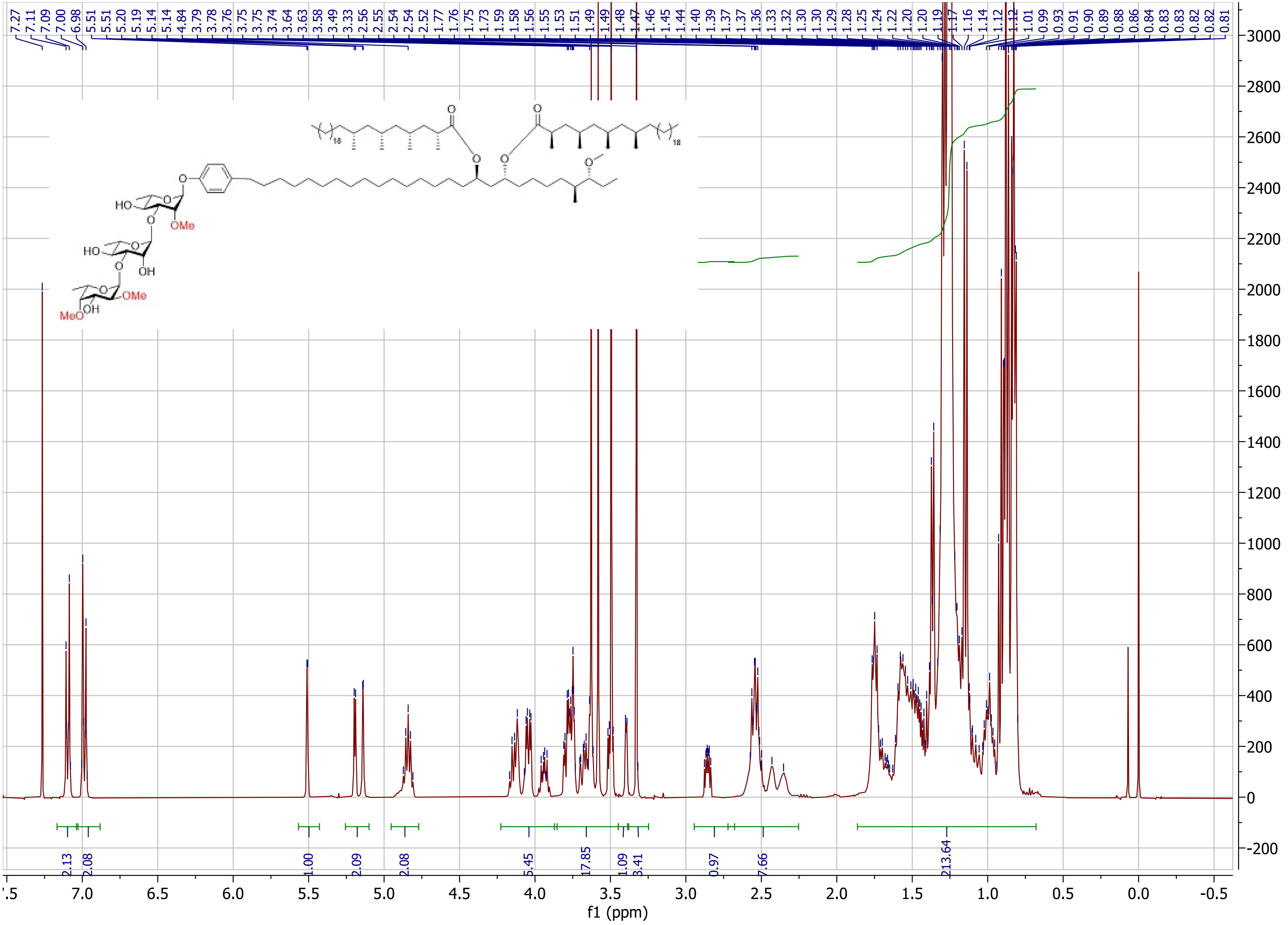

7.27 7.11 7.09 7.00 6.98 5.51 5.51 5.20 5.19 5.14 5.14 4.84 3.79 3.78 3.76 3.75 3.75 3.74 3.64 3.63 3.58 3.49 3.33 2.56 2.55 2.54 2.54 2.52 1.77 1.76 1.75 1.73 1.59 1.58 1.56 1.55 1.53 1.51 1.49 1.49 1.48 1.47 1.46 1.45 1.44 1.40 1.39 1.37 1.37 1.36 1.33 1.32 1.30 1.30 1.29 1.28 1.25 1.24 1.22 1.20 1.20 1.19 1.17 1.16 1.14 1.12 1.12 1.01 0.99 0.93 0.91 0.90 0.89 0.88 0.86 0.84 0.83 0.83 0.82 0.81
HO
OMe
HO
OH
OMe
MeO
OH
18
18
2.13
2.08
1.00
2.09
2.08
5.45
17.85
1.09
3.41
0.97
7.66
213.64
f1 (ppm)

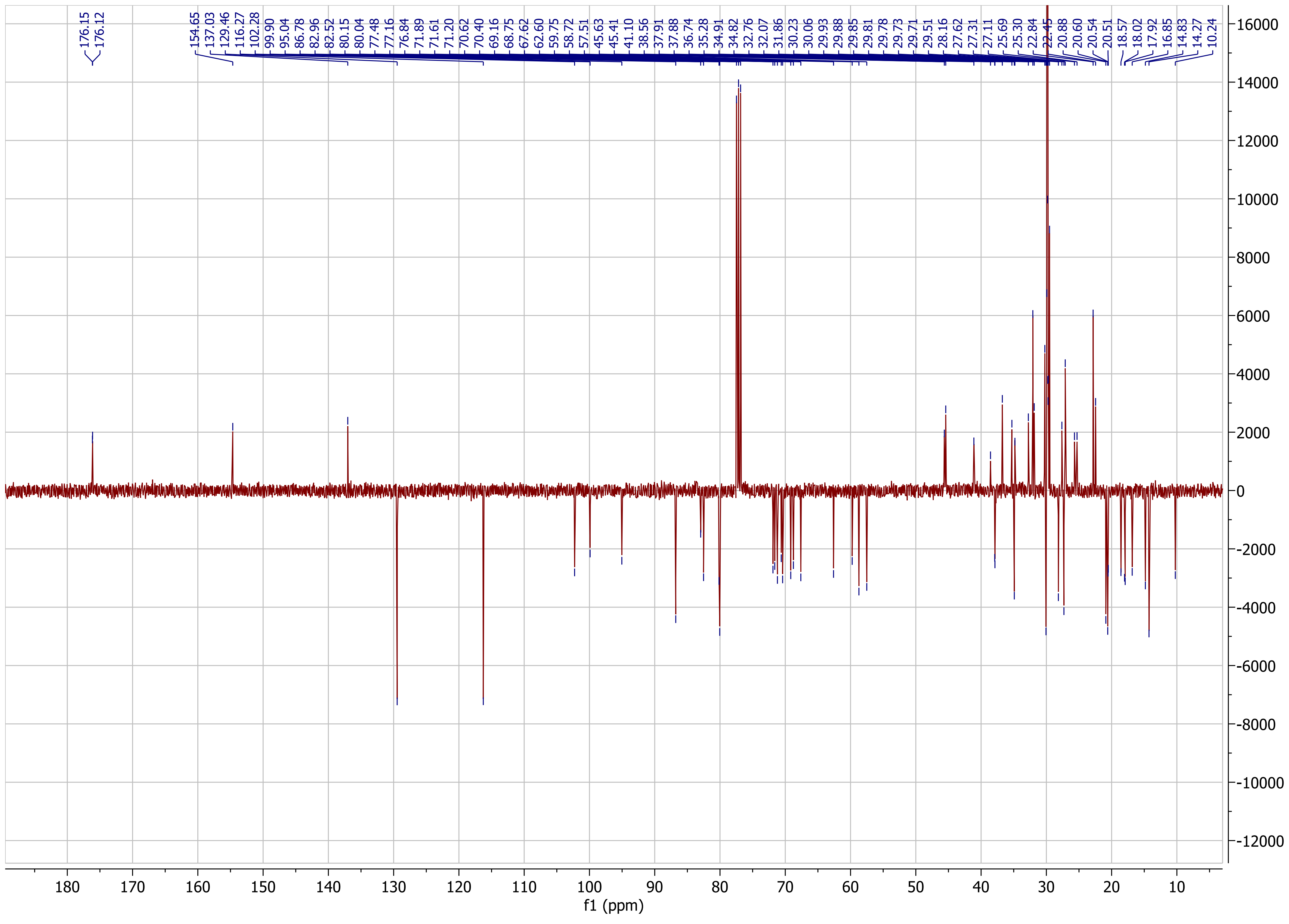

176.15
176.12
154.65
137.03
129.46
116.27
102.28
99.90
95.04
86.78
82.96
82.52
80.15
80.04
77.48
77.16
76.84
71.89
71.61
71.20
70.62
70.40
69.16
68.75
67.62
62.60
59.75
58.72
57.51
45.63
45.41
41.10
38.56
37.91
37.88
36.74
35.28
34.91
34.82
32.76
32.07
31.86
30.23
30.06
29.93
29.88
29.85
29.81
29.78
29.73
29.71
29.51
28.16
27.62
27.31
27.11
25.69
25.30
22.84
22.45
20.88
20.60
20.54
20.51
18.57
18.02
17.92
16.85
14.83
14.27
10.24
f1 (ppm)

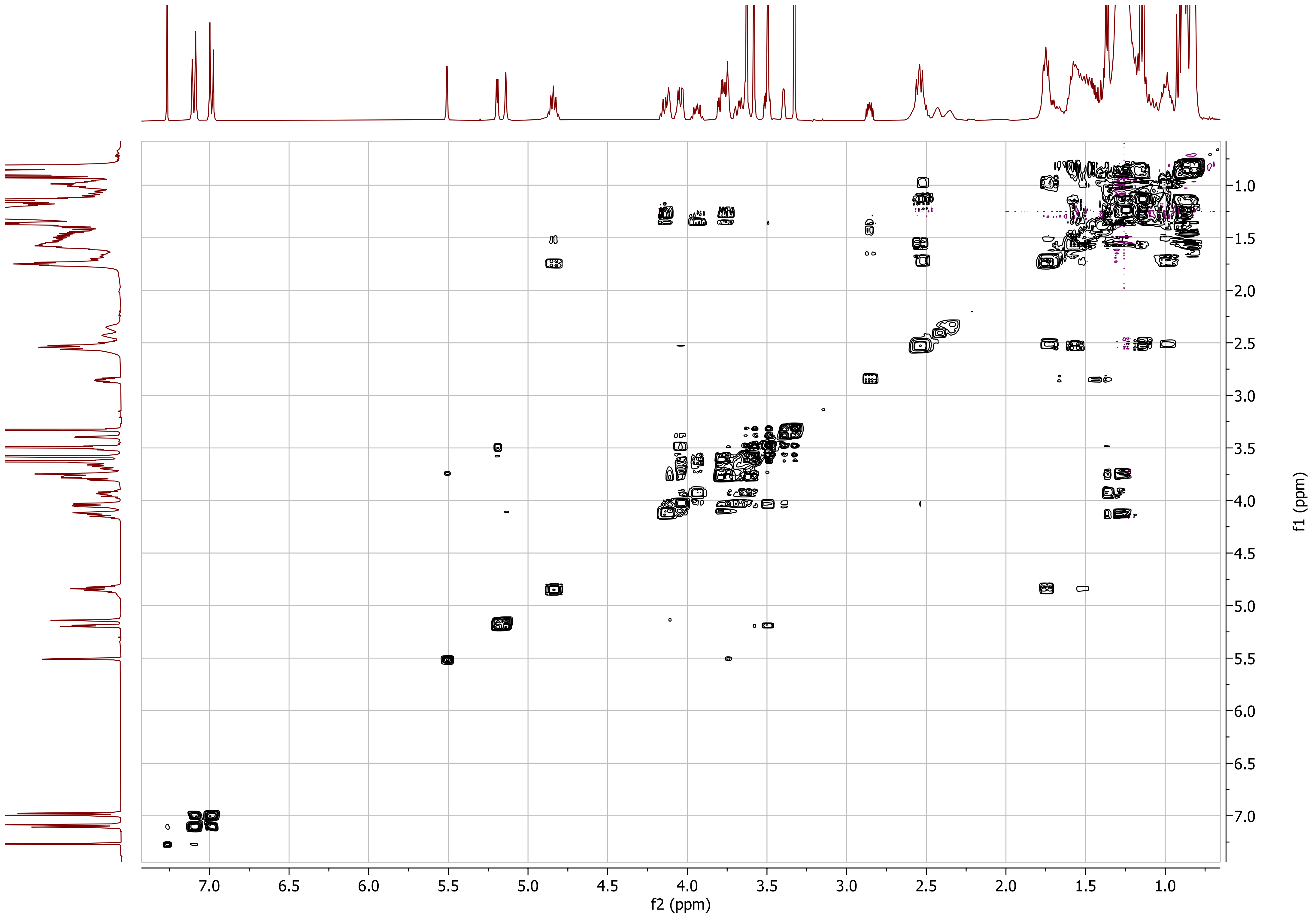

7.0
6.5
6.0
5.5
5.0
4.5
4.0
3.5
3.0
2.5
2.0
1.5
1.0
f2 (ppm)
f1 (ppm)

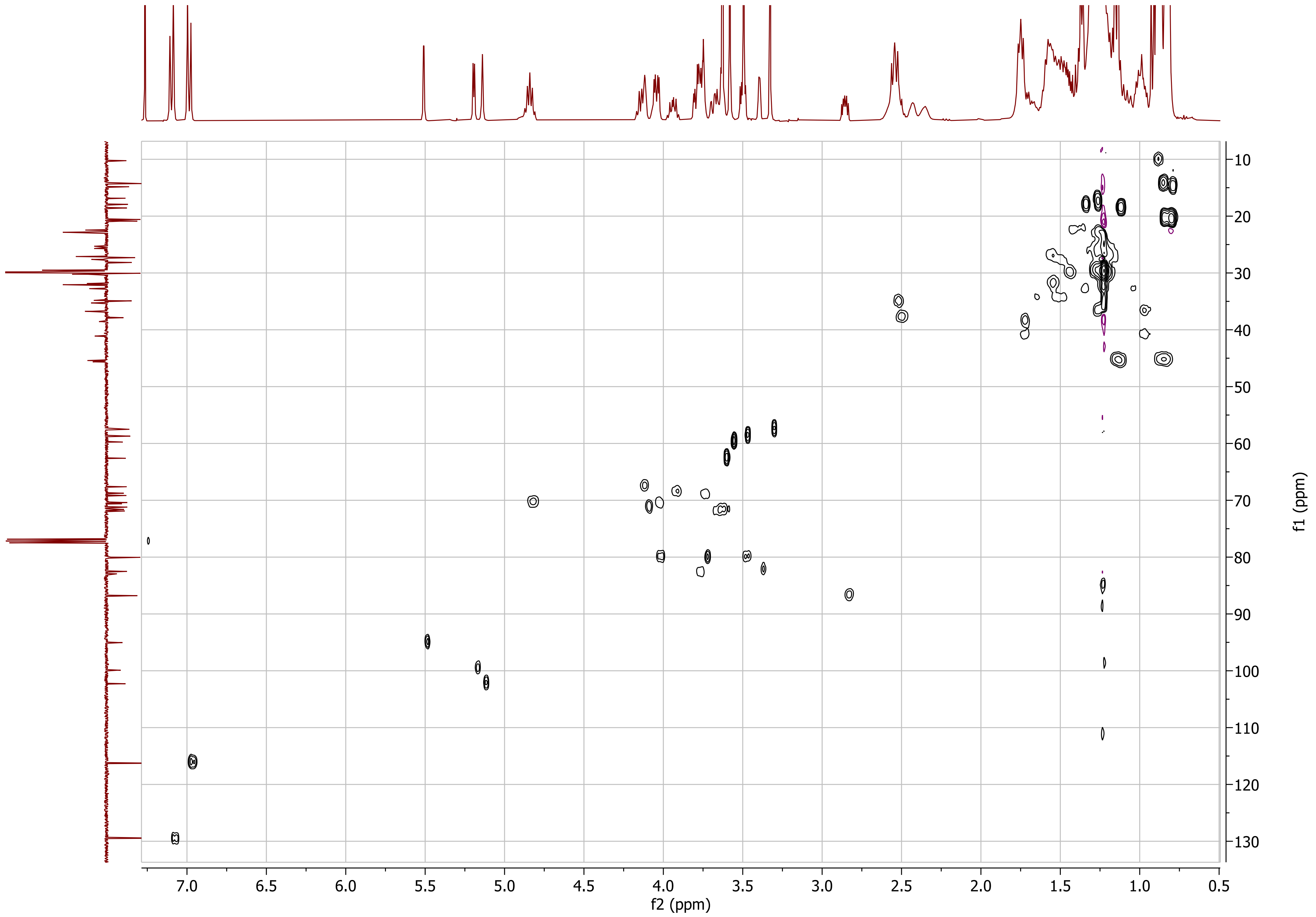

f2 (ppm)
f1 (ppm)
7.0
6.5
6.0
5.5
5.0
4.5
4.0
3.5
3.0
2.5
2.0
1.5
1.0
0.5
10
20
30
40
50
60
70
80
90
100
110
120
130

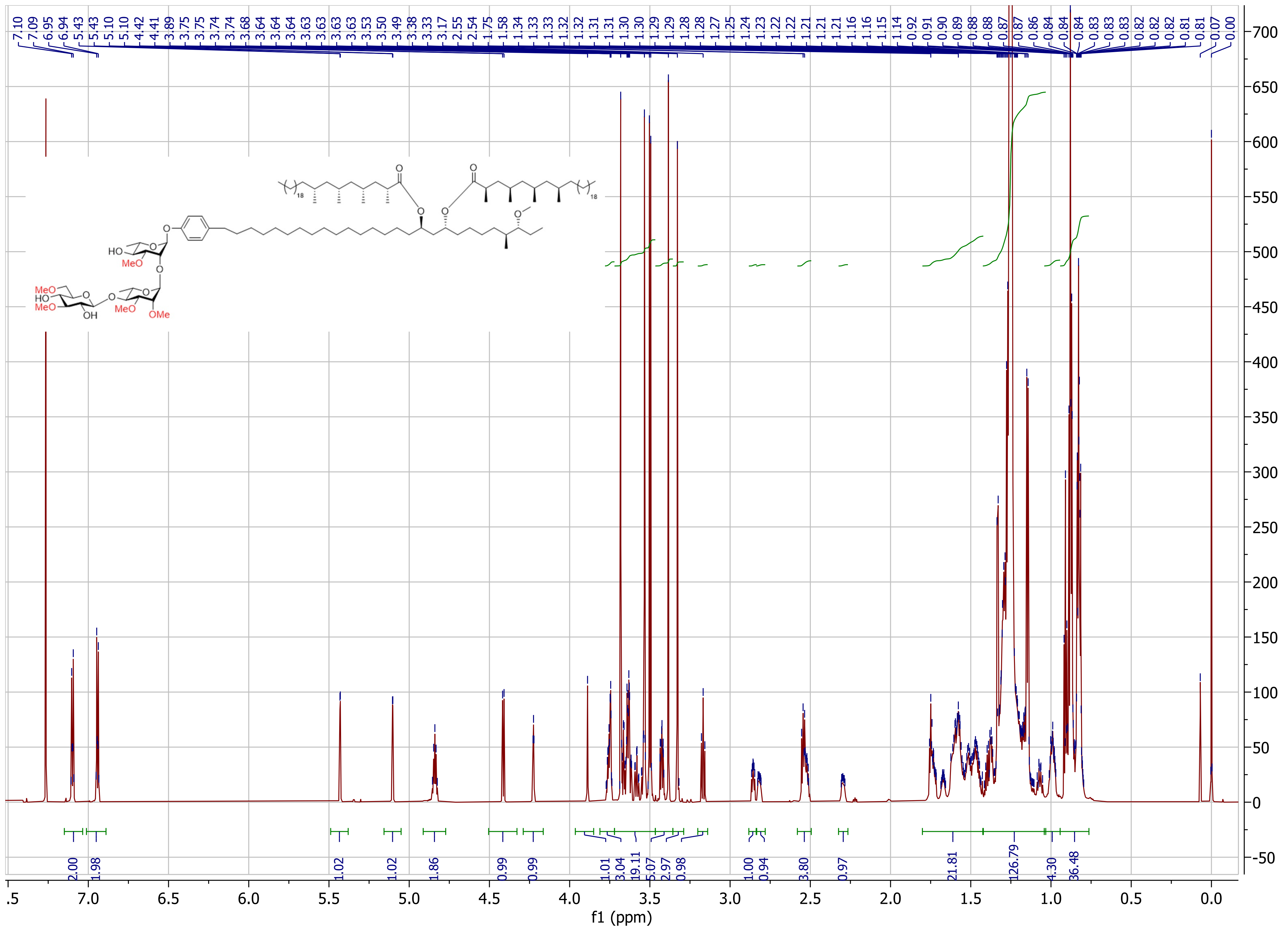

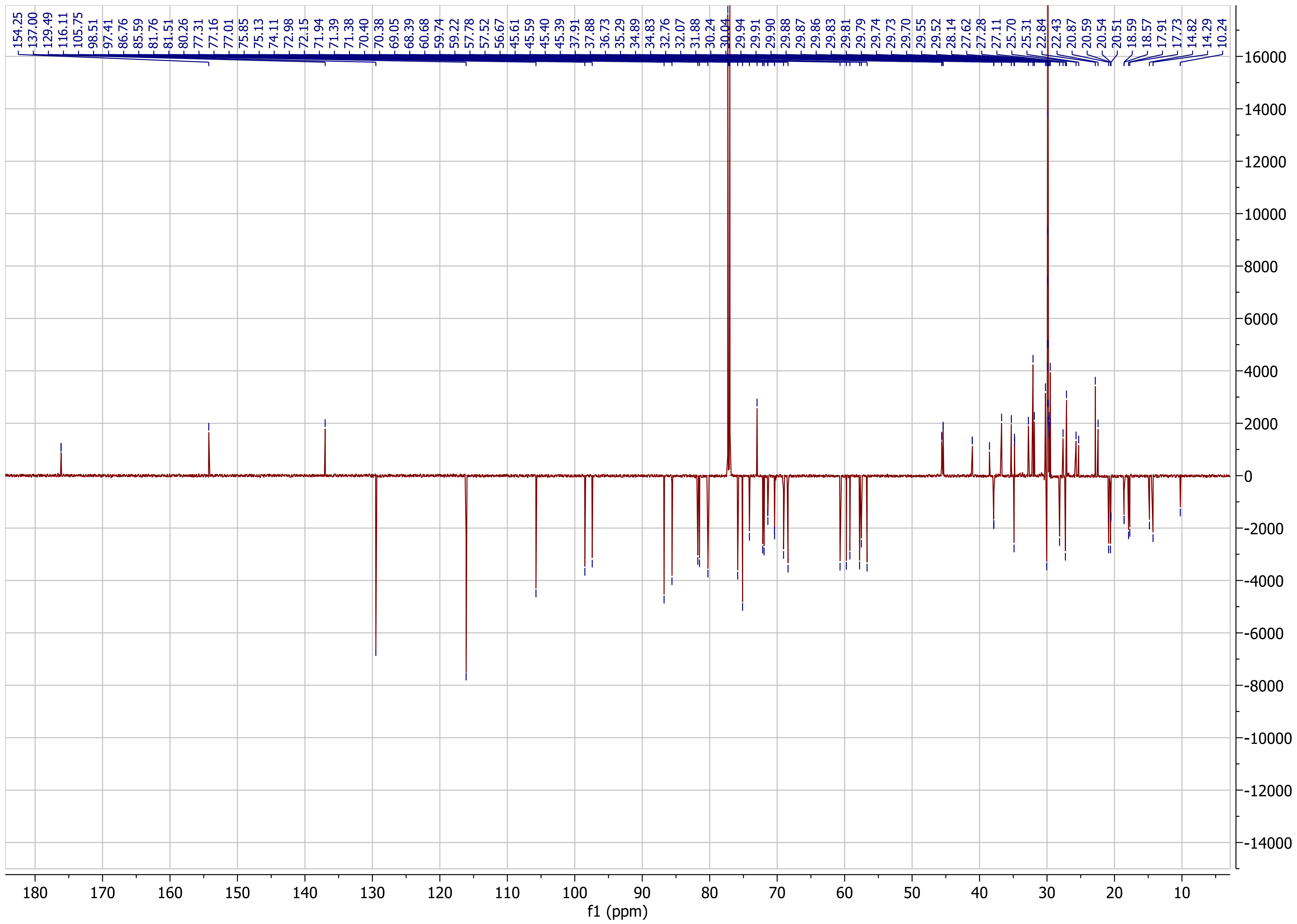
154.25
137.00
129.49
116.11
105.75
98.51
97.41
86.76
85.59
81.76
81.51
80.26
77.31
77.16
77.01
75.85
75.13
74.11
72.98
72.15
71.94
71.39
71.38
70.40
70.38
69.05
68.39
60.68
59.74
59.22
57.78
57.52
56.67
45.61
45.59
45.40
45.39
37.91
37.88
36.73
35.29
34.89
34.83
32.76
32.07
31.88
30.24
30.04
29.94
29.91
29.90
29.88
29.87
29.86
29.83
29.81
29.79
29.74
29.73
29.70
29.55
29.52
28.14
27.62
27.28
27.11
25.70
25.31
22.84
22.43
20.87
20.59
20.54
20.51
18.59
18.57
17.91
17.73
14.82
14.29
10.24
16000
14000
12000
10000
8000
6000
4000
2000
0
-2000
-4000
-6000
-8000
-10000
-12000
-14000
180
170
160
150
140
130
120
110
100
90
80
70
60
50
40
30
20
10
f1 (ppm)

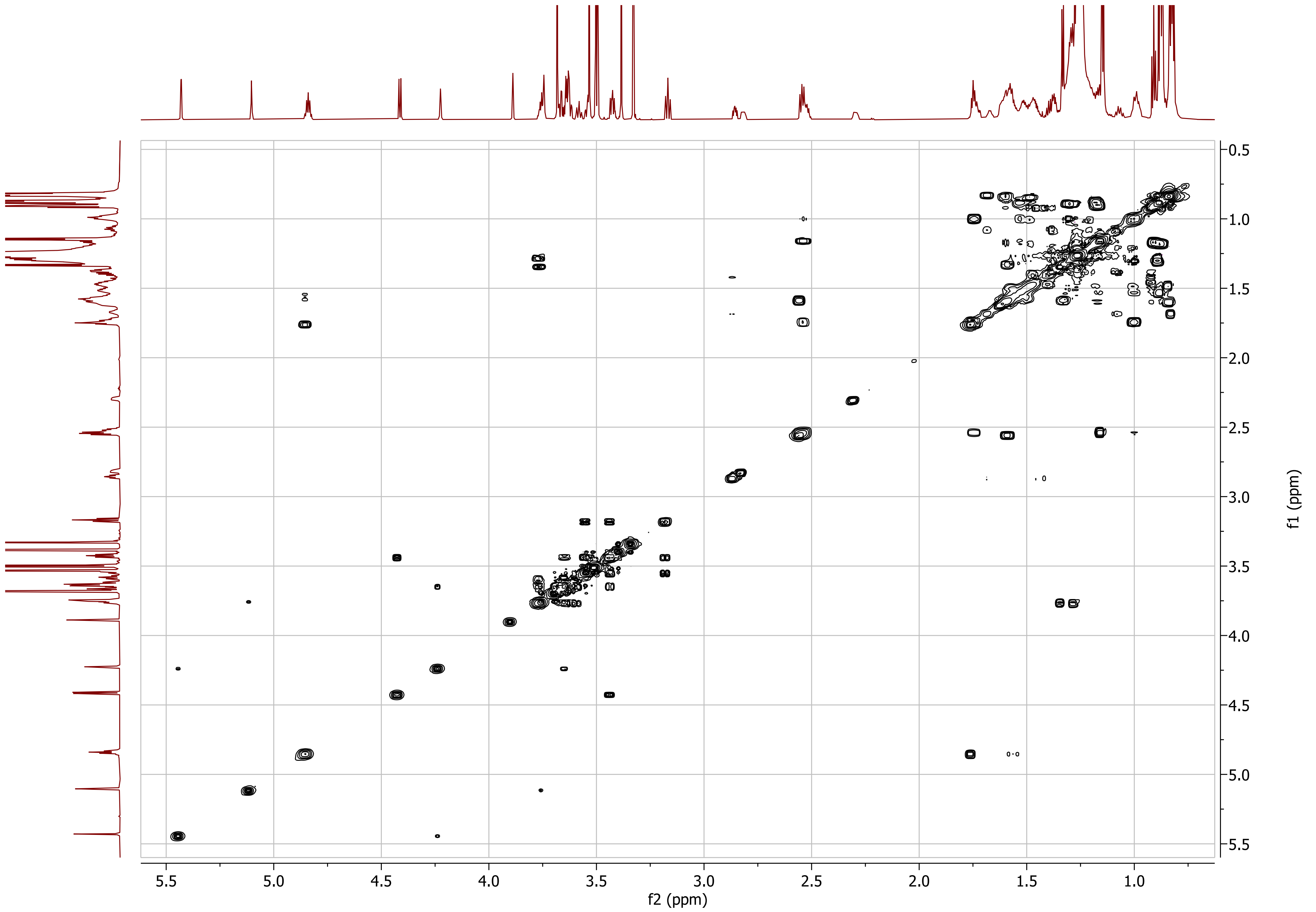

5.5
5.0
4.5
4.0
3.5
3.0
2.5
2.0
1.5
1.0
f2 (ppm)
0.5
1.0
1.5
2.0
2.5
3.0
3.5
4.0
4.5
5.0
5.5
f1 (ppm)

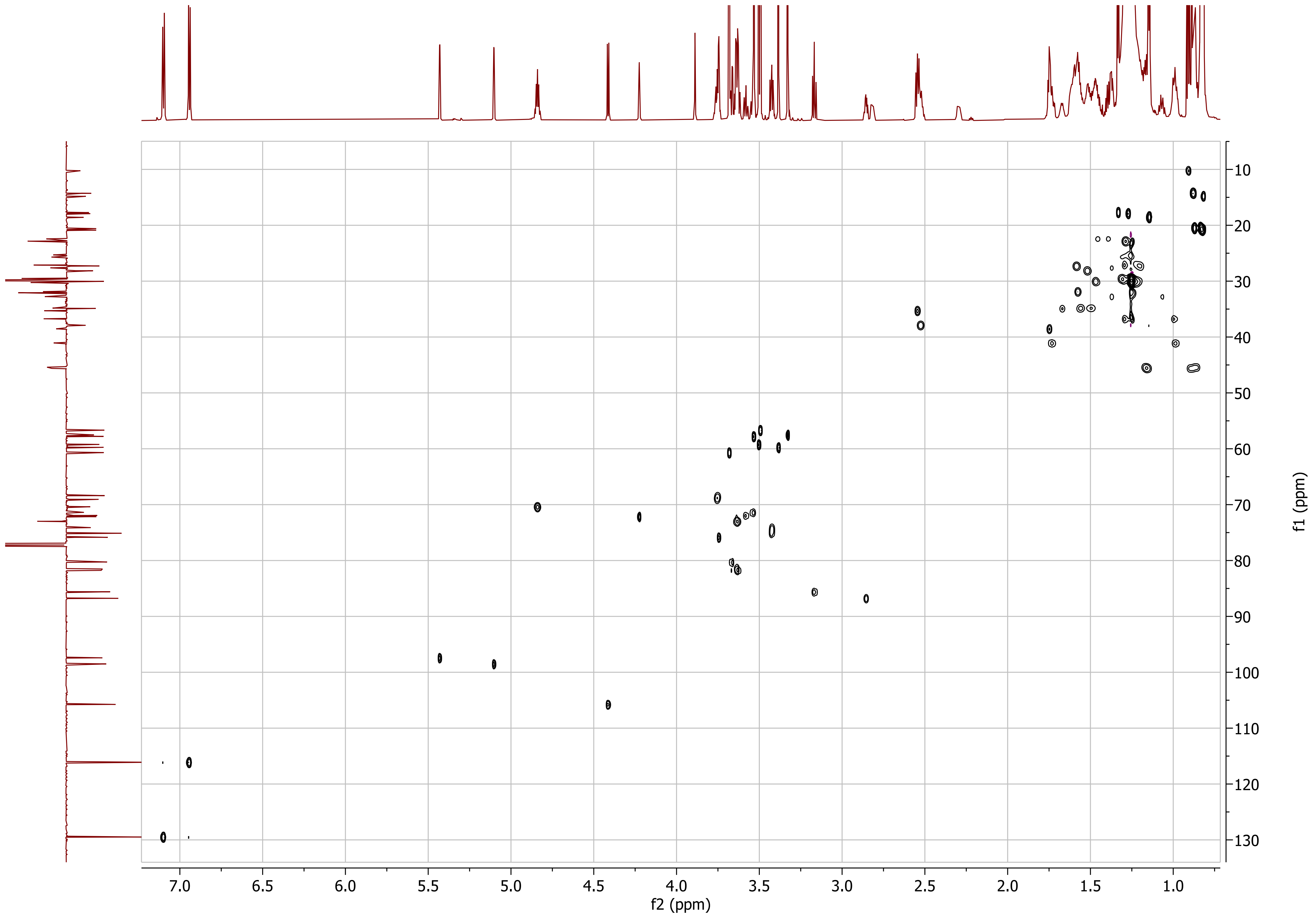

f2 (ppm)
f1 (ppm)
7.0
6.5
6.0
5.5
5.0
4.5
4.0
3.5
3.0
2.5
2.0
1.5
1.0
10
20
30
40
50
60
70
80
90
100
110
120
130

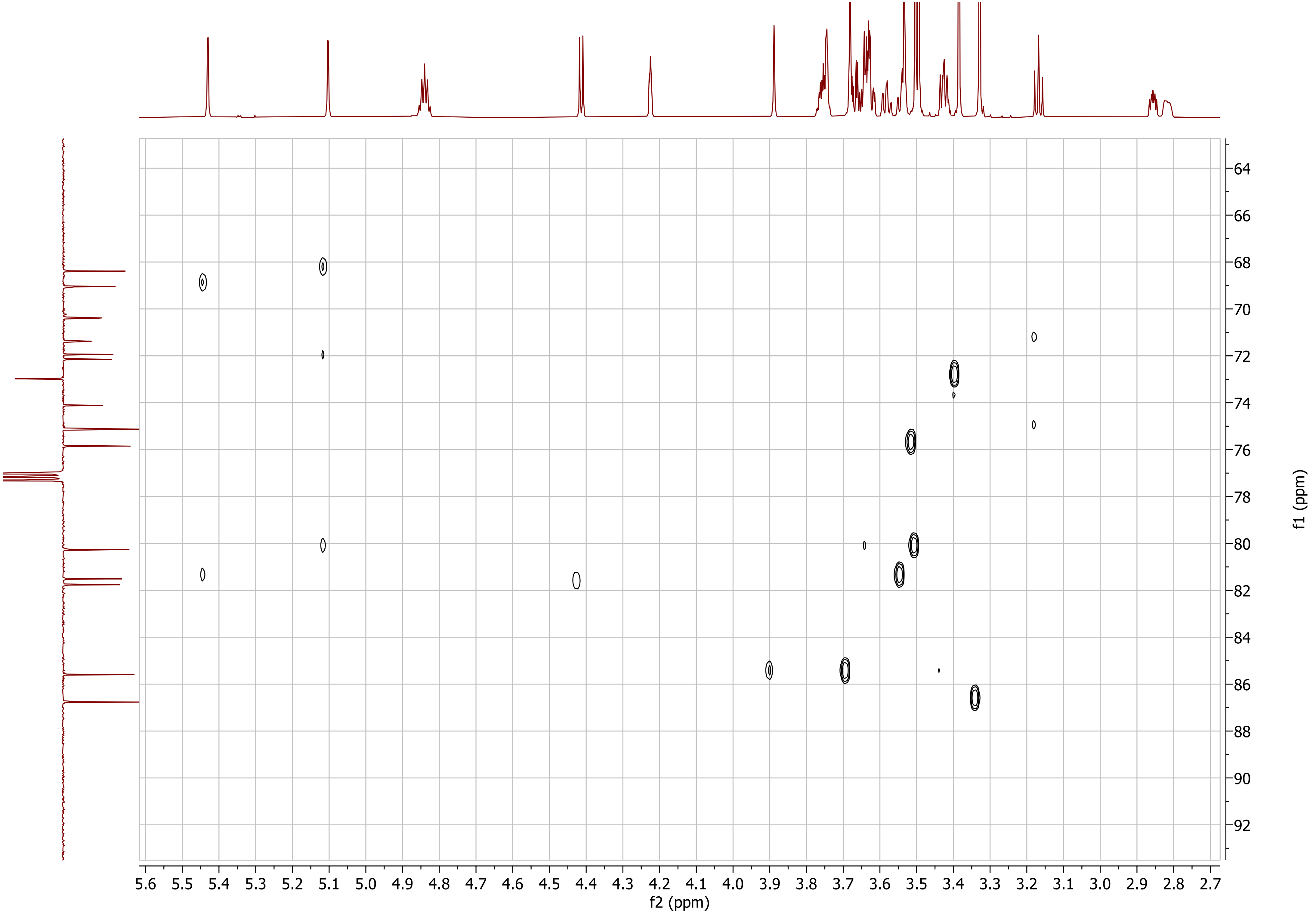

5.6 5.5 5.4 5.3 5.2 5.1 5.0 4.9 4.8 4.7 4.6 4.5 4.4 4.3 4.2 4.1 4.0 3.9 3.8 3.7 3.6 3.5 3.4 3.3 3.2 3.1 3.0 2.9 2.8 2.7
f2 (ppm)
64 66 68 70 72 74 76 78 80 82 84 86 88 90 92
f1 (ppm)

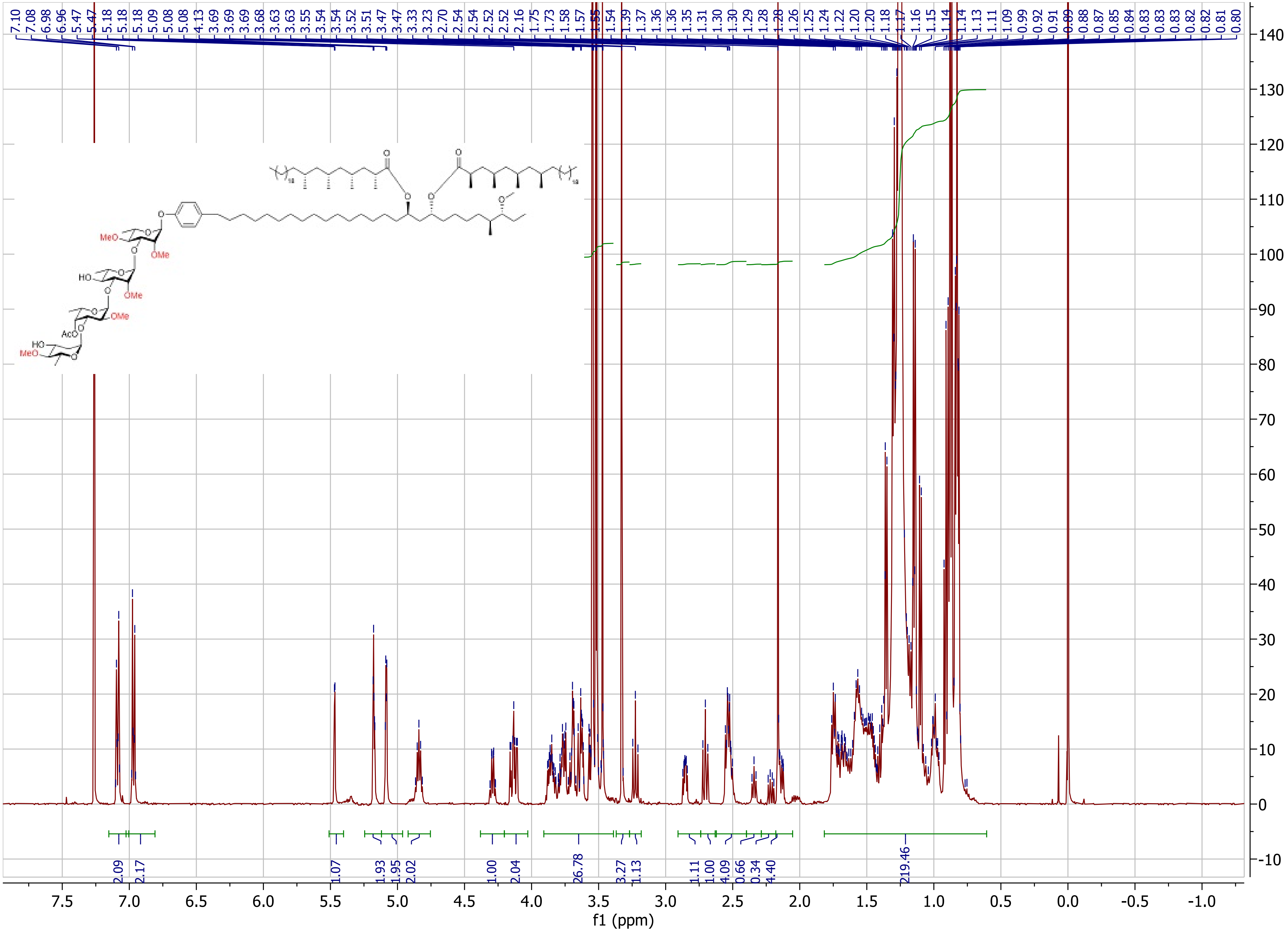

MeO
OMe
HO
OMe
OMe
AcO
HO
MeO
f1 (ppm)

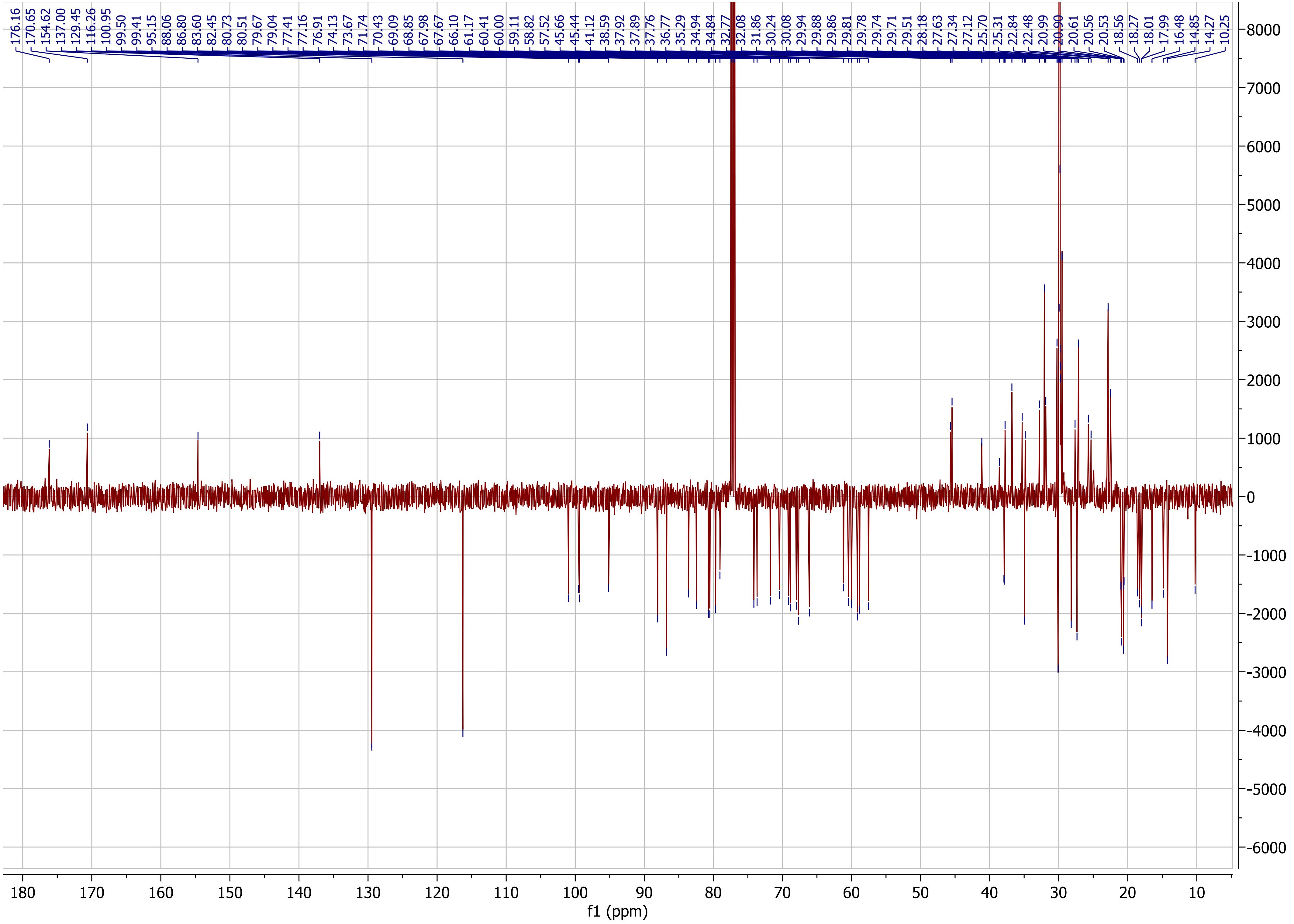
176.16
170.65
154.62
137.00
129.45
116.26
100.95
99.50
99.41
95.15
88.06
86.80
83.60
82.45
80.73
80.51
79.67
79.04
77.41
77.16
76.91
74.13
73.67
71.74
70.43
69.09
68.85
67.98
67.67
66.10
61.17
60.41
60.00
59.11
58.82
57.52
45.66
45.44
41.12
38.59
37.92
37.89
37.76
36.77
35.29
34.94
34.84
32.77
32.08
31.86
30.24
30.08
29.94
29.88
29.86
29.81
29.78
29.74
29.71
29.51
28.18
27.63
27.34
27.12
25.70
25.31
22.84
22.48
20.99
20.90
20.61
20.56
20.53
18.56
18.27
18.01
17.99
16.48
14.85
14.27
10.25
8000
7000
6000
5000
4000
3000
2000
1000
0
-1000
-2000
-3000
-4000
-5000
-6000
180
170
160
150
140
130
120
110
100
90
80
70
60
50
40
30
20
10
f1 (ppm)

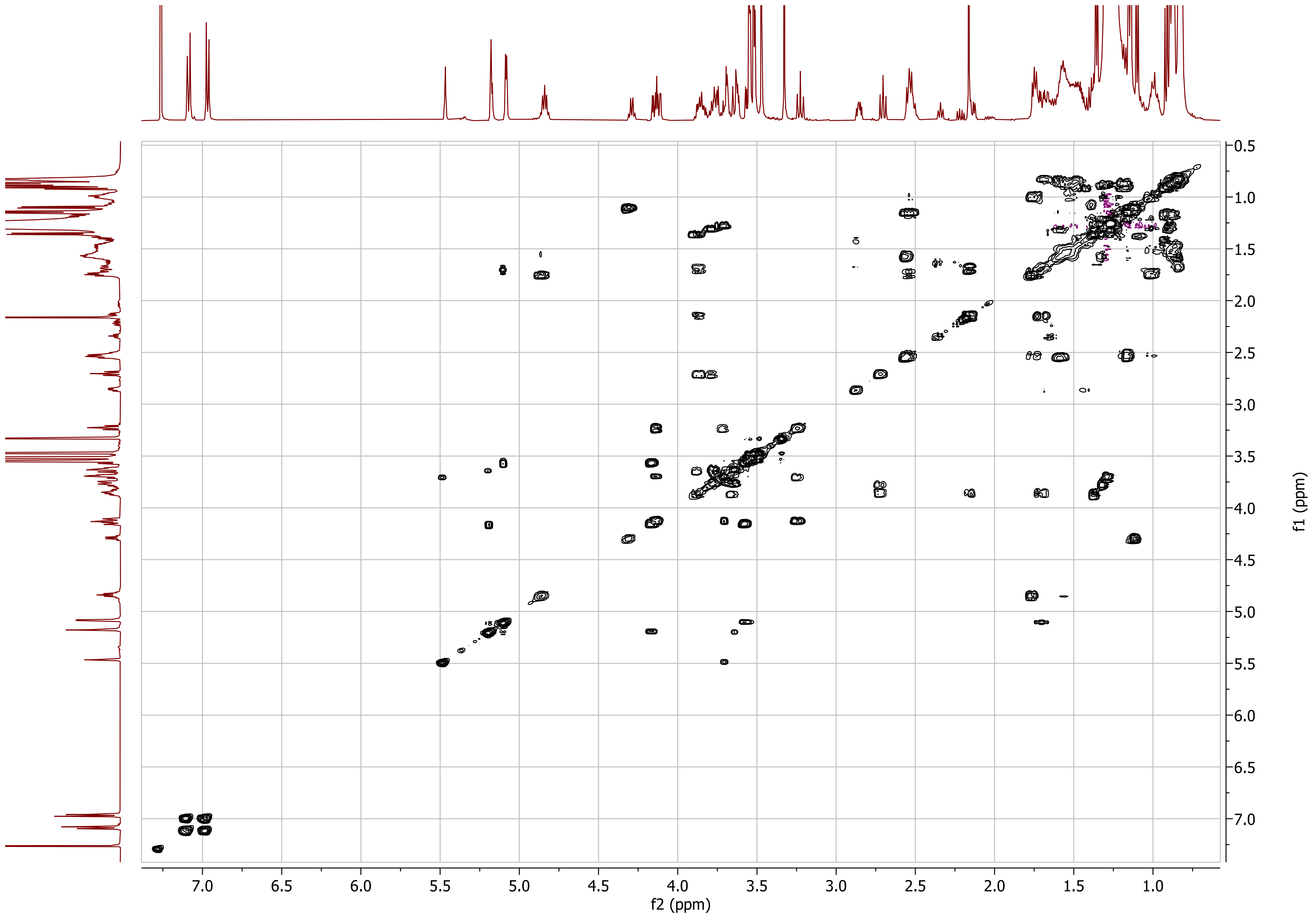

f2 (ppm)
f1 (ppm)

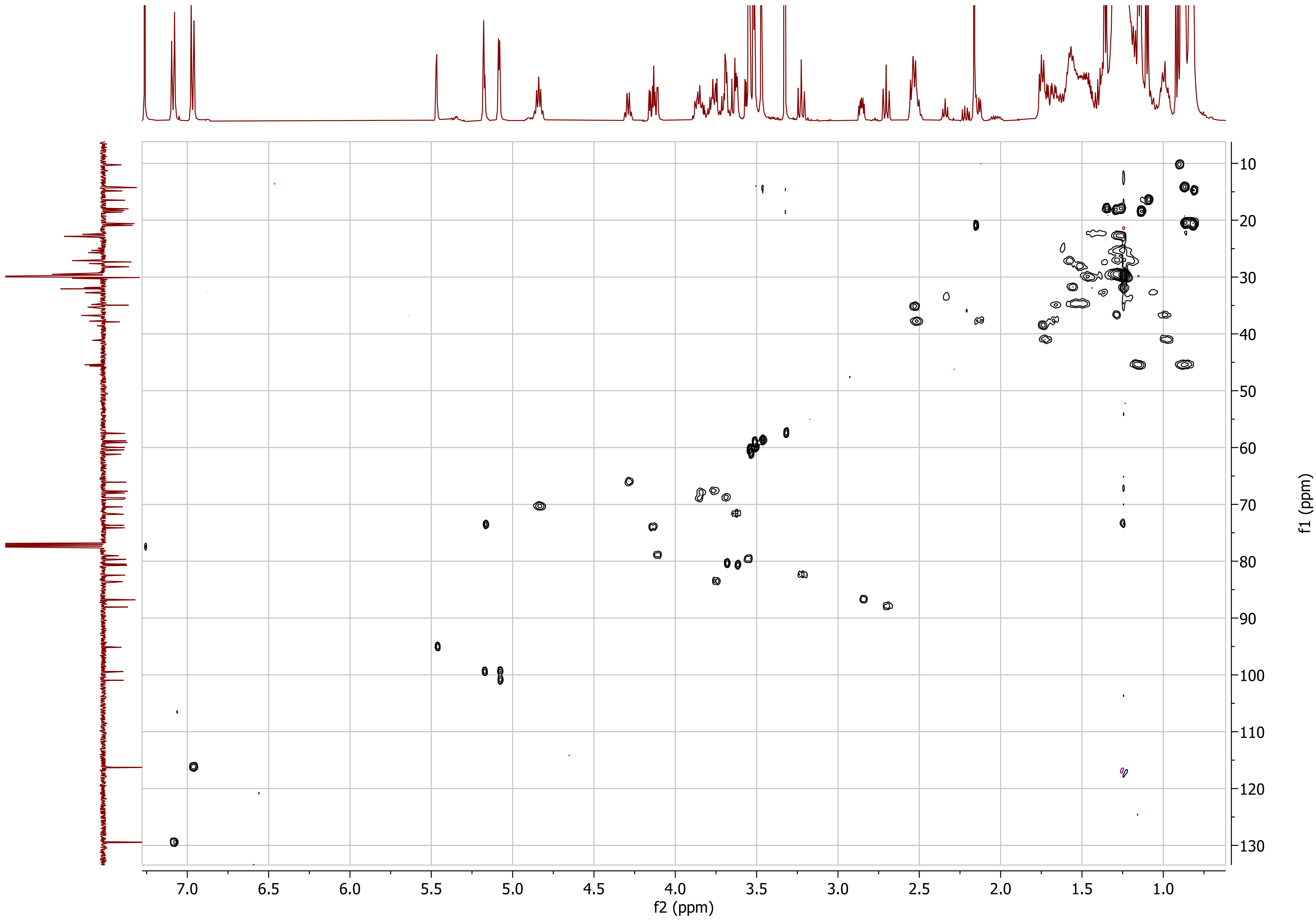

f2 (ppm)
f1 (ppm)

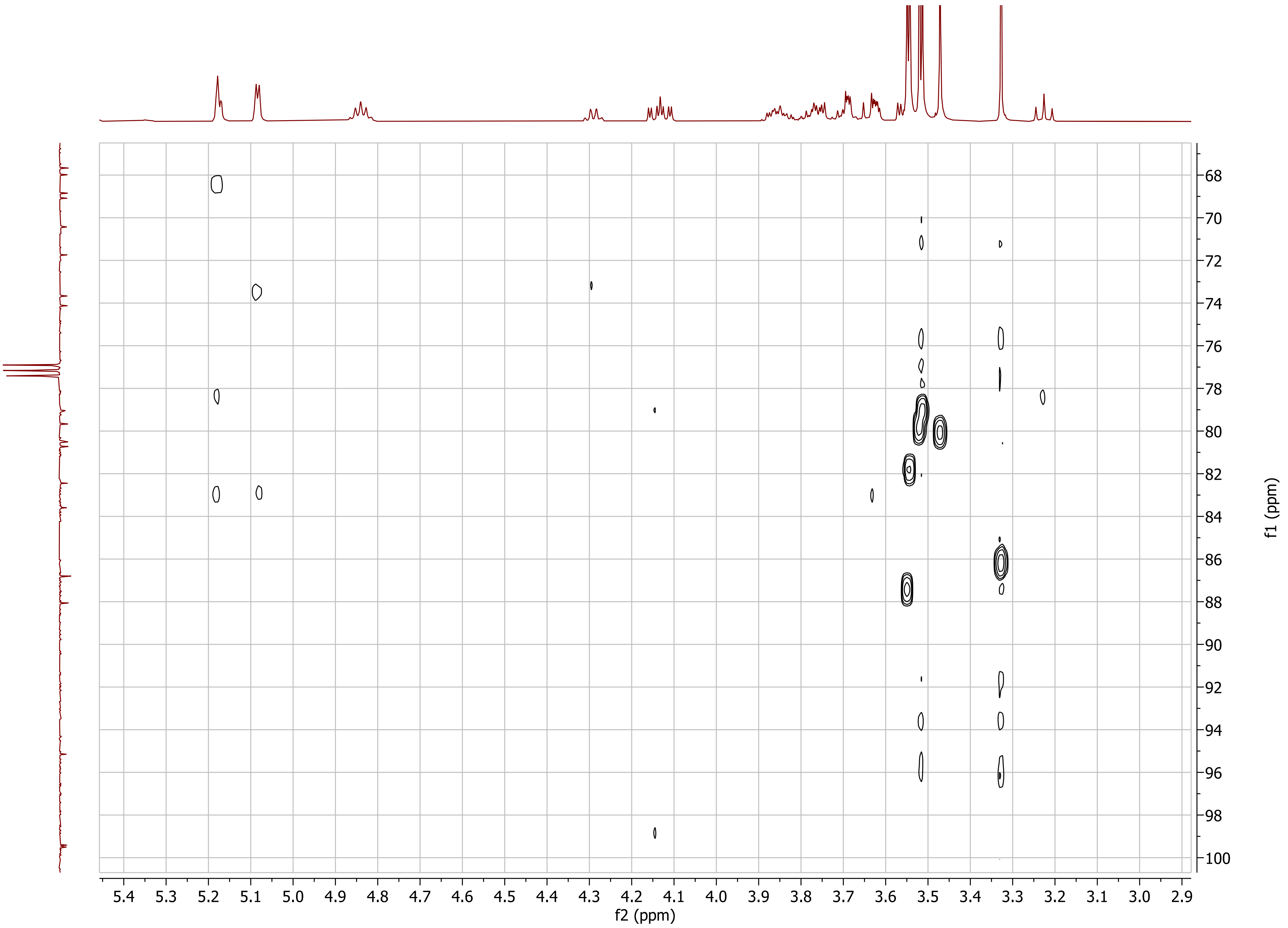

f2 (ppm)
f1 (ppm)

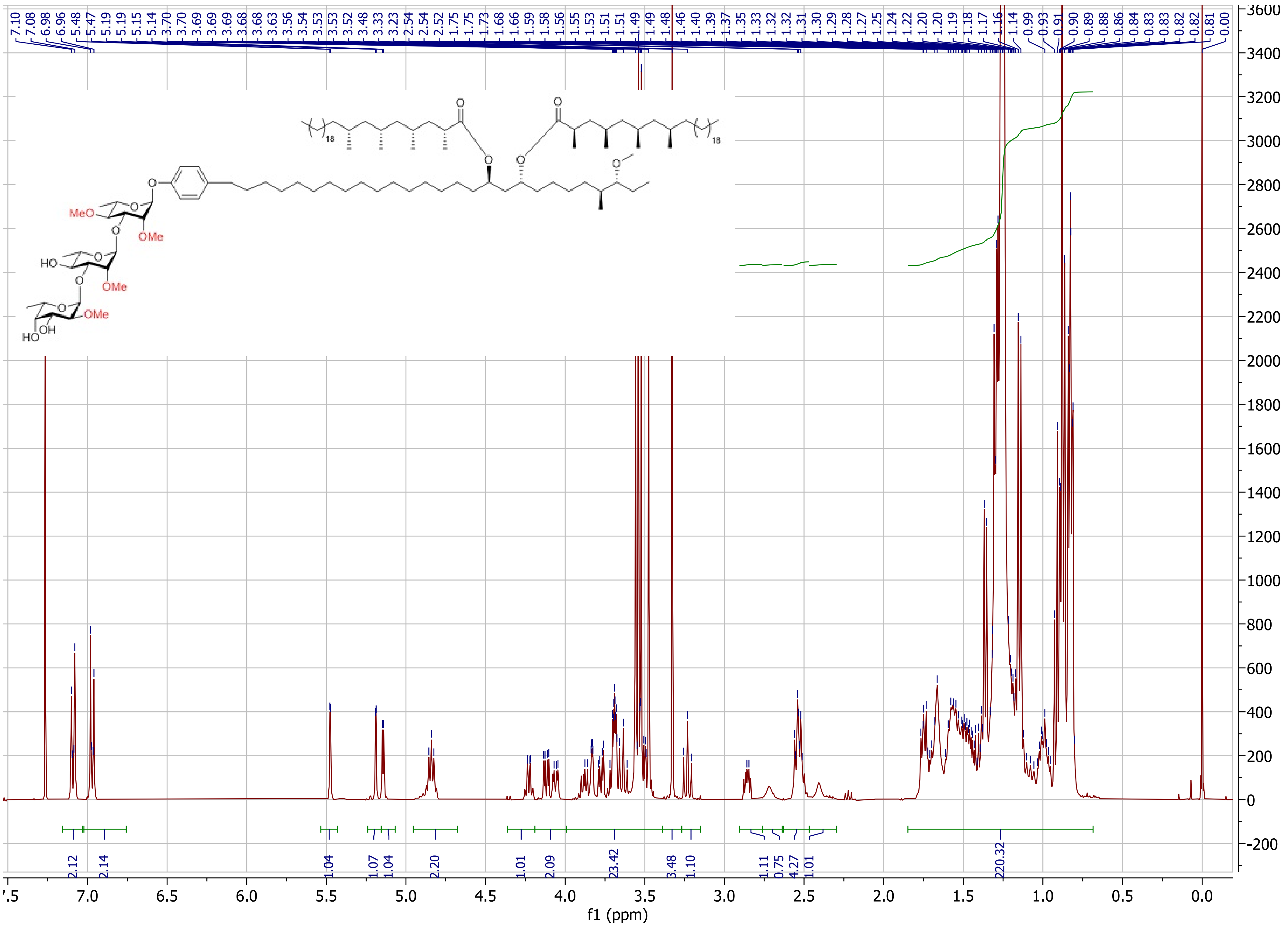

f1 (ppm)
MeO
OMe
HO
OMe
OMe
HO
OH
18
18

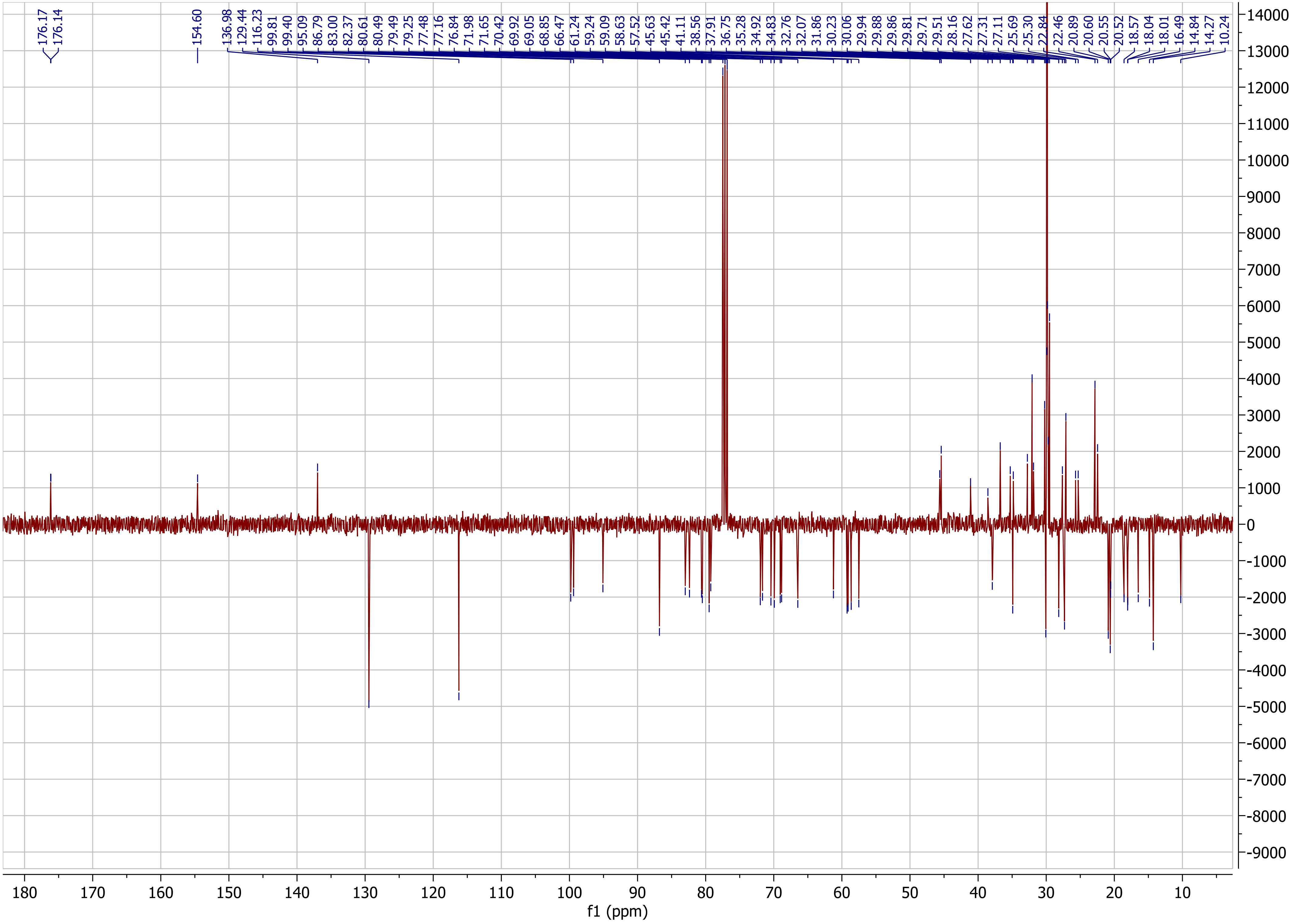
176.17
176.14
154.60
136.98
129.44
116.23
99.81
99.40
95.09
86.79
83.00
82.37
80.61
80.49
79.49
79.25
77.48
77.16
76.84
71.98
71.65
70.42
69.92
69.05
68.85
66.47
61.24
59.24
59.09
58.63
57.52
45.63
45.42
41.11
38.56
37.91
36.75
35.28
34.92
34.83
32.76
32.07
31.86
30.23
30.06
29.94
29.88
29.86
29.81
29.71
29.51
28.16
27.62
27.31
27.11
25.69
25.30
22.84
22.46
20.89
20.60
20.55
20.52
18.57
18.04
18.01
16.49
14.84
14.27
10.24
f1 (ppm)

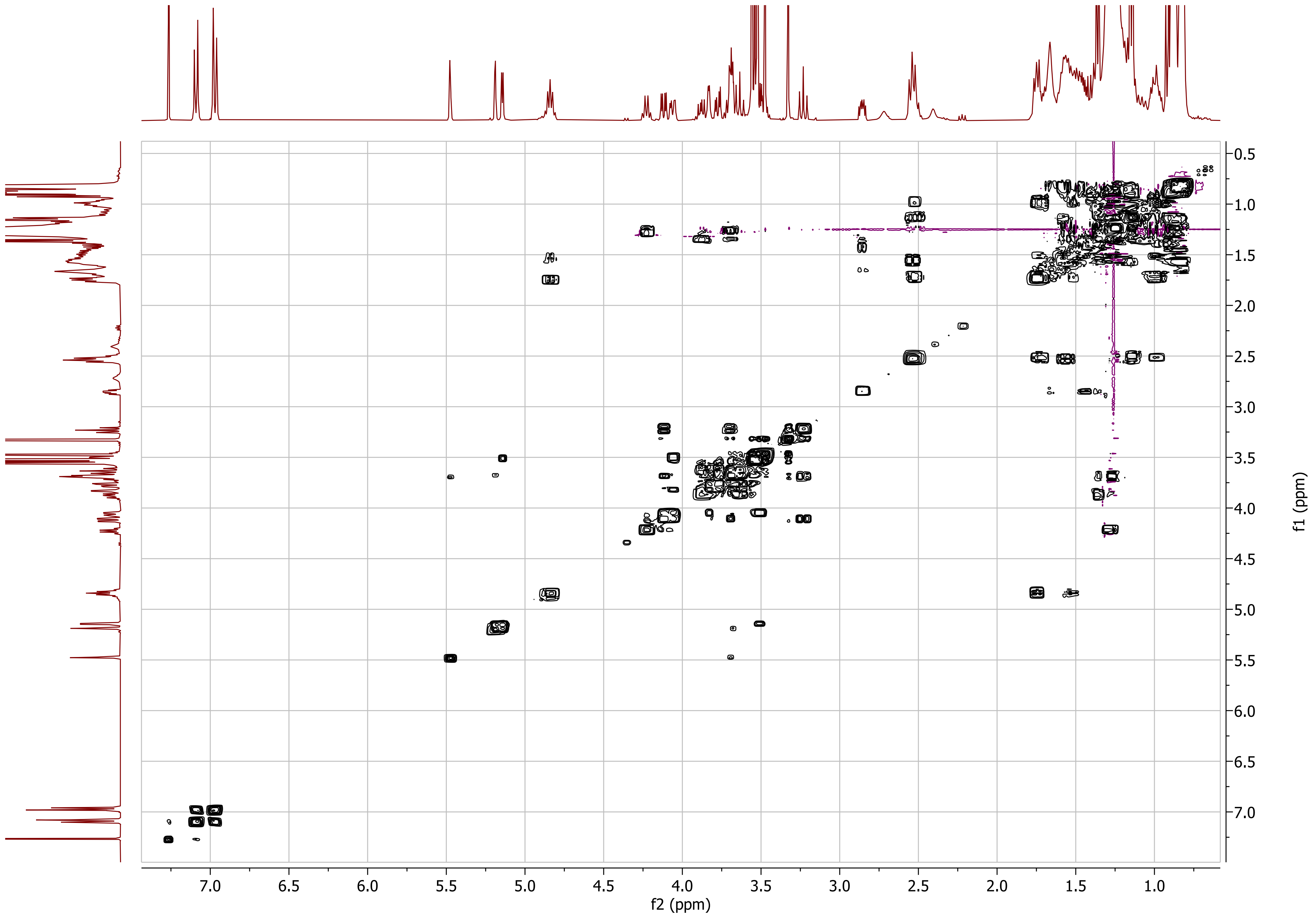

f2 (ppm)
f1 (ppm)

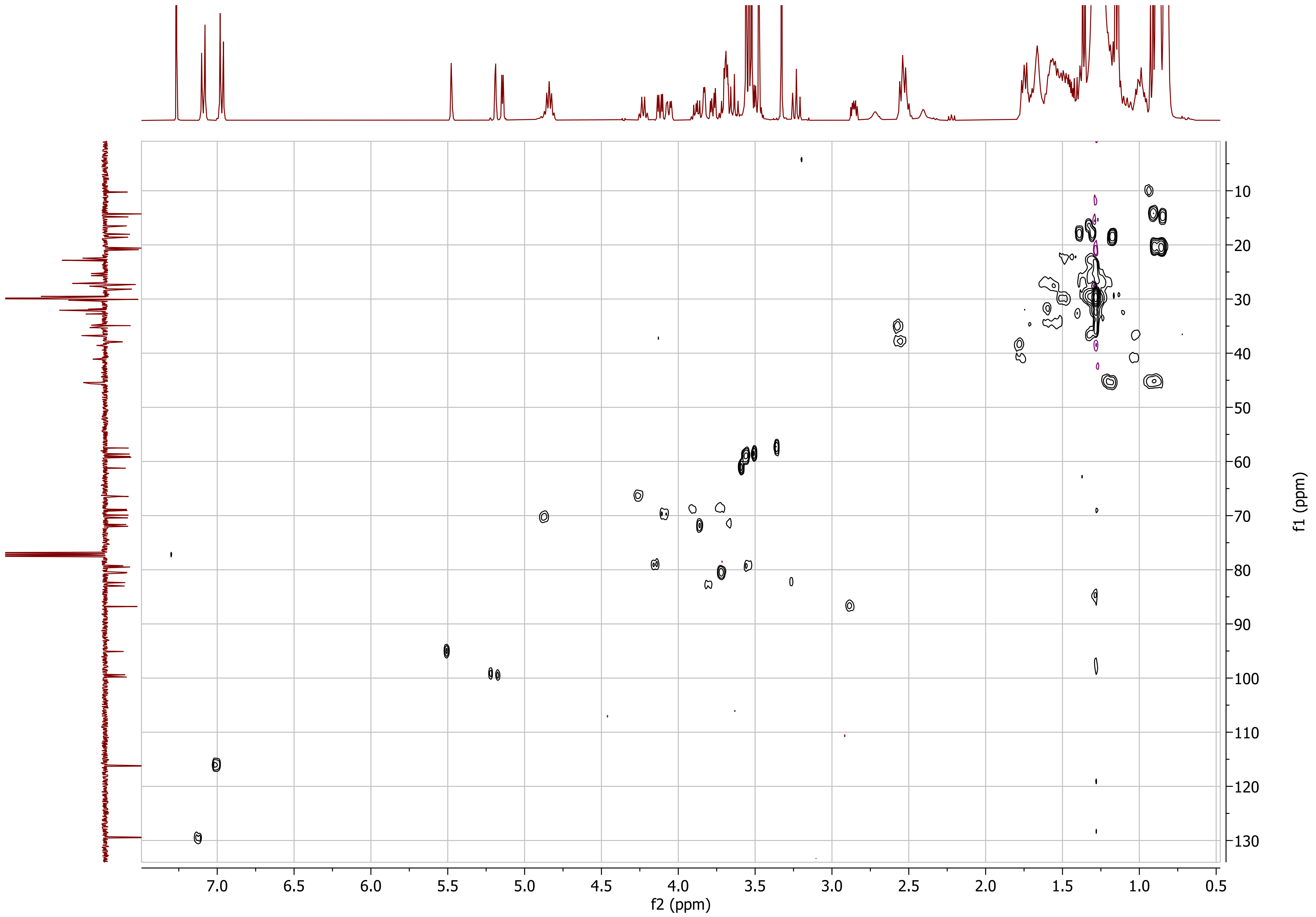
f2 (ppm)
f1 (ppm)

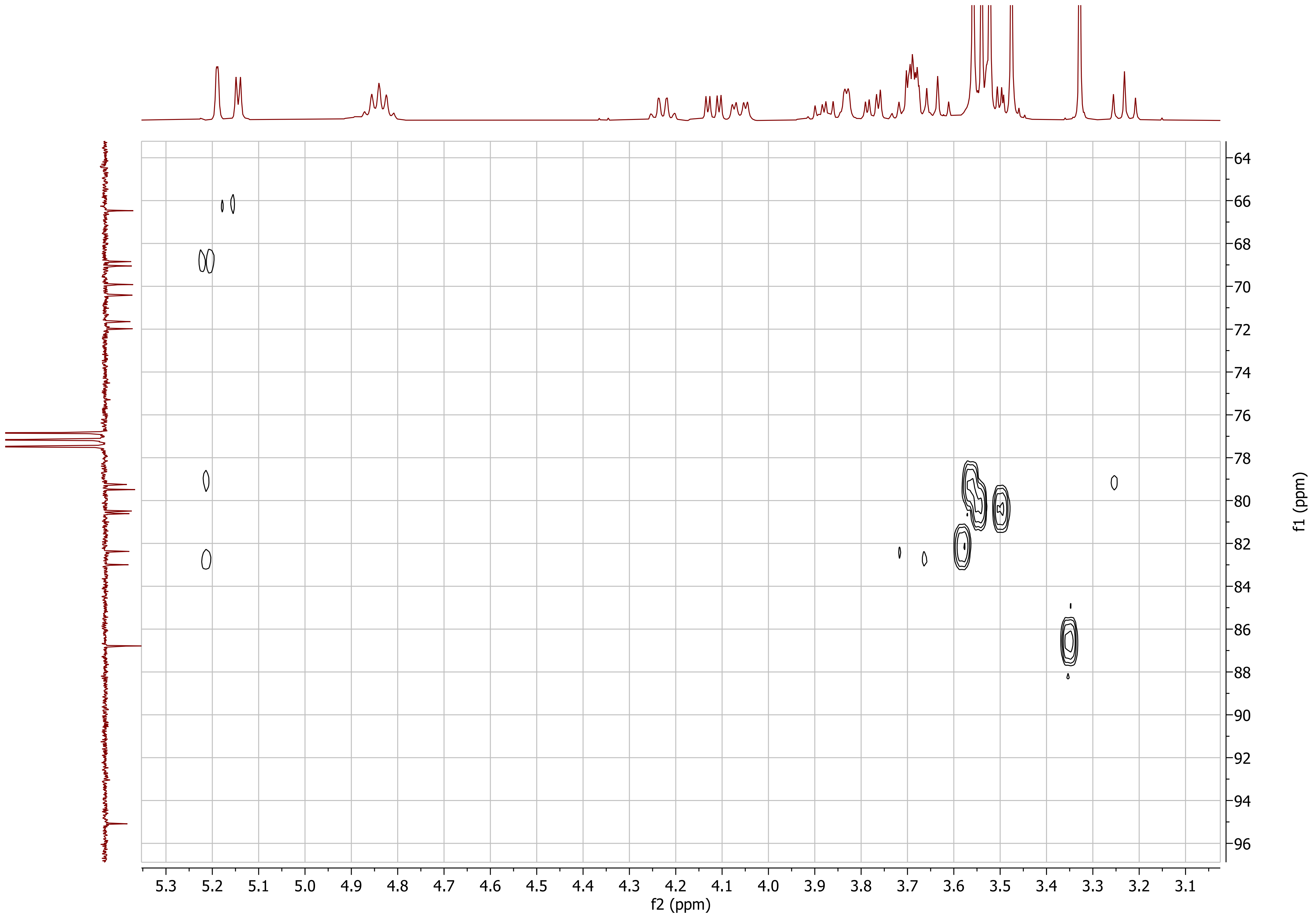

5.3 5.2 5.1 5.0 4.9 4.8 4.7 4.6 4.5 4.4 4.3 4.2 4.1 4.0 3.9 3.8 3.7 3.6 3.5 3.4 3.3 3.2 3.1
f2 (ppm)
64 66 68 70 72 74 76 78 80 82 84 86 88 90 92 94 96
f1 (ppm)

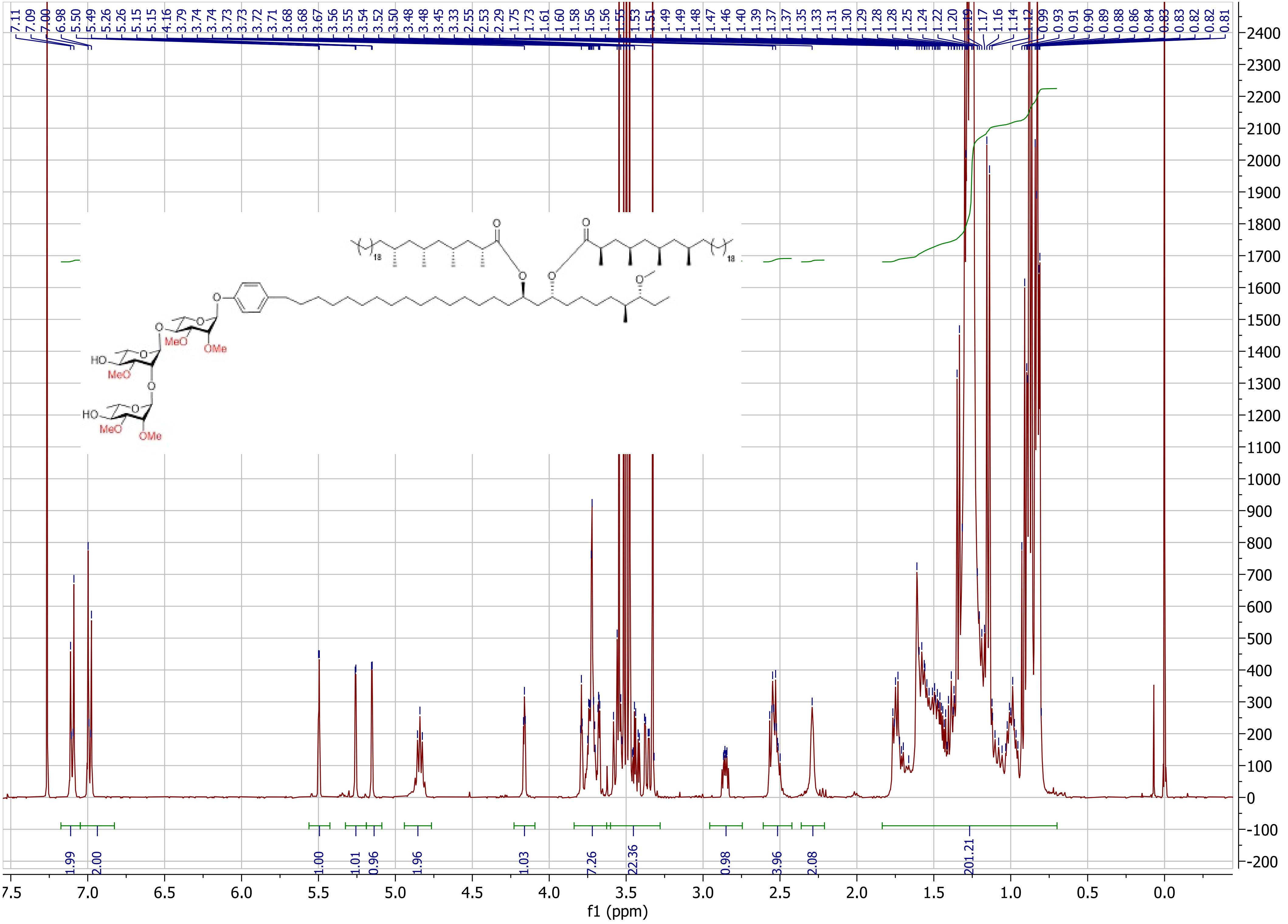

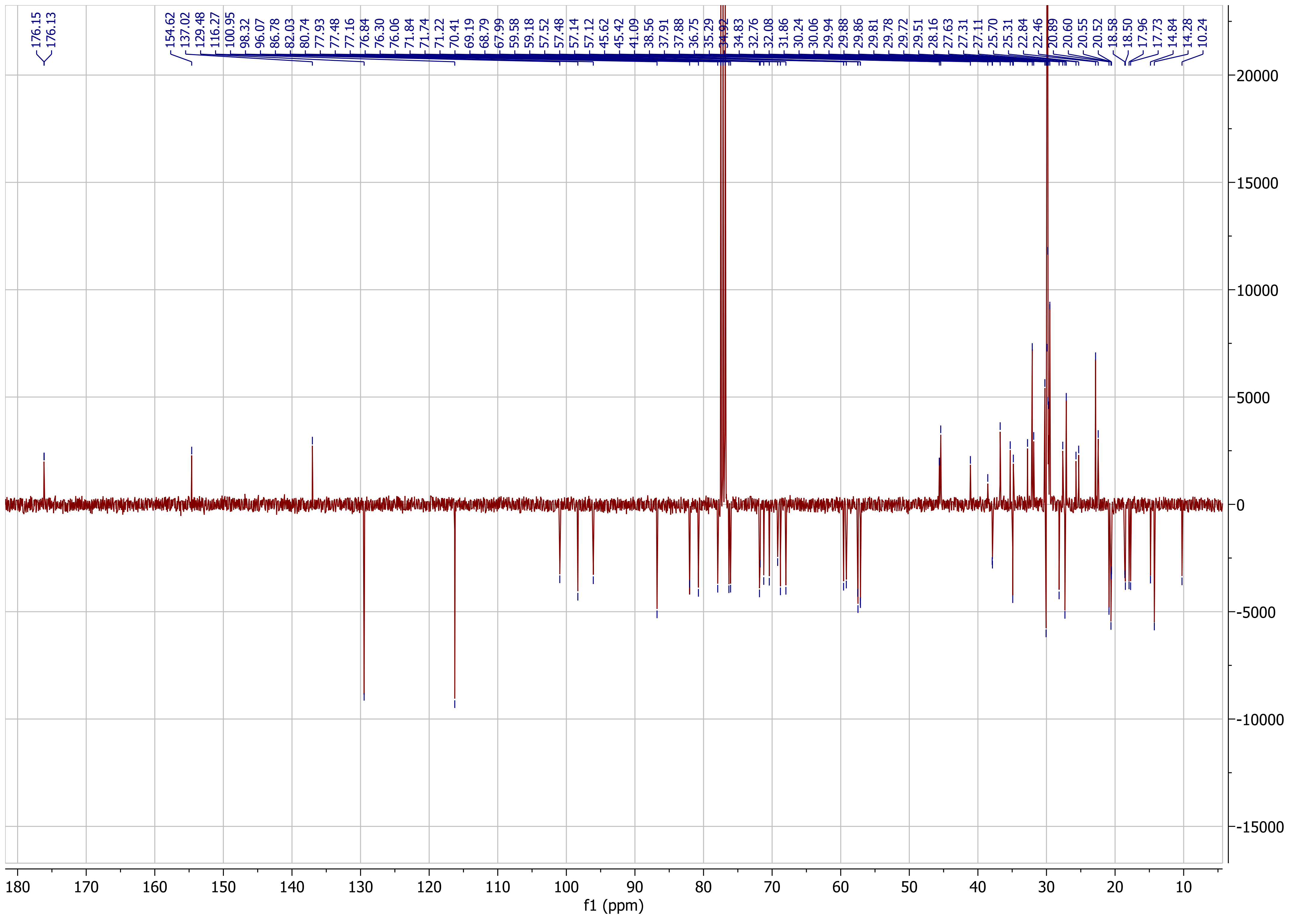
176.15
176.13
154.62
137.02
129.48
116.27
100.95
98.32
96.07
86.78
82.03
80.74
77.93
77.48
77.16
76.84
76.30
76.06
71.84
71.74
71.22
70.41
69.19
68.79
67.99
59.58
59.18
57.52
57.48
57.14
57.12
45.62
45.42
41.09
38.56
37.91
37.88
36.75
35.29
34.92
34.83
32.76
32.08
31.86
30.24
30.06
29.94
29.88
29.86
29.81
29.78
29.72
29.51
28.16
27.63
27.31
27.11
25.70
25.31
22.84
22.46
20.89
20.60
20.55
20.52
18.58
18.50
17.96
17.73
14.84
14.28
10.24
f1 (ppm)

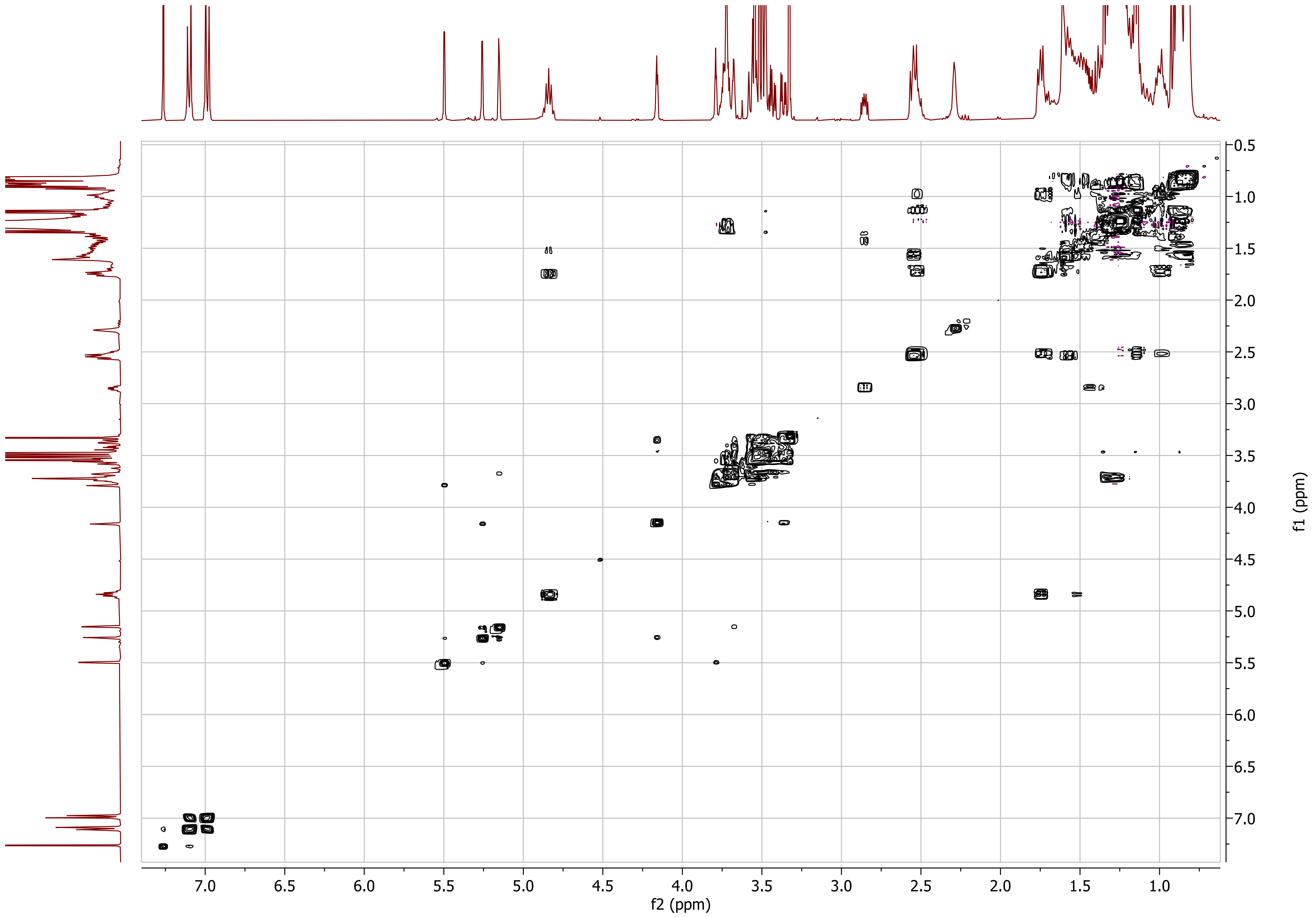

f2 (ppm)
f1 (ppm)

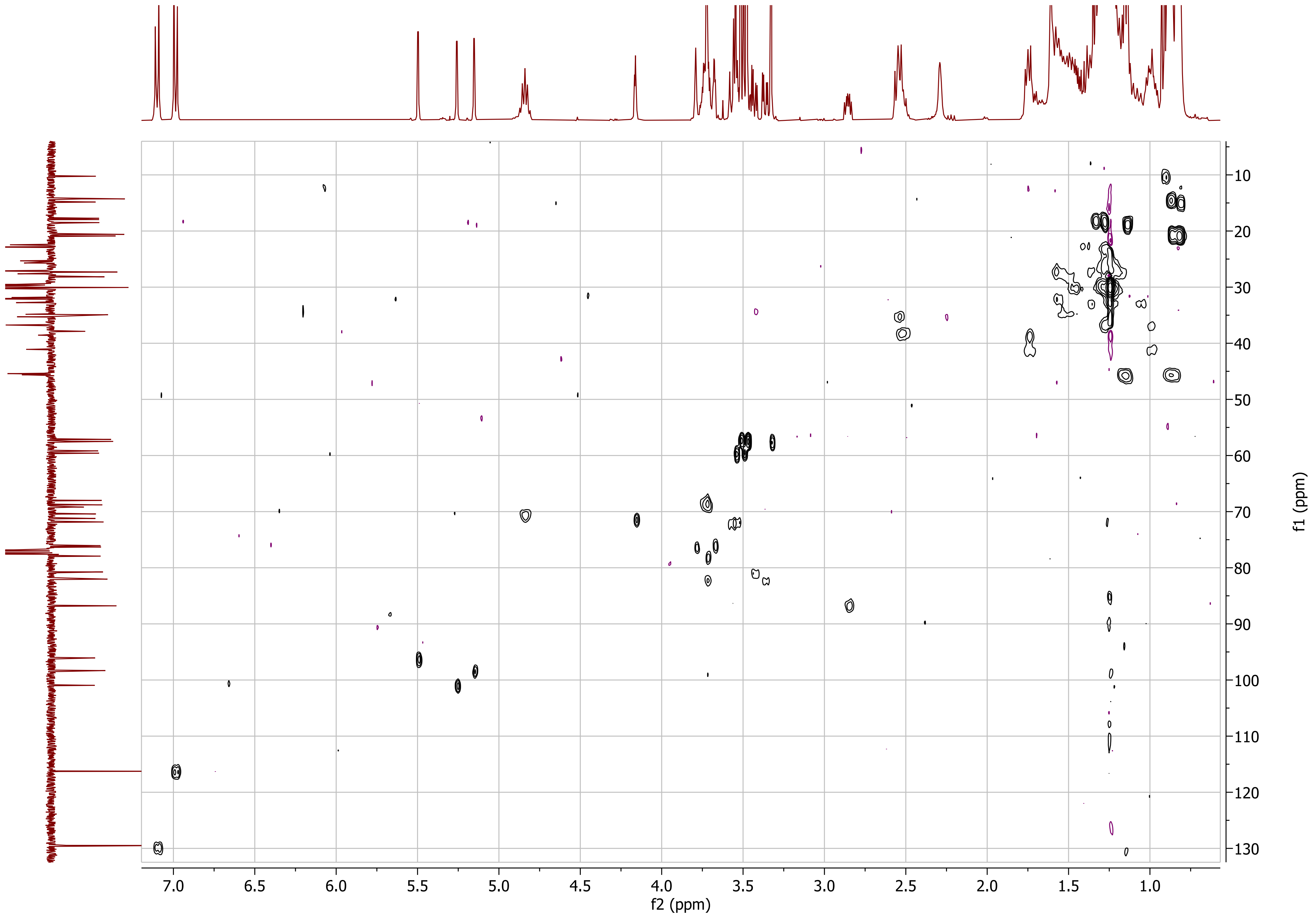

7.0
6.5
6.0
5.5
5.0
4.5
4.0
3.5
3.0
2.5
2.0
1.5
1.0
f2 (ppm)
10
20
30
40
50
60
70
80
90
100
110
120
130
f1 (ppm)

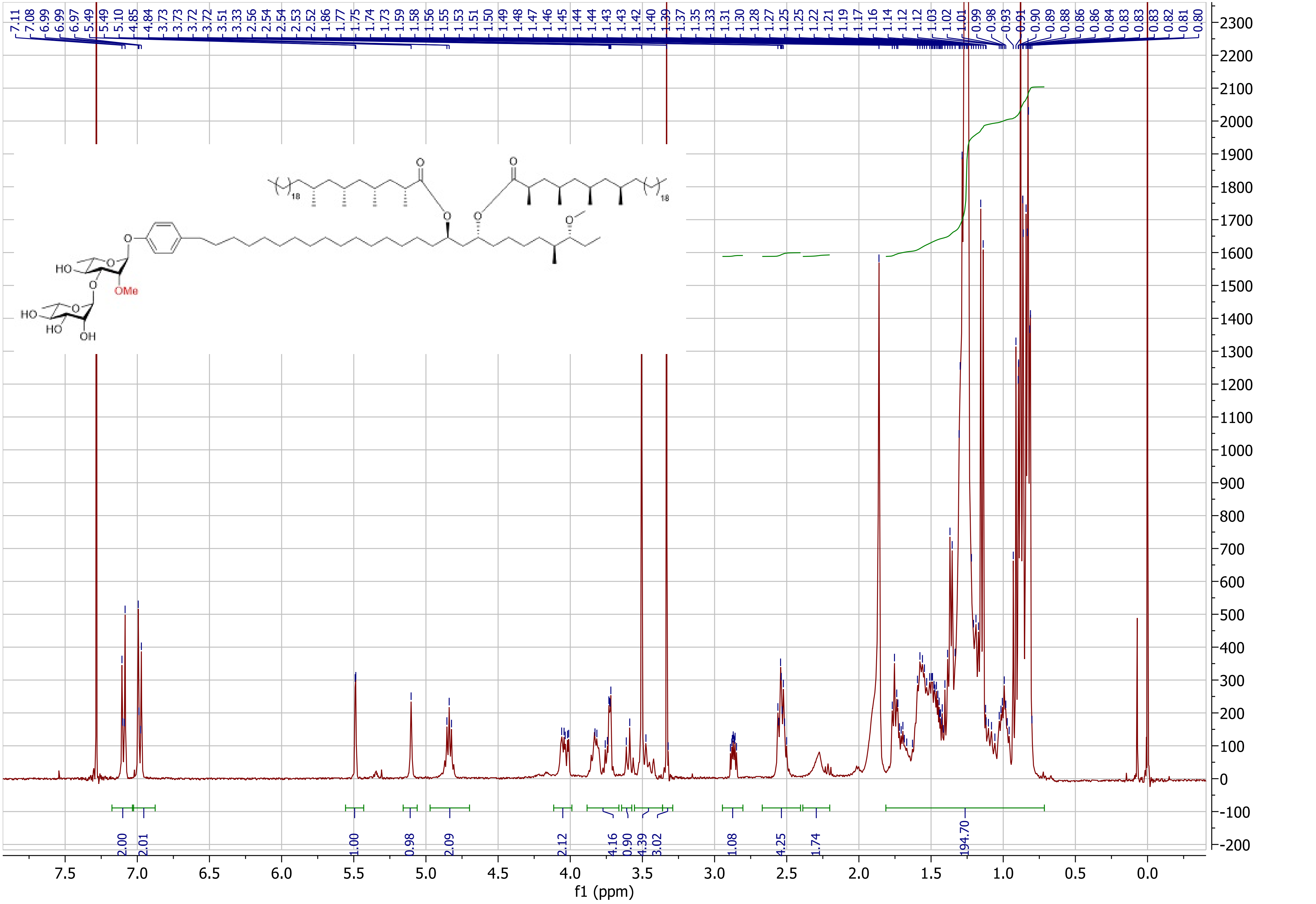

7.11 7.08 6.99 6.99 6.97 5.49 5.49 5.10 4.85 4.84 3.73 3.73 3.72 3.72 3.51 3.33 2.56 2.54 2.54 2.53 2.52 1.86 1.77 1.75 1.74 1.73 1.59 1.58 1.56 1.55 1.53 1.51 1.50 1.49 1.48 1.47 1.46 1.45 1.44 1.44 1.43 1.43 1.42 1.40 1.39 1.37 1.35 1.33 1.31 1.30 1.28 1.27 1.25 1.25 1.22 1.21 1.19 1.17 1.16 1.14 1.12 1.12 1.03 1.02 1.01 0.99 0.98 0.93 0.91 0.90 0.89 0.88 0.86 0.86 0.84 0.83 0.83 0.83 0.82 0.81 0.80
HO
OMe
HO
HO
OH
18
18
2.00
2.01
1.00
0.98
2.09
2.12
4.16
0.90
4.39
3.02
1.08
4.25
1.74
194.70
f1 (ppm)

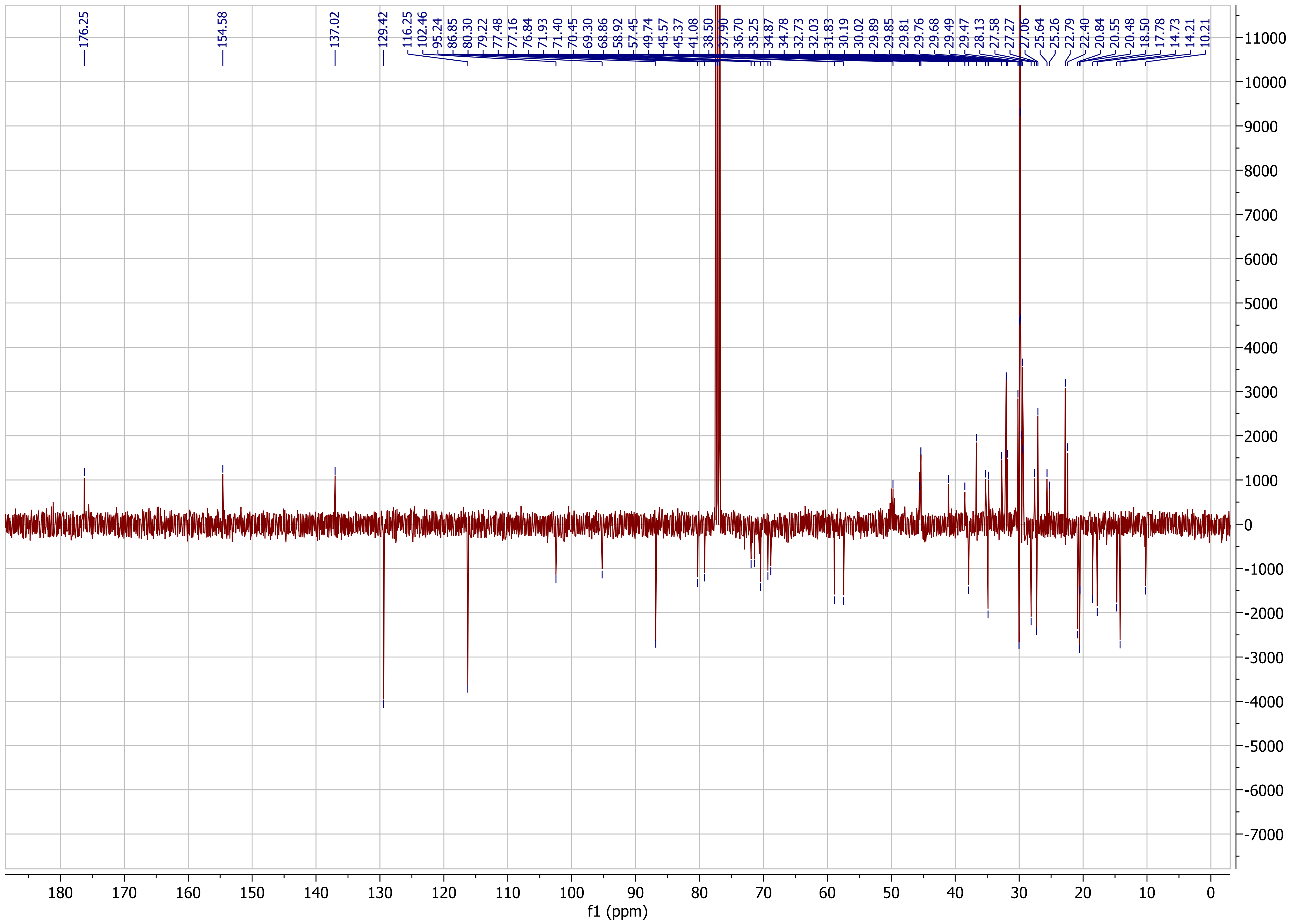
176.25
154.58
137.02
129.42
116.25
102.46
95.24
86.85
80.30
79.22
77.48
77.16
76.84
71.93
71.40
70.45
69.30
68.86
58.92
57.45
49.74
45.57
45.37
41.08
38.50
37.90
36.70
35.25
34.87
34.78
32.73
32.03
31.83
30.19
30.02
29.89
29.85
29.81
29.76
29.68
29.49
29.47
28.13
27.58
27.27
27.06
25.64
25.26
22.79
22.40
20.84
20.55
20.48
18.50
17.78
14.73
14.21
10.21
f1 (ppm)

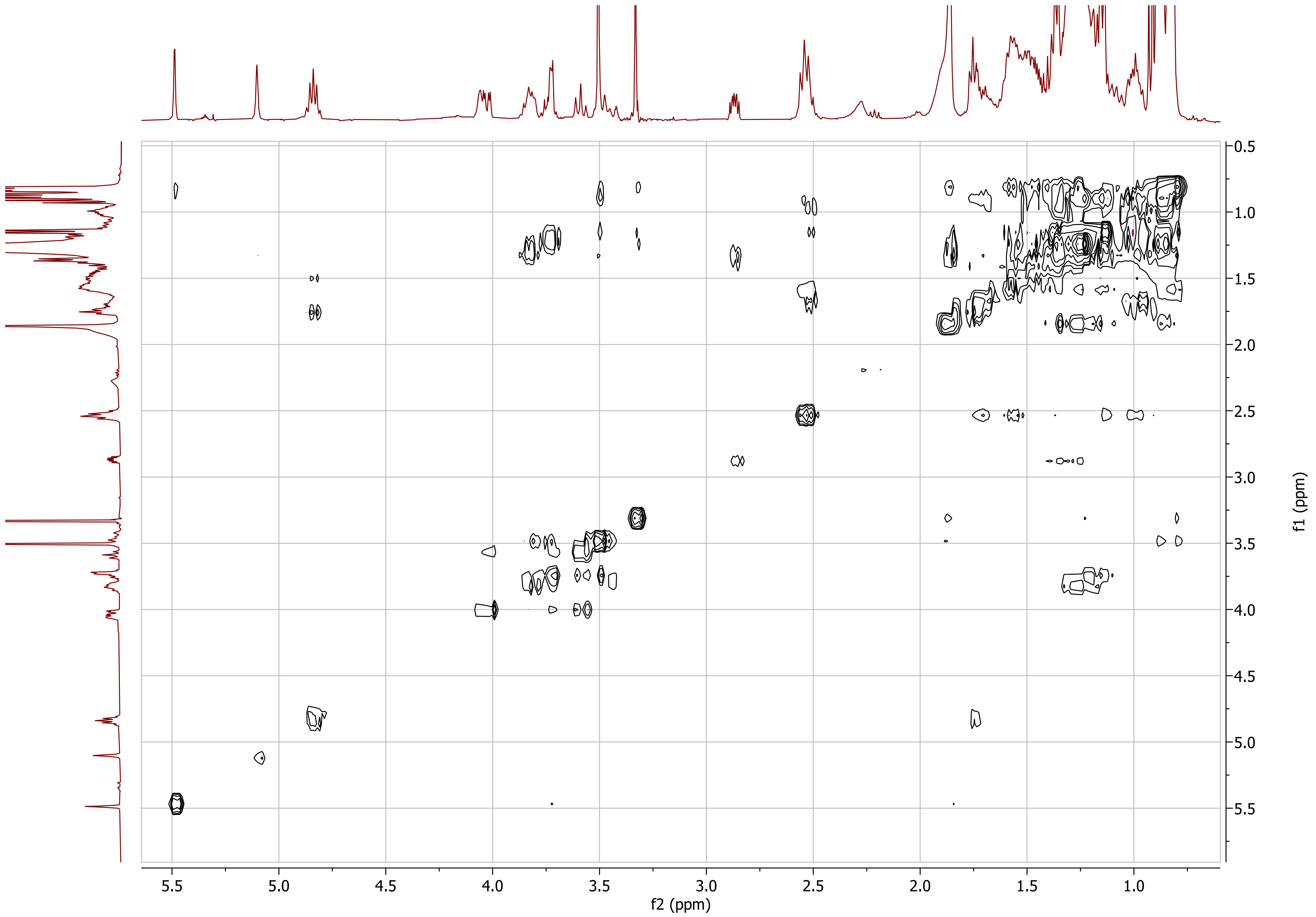

0.5
1.0
1.5
2.0
2.5
3.0
3.5
4.0
4.5
5.0
5.5
f1 (ppm)
5.5
5.0
4.5
4.0
3.5
3.0
2.5
2.0
1.5
1.0
f2 (ppm)

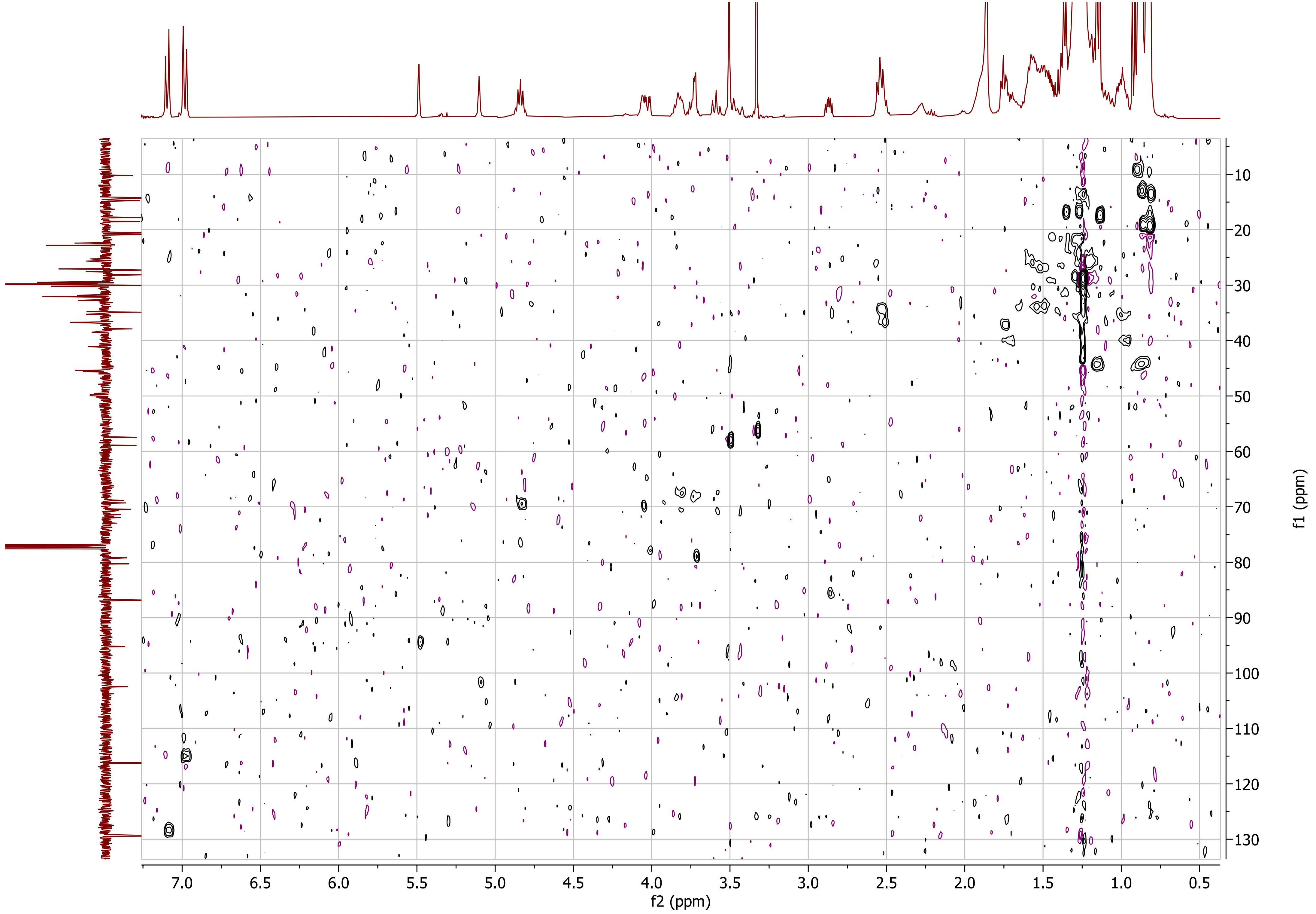

f2 (ppm)
f1 (ppm)

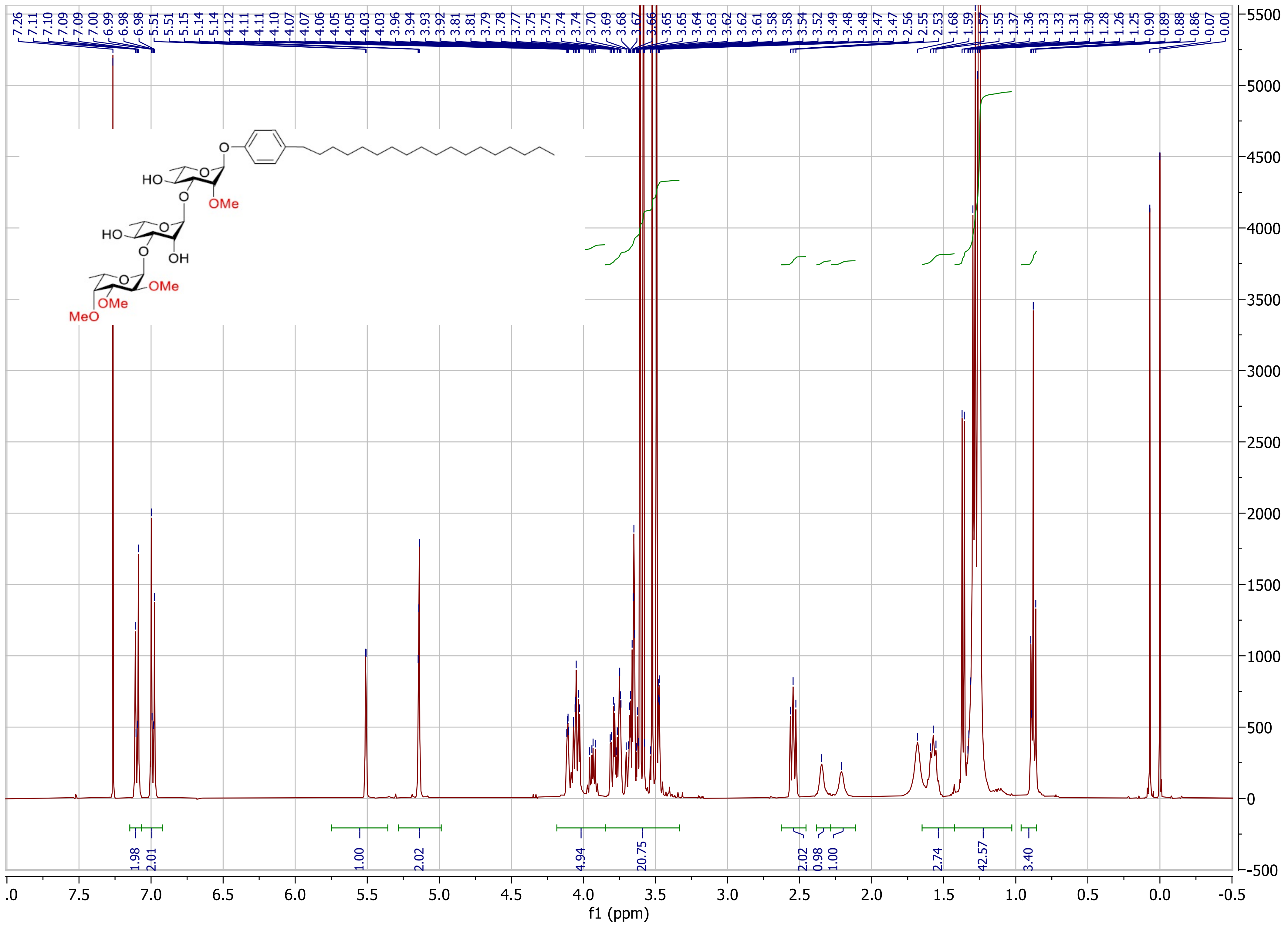

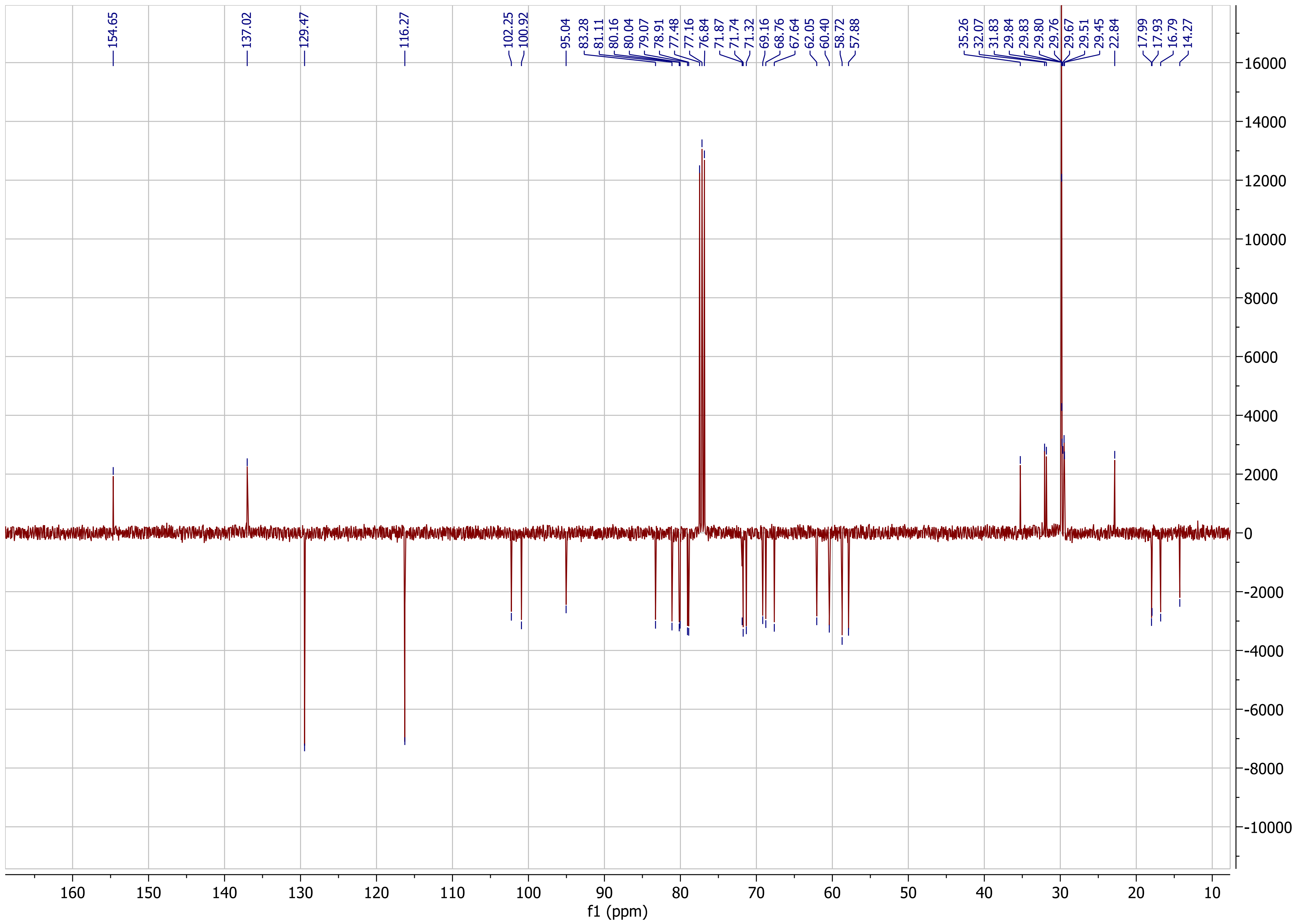
154.65
137.02
129.47
116.27
102.25
100.92
95.04
83.28
81.11
80.16
80.04
79.07
78.91
77.48
77.16
76.84
71.87
71.74
71.32
69.16
68.76
67.64
62.05
60.40
58.72
57.88
35.26
32.07
31.83
29.84
29.83
29.80
29.76
29.67
29.51
29.45
22.84
17.99
17.93
16.79
14.27
f1 (ppm)

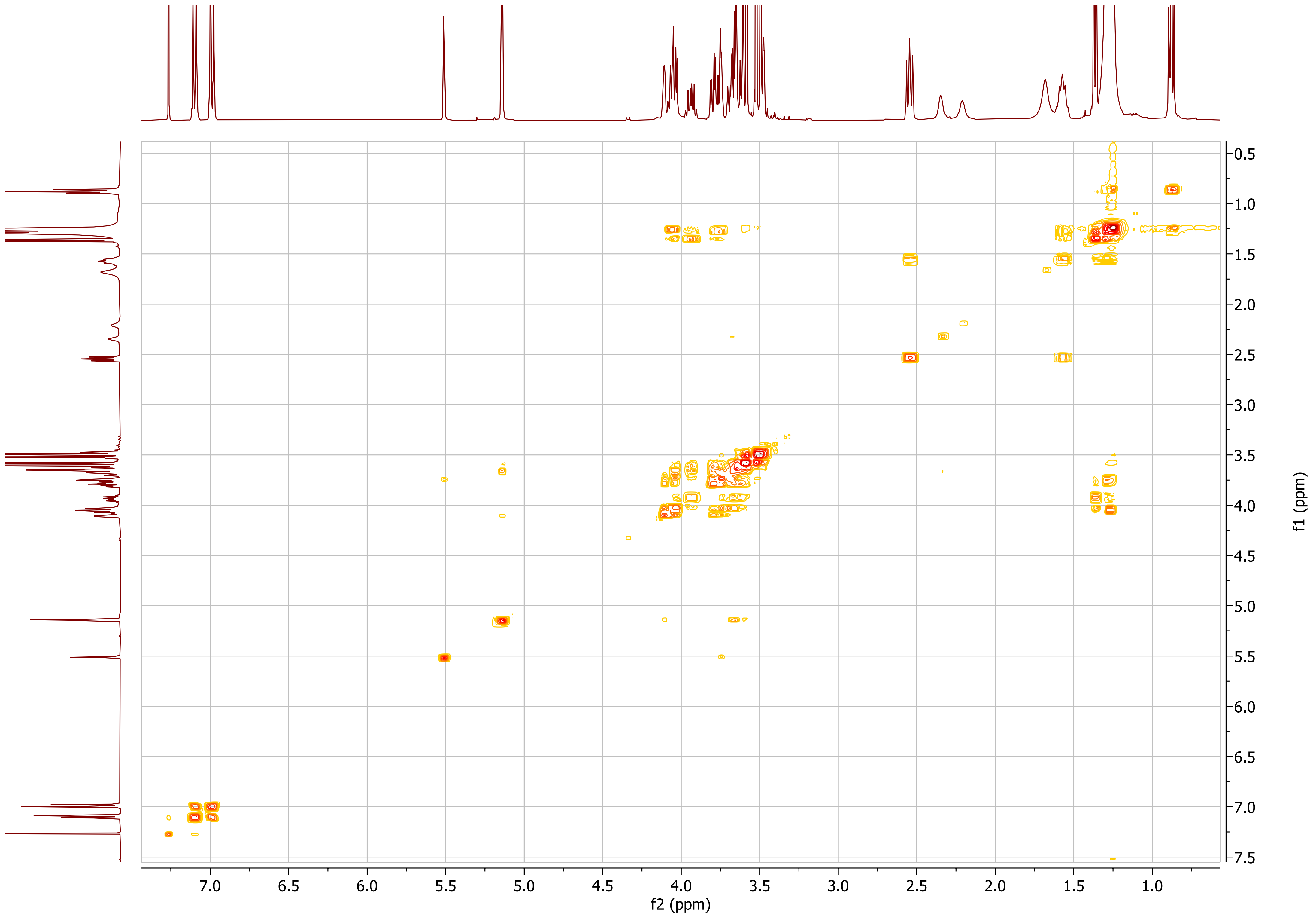

f2 (ppm)
f1 (ppm)
7.0
6.5
6.0
5.5
5.0
4.5
4.0
3.5
3.0
2.5
2.0
1.5
1.0
0.5
7.5

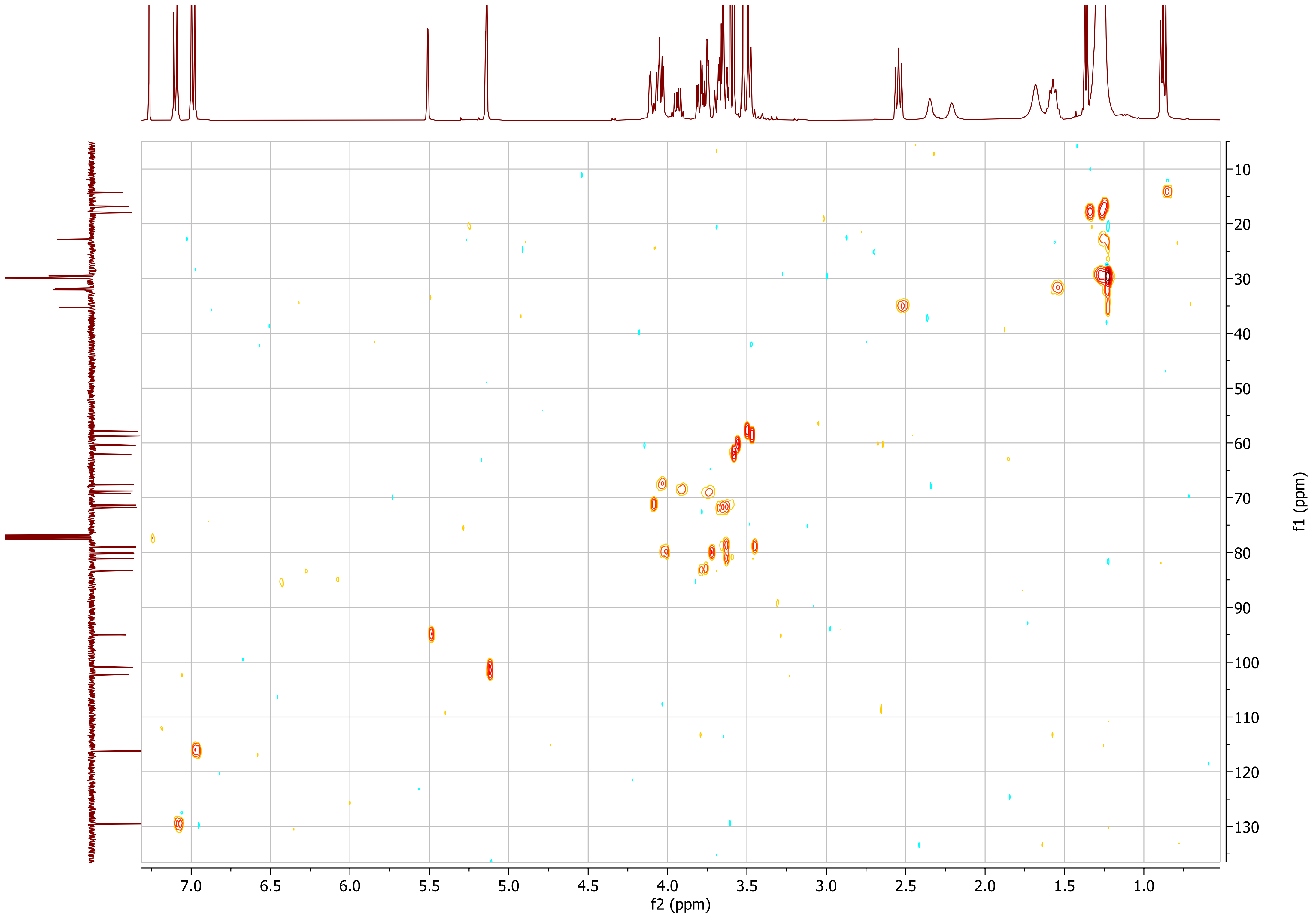
f2 (ppm)
f1 (ppm)

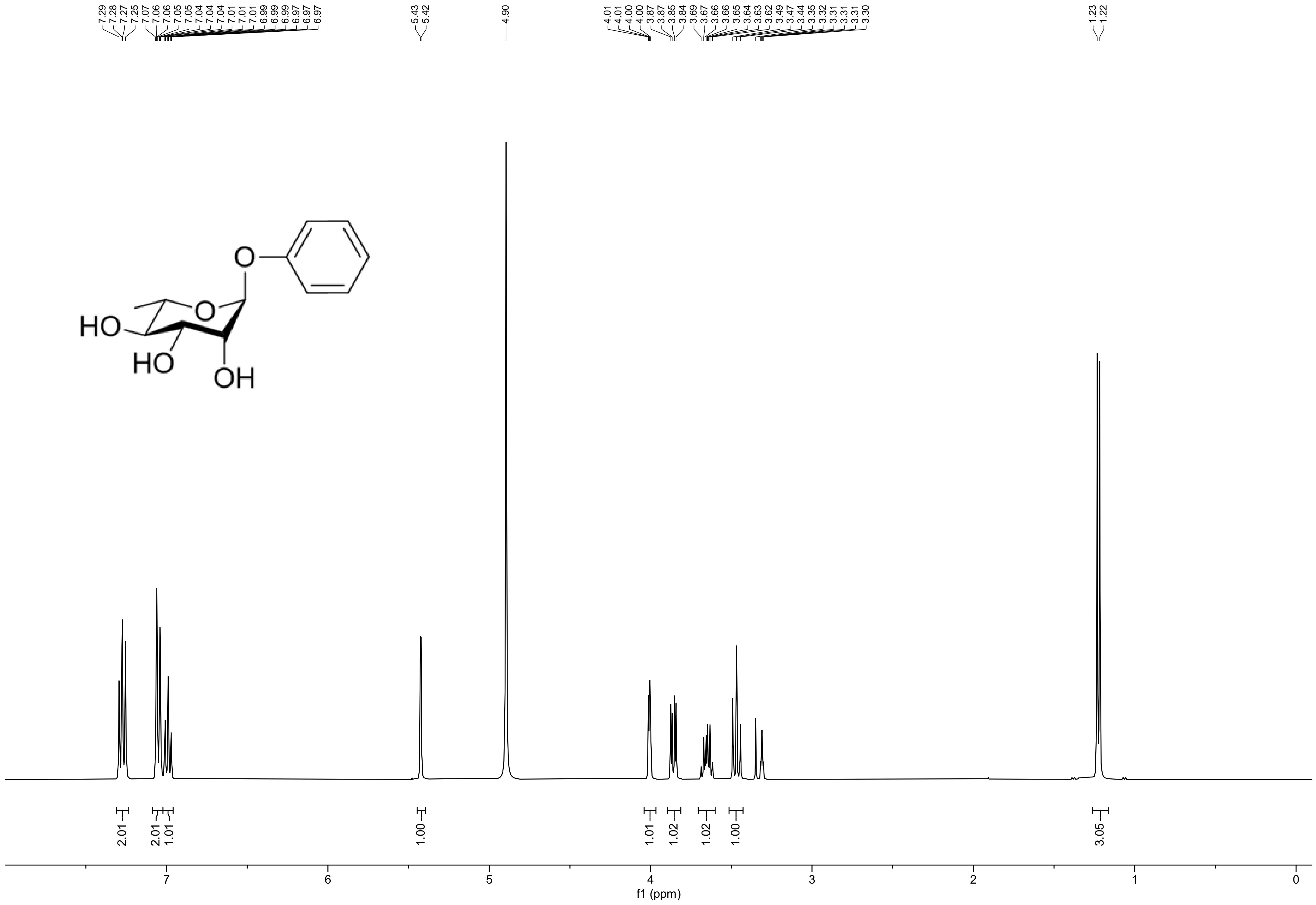
7.29
7.28
7.27
7.25
7.07
7.06
7.06
7.05
7.04
7.04
7.01
7.01
6.99
6.99
6.97
6.97
5.43
5.42
4.90
4.01
4.01
4.00
4.00
3.87
3.87
3.85
3.84
3.69
3.67
3.66
3.66
3.65
3.64
3.63
3.62
3.49
3.47
3.44
3.35
3.32
3.31
3.31
3.31
3.30
1.23
1.22
HO
HO
OH
O
O
2.01
2.01
1.01
1.00
1.01
1.02
1.02
1.00
3.05
7
6
5
4
3
2
1
0
f1 (ppm)

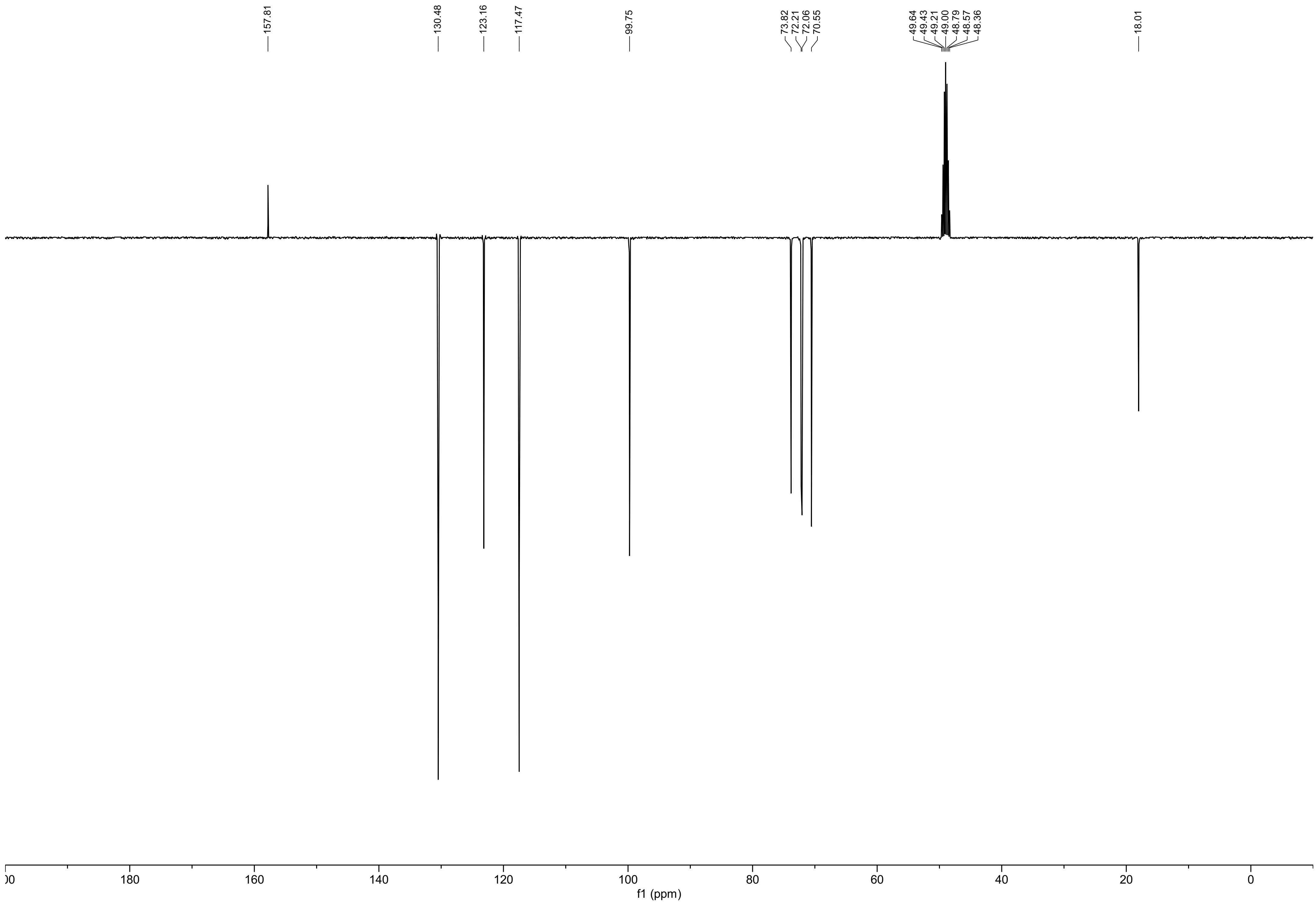
157.81
130.48
123.16
117.47
99.75
73.82
72.21
72.06
70.55
49.64
49.43
49.21
49.00
48.79
48.57
48.36
18.01
180
160
140
120
100
80
60
40
20
0
f1 (ppm)

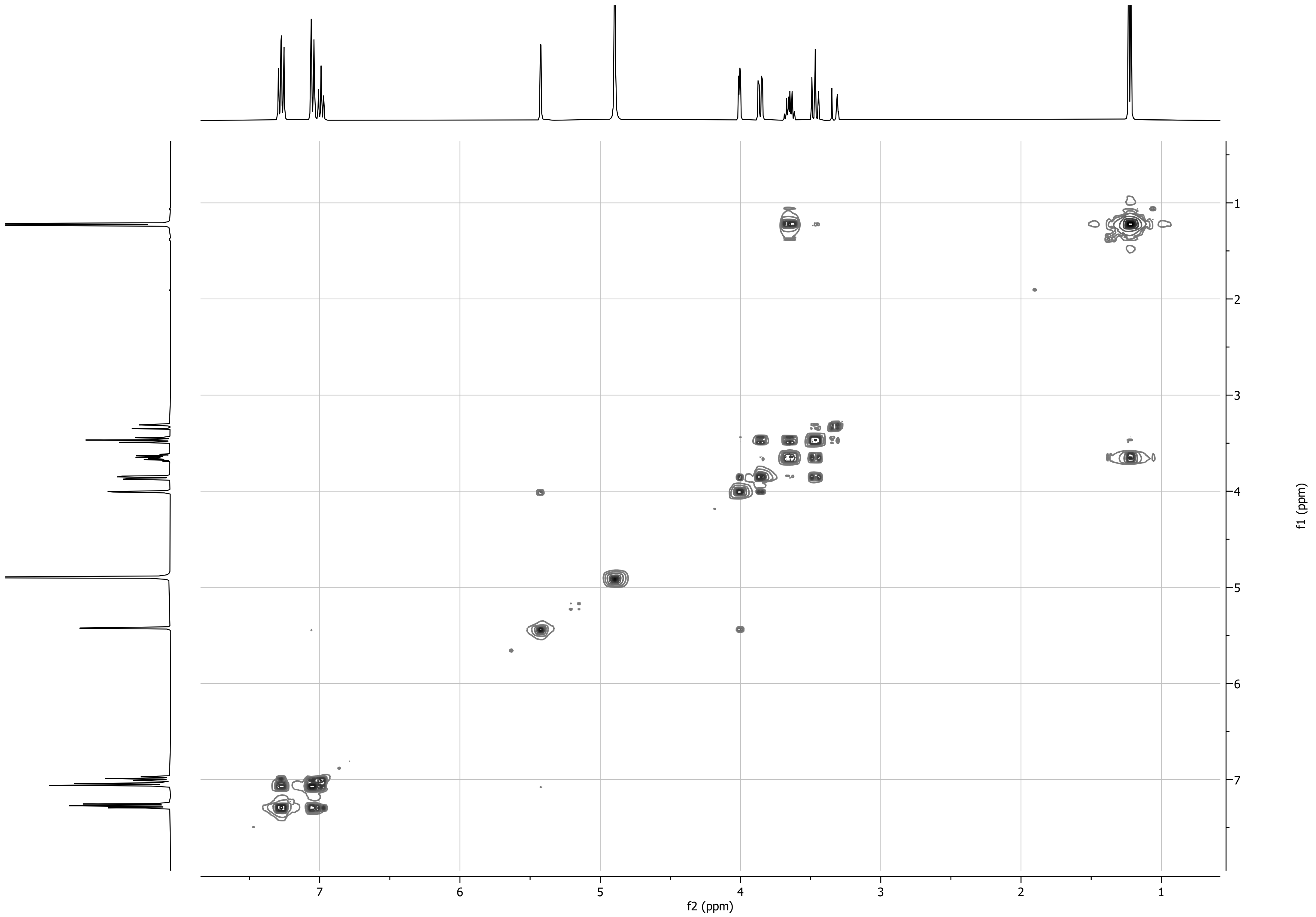

f2 (ppm)
f1 (ppm)

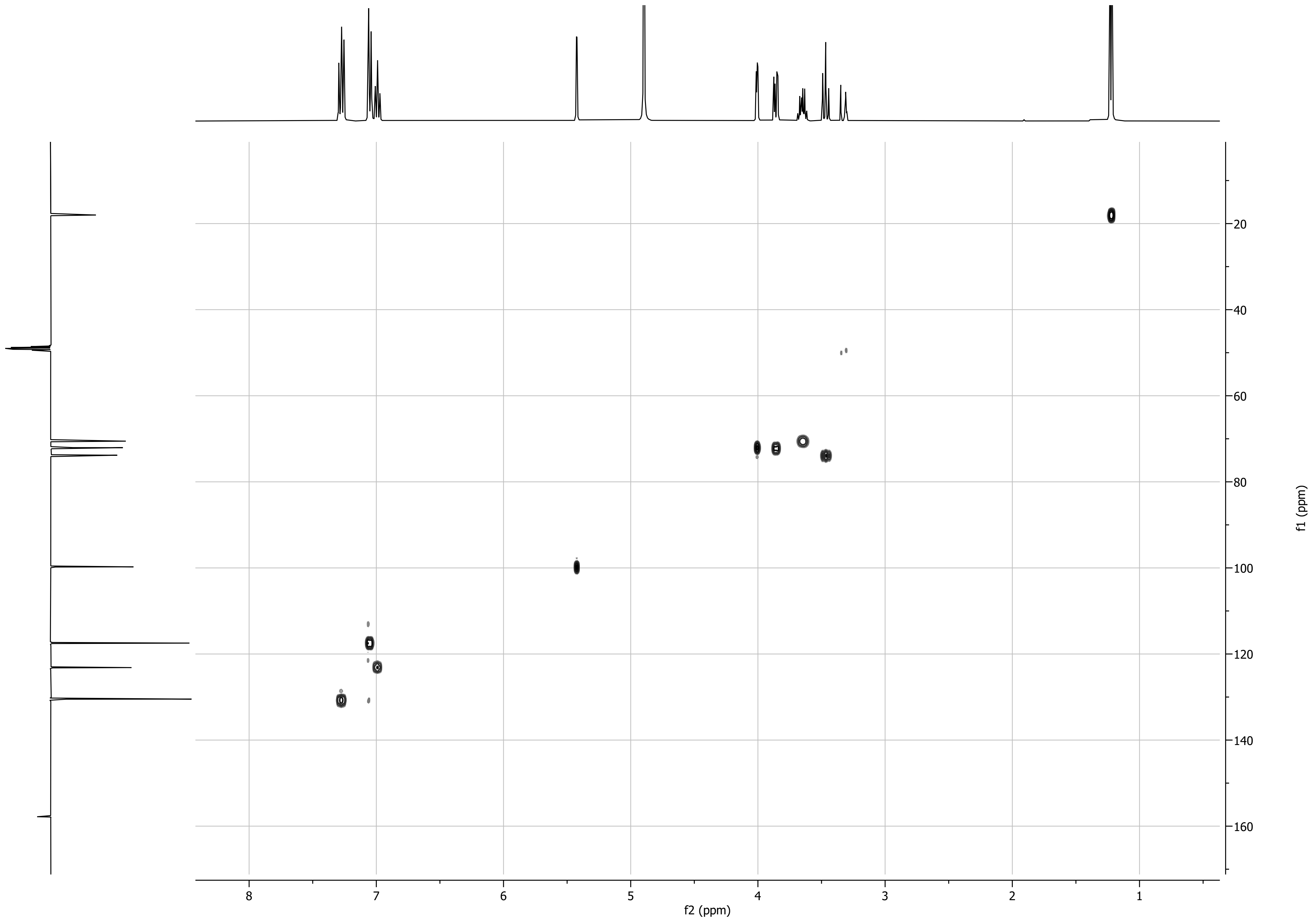

f1 (ppm)
f2 (ppm)

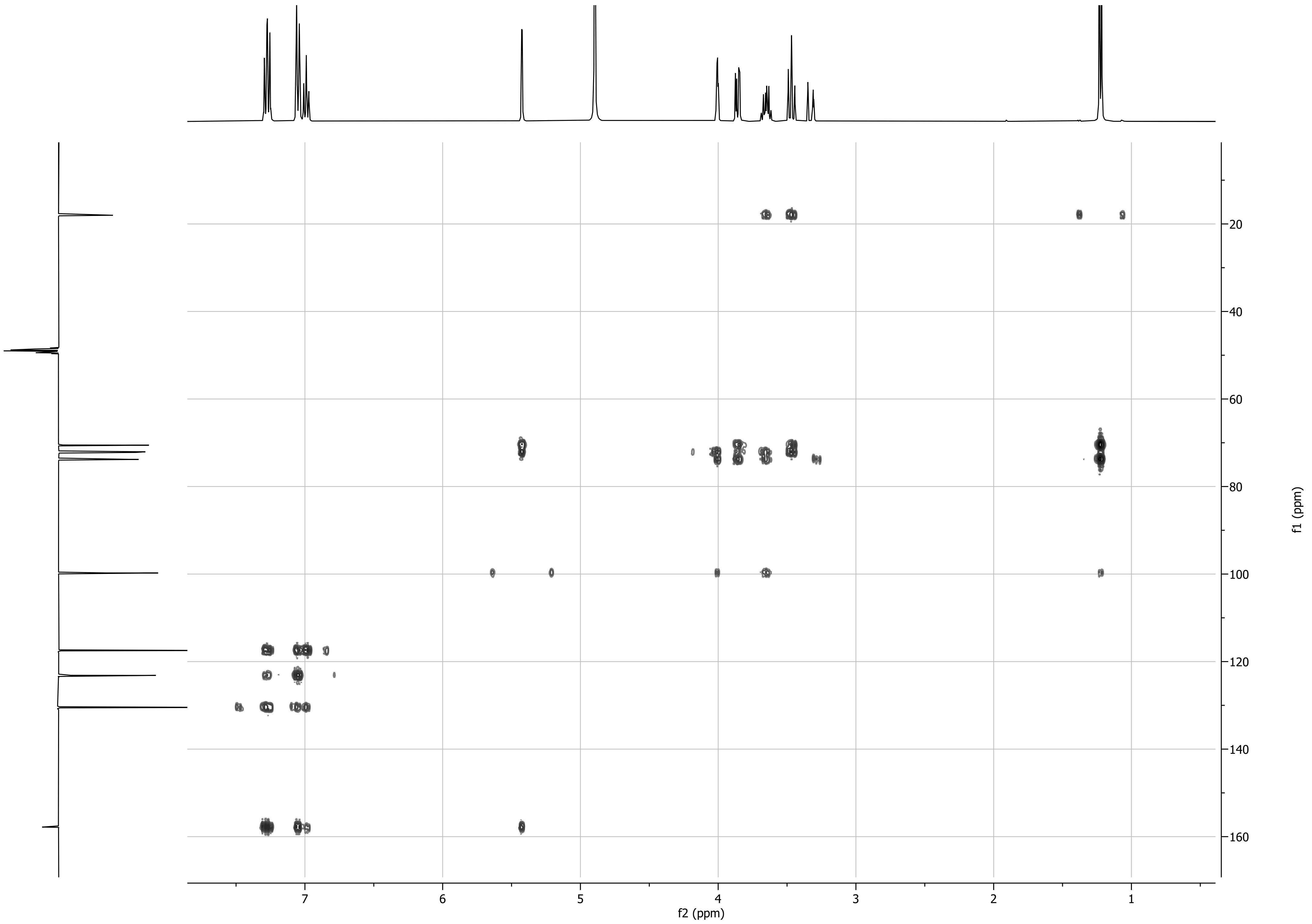

f2 (ppm)
f1 (ppm)
7
6
5
4
3
2
1
20
40
60
80
100
120
140
160

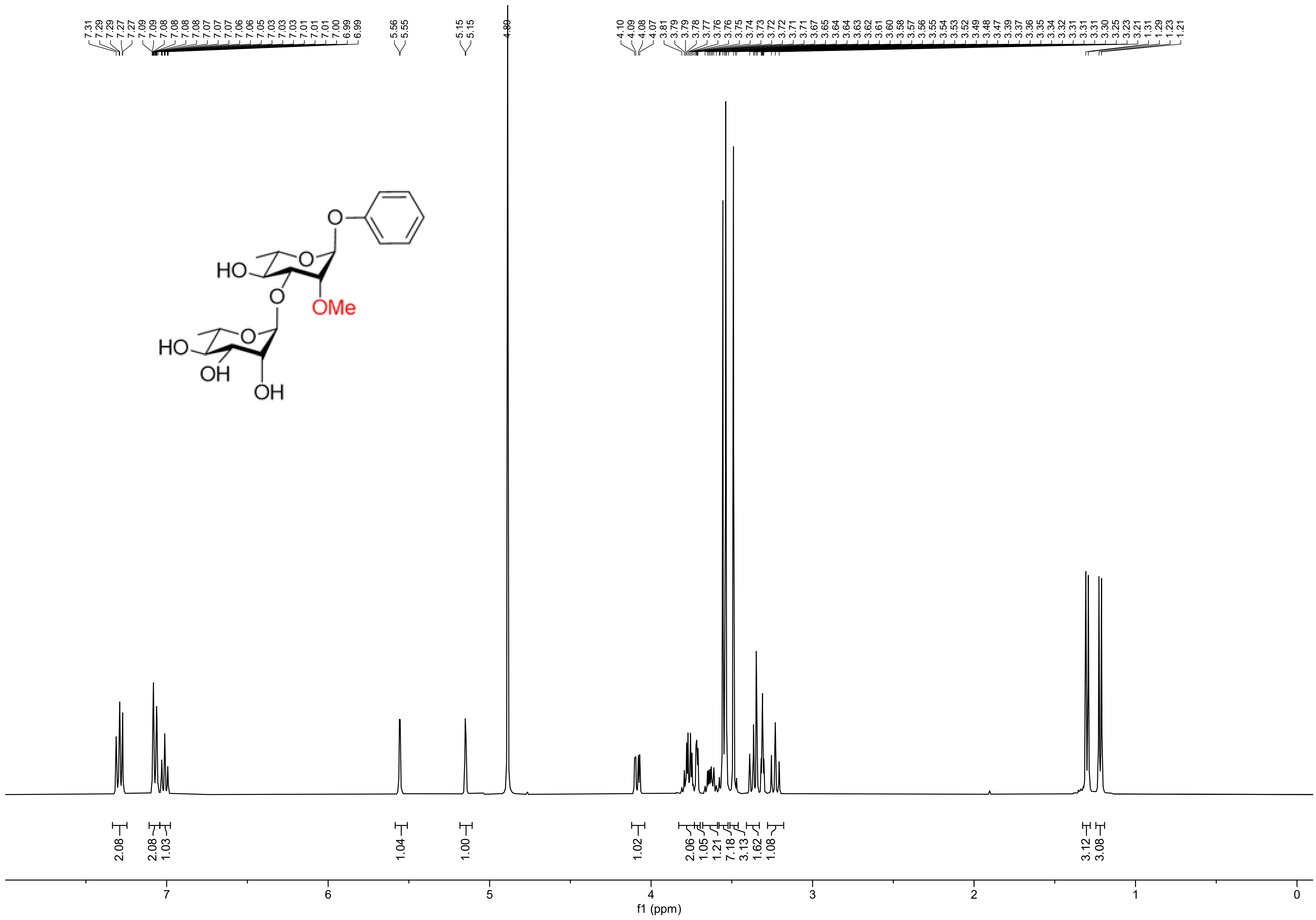
7.31 7.29 7.29 7.27 7.27 7.09 7.09 7.08 7.08 7.08 7.07 7.07 7.06 7.06 7.05 7.03 7.03 7.03 7.01 7.01 7.01 7.00 6.99 6.99
5.56 5.55
5.15 5.15
4.89
4.10 4.09 4.08 4.07 3.81 3.79 3.78 3.77 3.76 3.76 3.75 3.74 3.73 3.72 3.72 3.71 3.71 3.67 3.65 3.64 3.64 3.63 3.62 3.61 3.60 3.58 3.57 3.56 3.55 3.54 3.53 3.52 3.49 3.48 3.47 3.39 3.37 3.36 3.35 3.34 3.32 3.31 3.31 3.31 3.30 3.25 3.23 3.21 1.31 1.29 1.23 1.21
HO
O
O
OMe
HO
OH
OH
2.08
2.08
1.03
1.04
1.00
1.02
2.06
1.05
1.21
7.18
3.13
1.62
1.08
3.12
3.08
7
6
5
4
3
2
1
0
f1 (ppm)

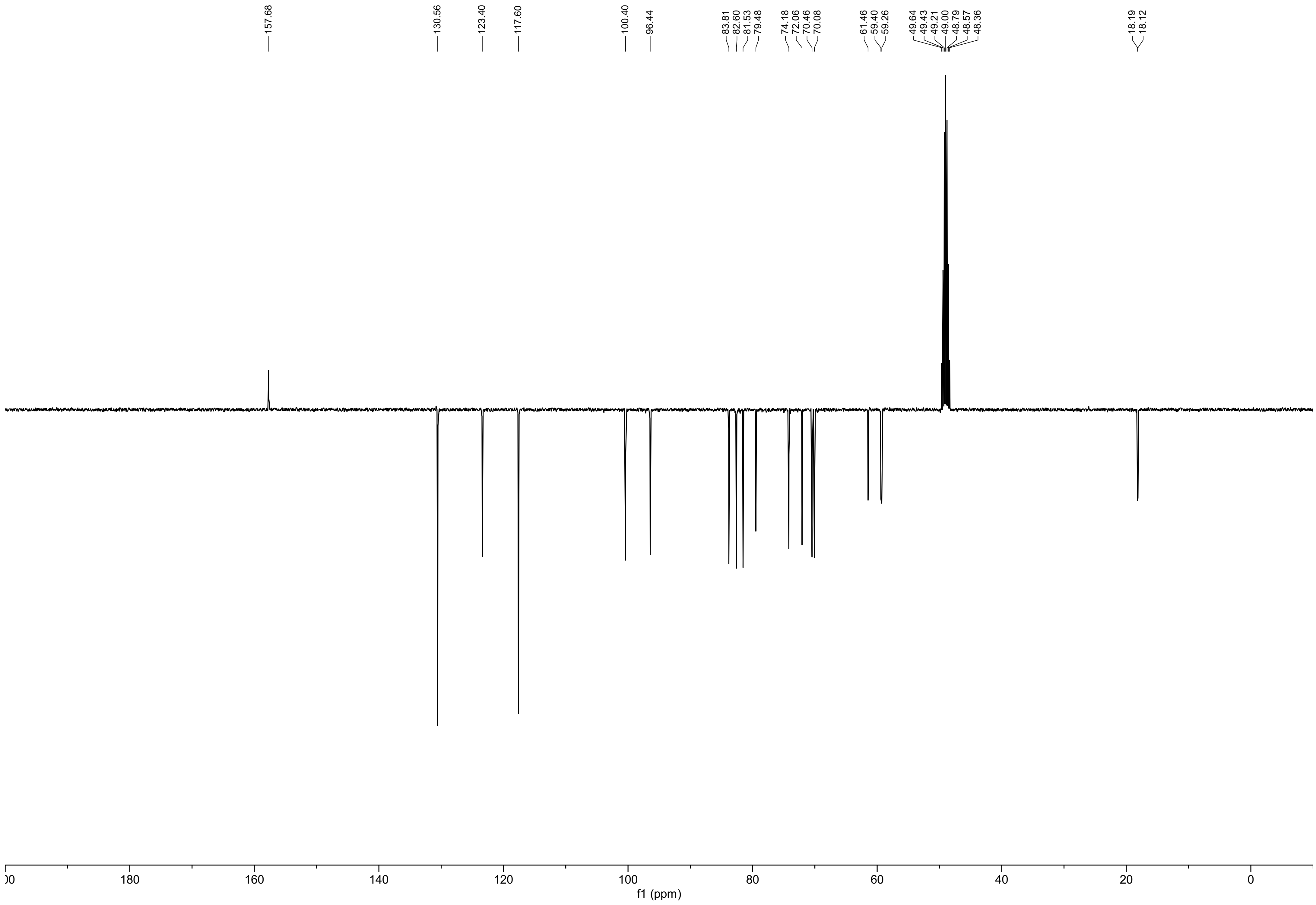

157.68
130.56
123.40
117.60
100.40
96.44
83.81
82.60
81.53
79.48
74.18
72.06
70.46
70.08
61.46
59.40
59.26
49.64
49.43
49.21
49.00
48.79
48.57
48.36
18.19
18.12
180
160
140
120
100
80
60
40
20
0
f1 (ppm)

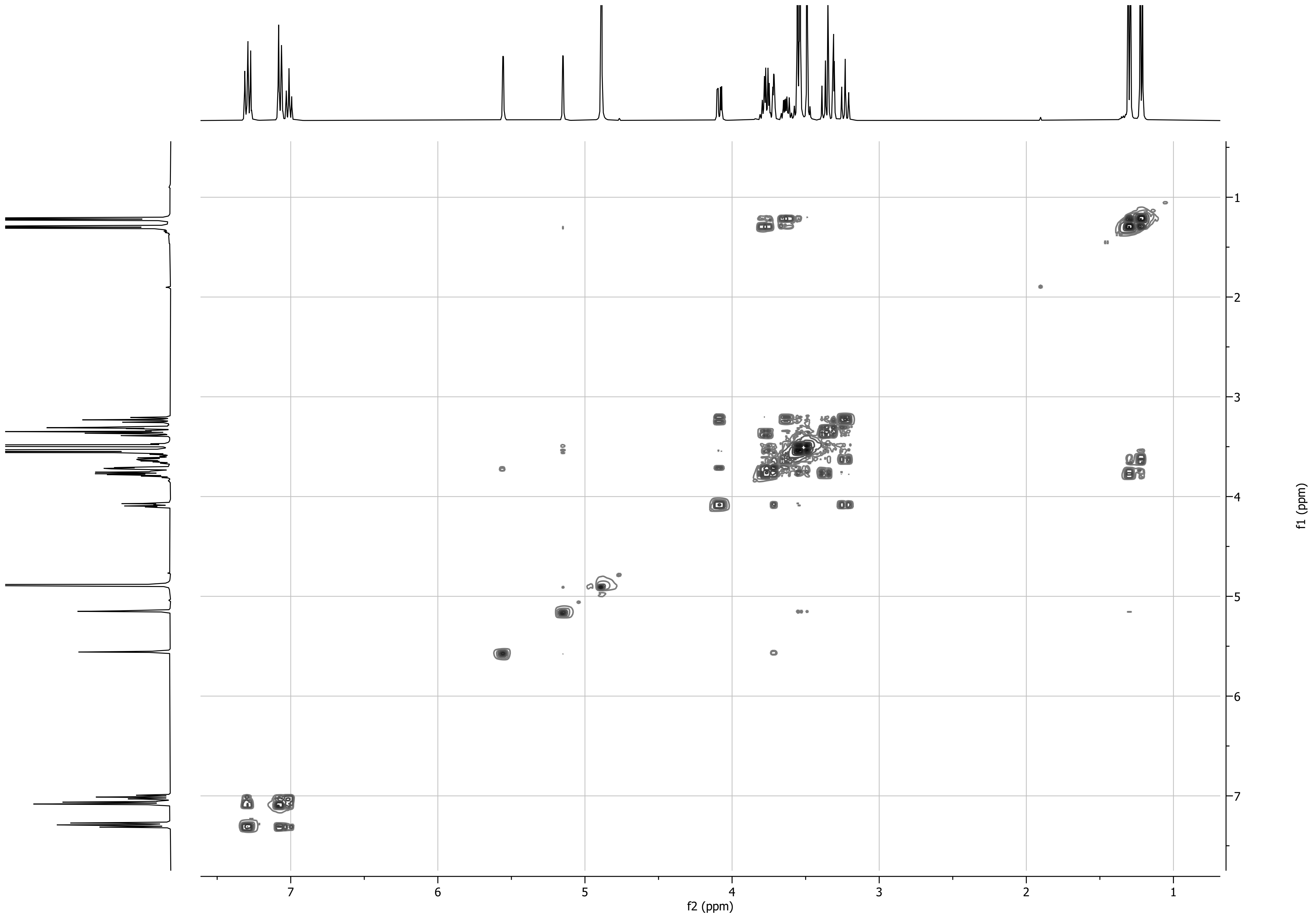

f2 (ppm)
f1 (ppm)
7
6
5
4
3
2
1

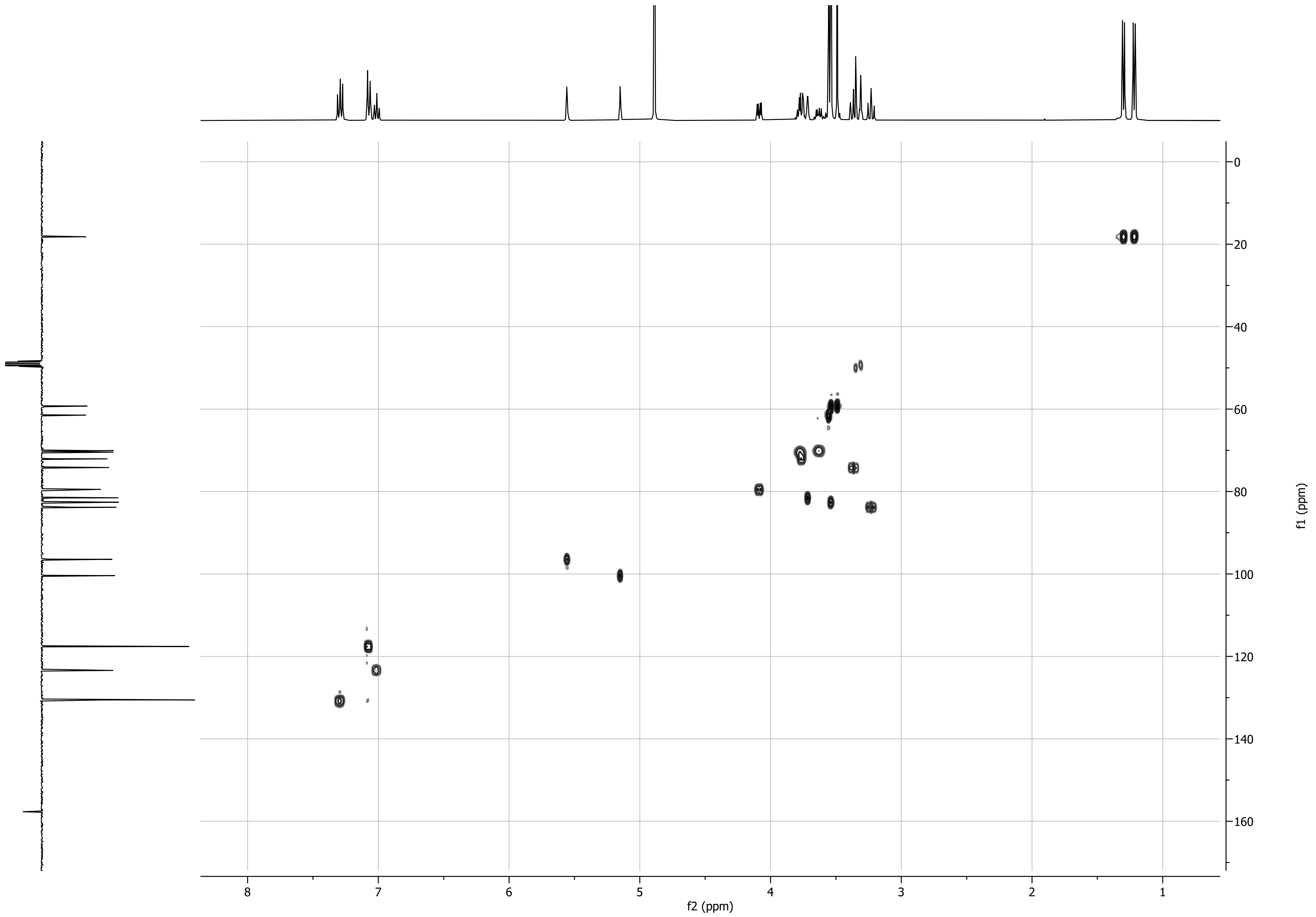

f1 (ppm)
f2 (ppm)

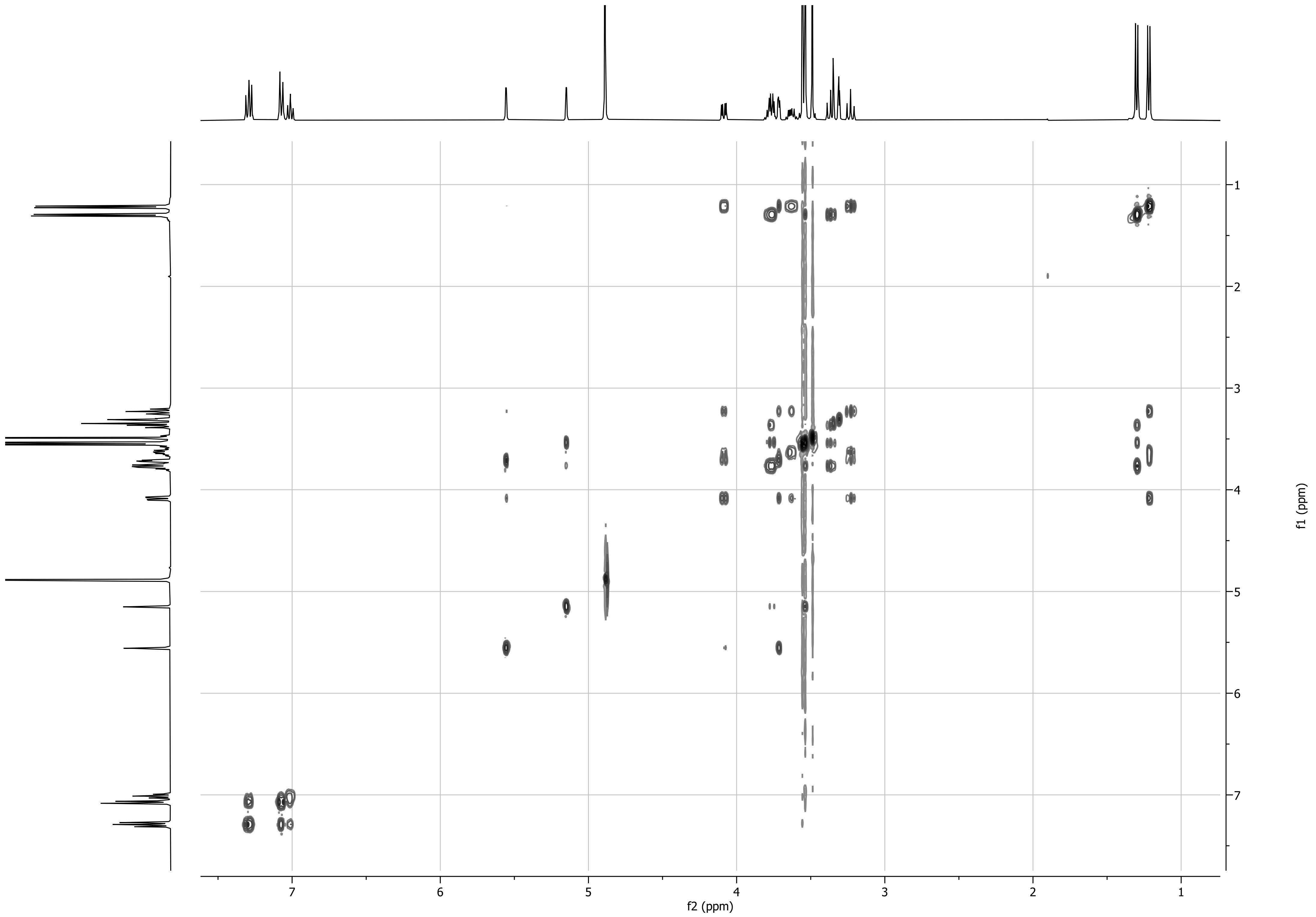
f2 (ppm)
f1 (ppm)
7
6
5
4
3
2
1

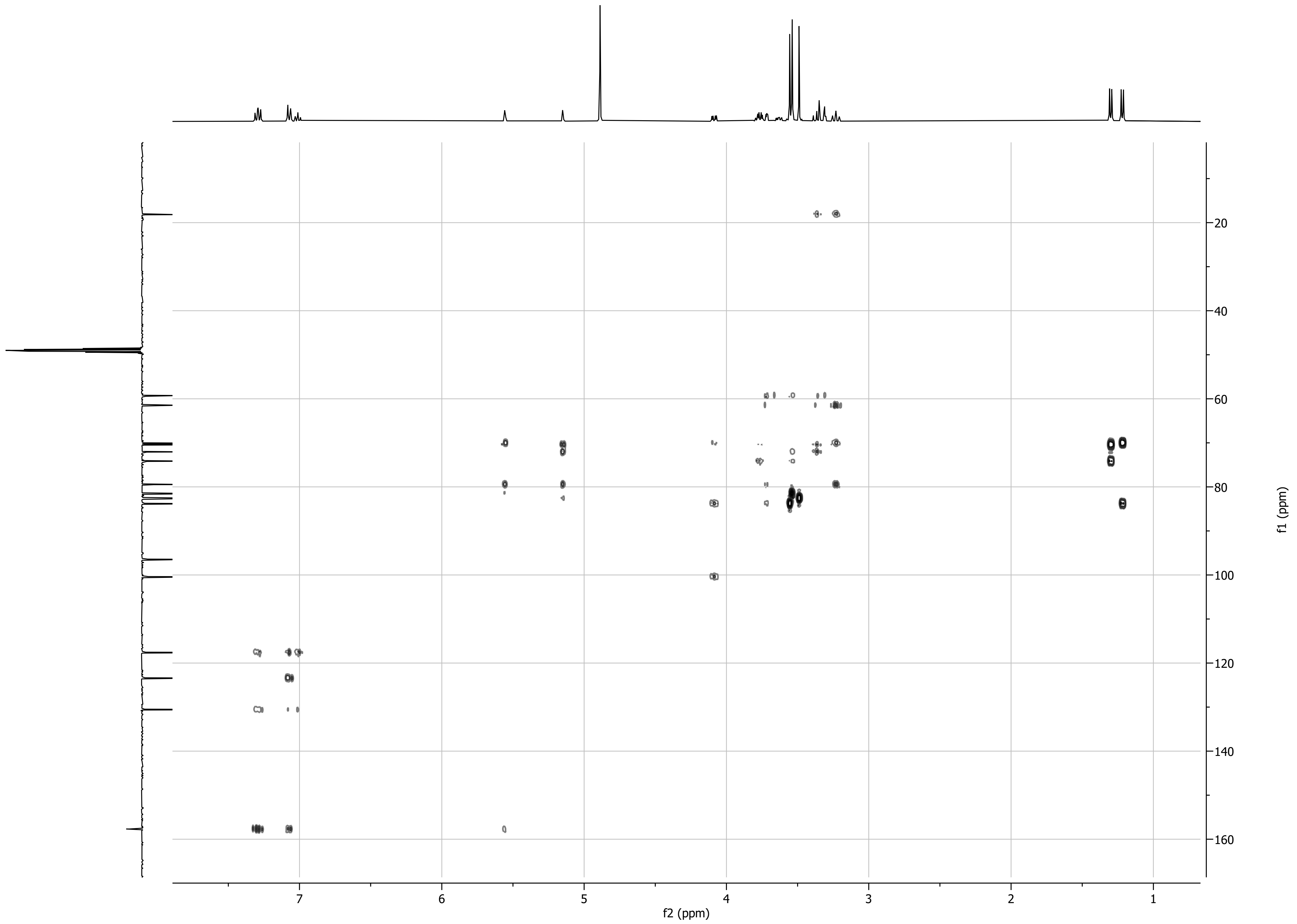

f2 (ppm)
f1 (ppm)
7
6
5
4
3
2
1
20
40
60
80
100
120
140
160

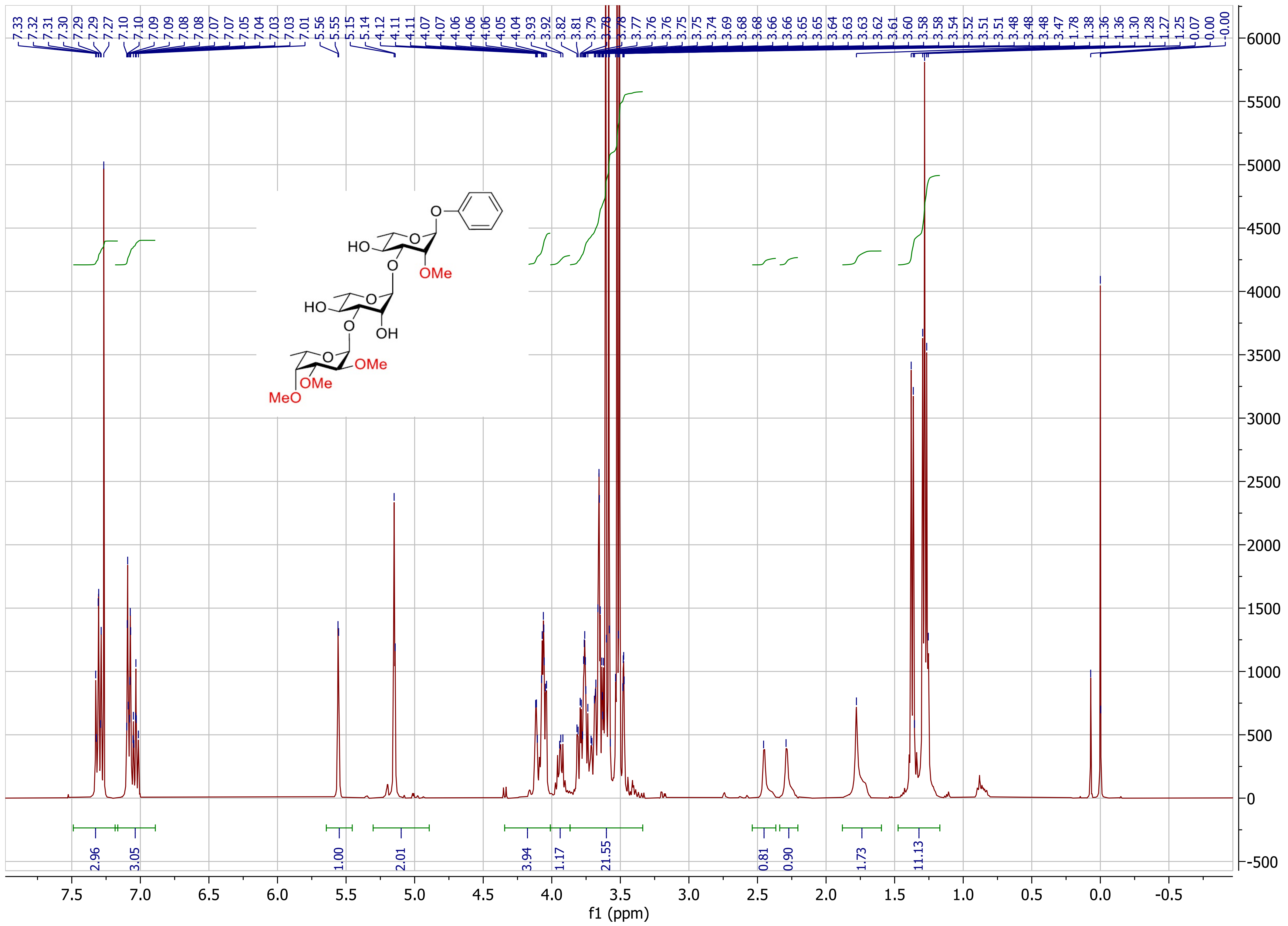

HO
O
O
OMe
HO
O
O
OH
O
OMe
OMe
MeO
f1 (ppm)

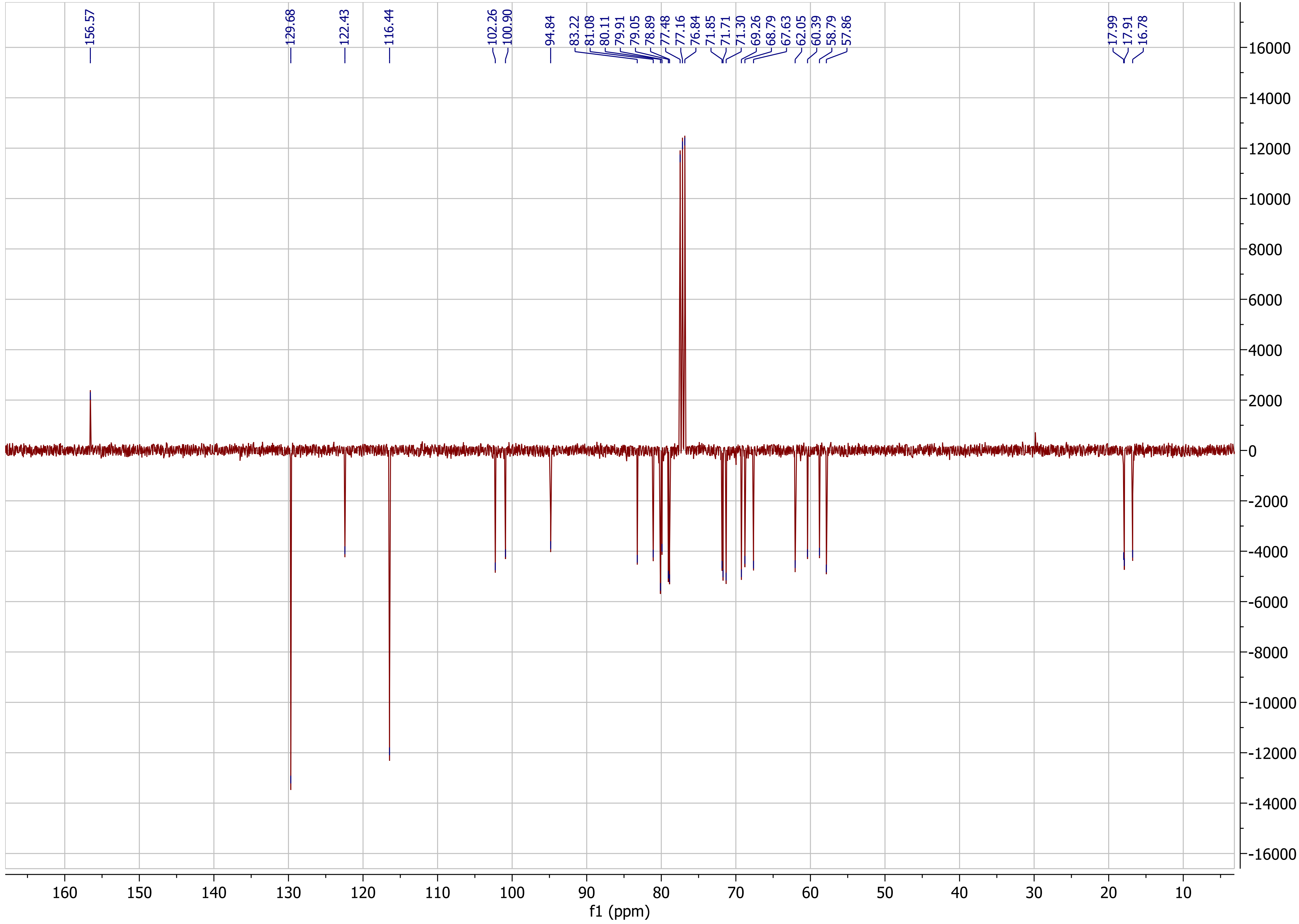

156.57
129.68
122.43
116.44
102.26
100.90
94.84
83.22
81.08
80.11
79.91
79.05
78.89
77.48
77.16
76.84
71.85
71.71
71.30
69.26
68.79
67.63
62.05
60.39
58.79
57.86
17.99
17.91
16.78
f1 (ppm)

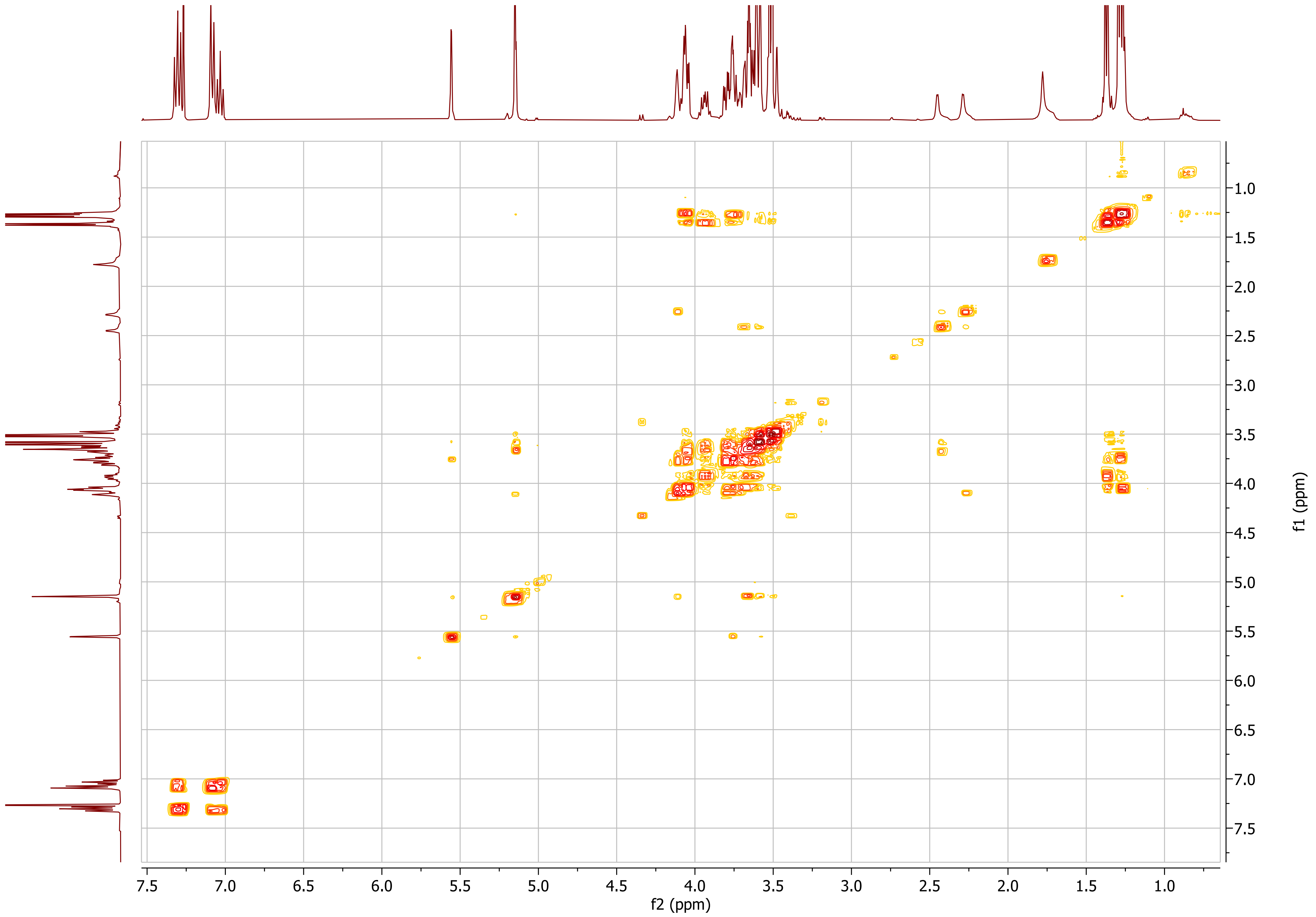
7.5
7.0
6.5
6.0
5.5
5.0
4.5
4.0
3.5
3.0
2.5
2.0
1.5
1.0
f2 (ppm)
f1 (ppm)

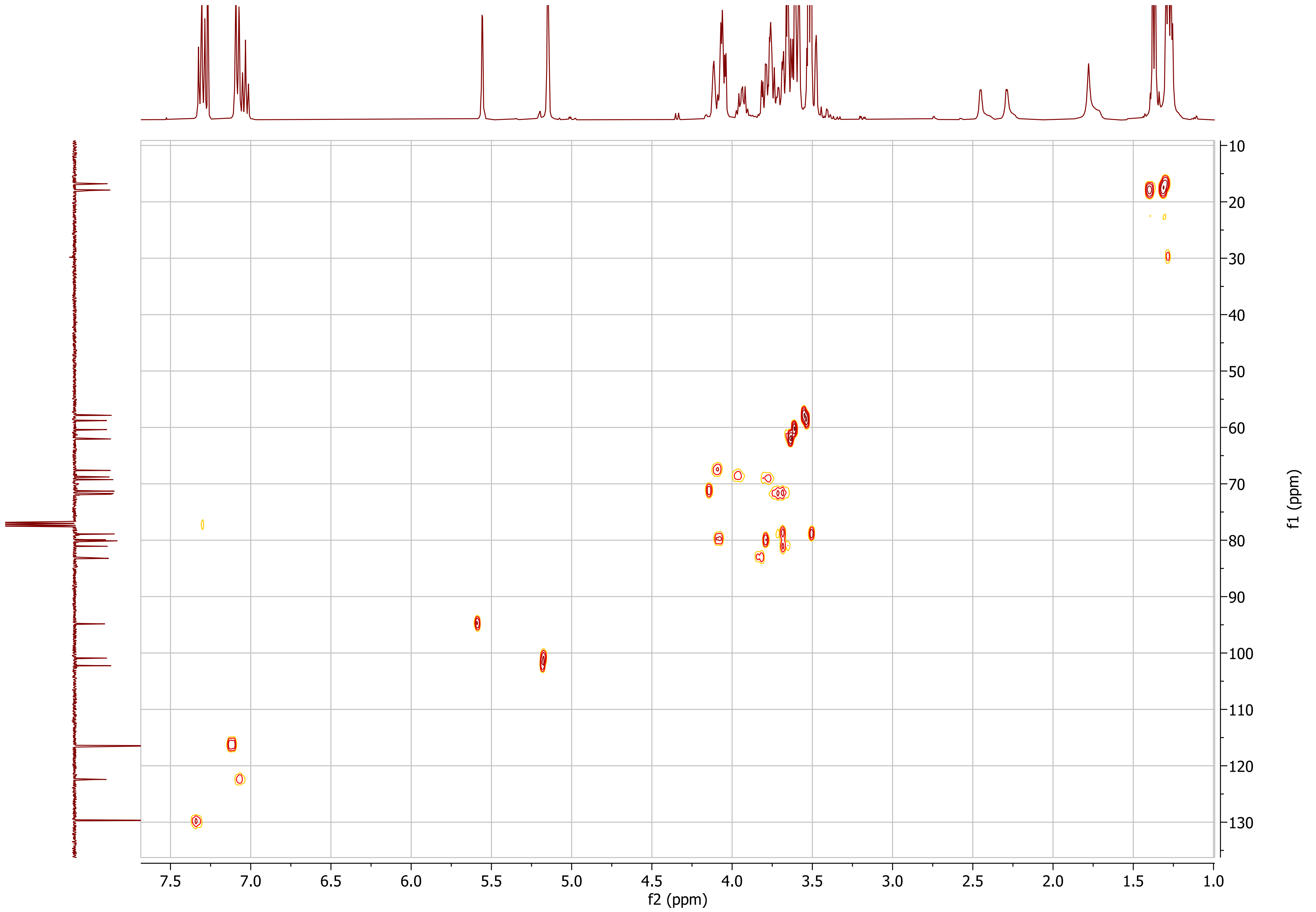

7.5
7.0
6.5
6.0
5.5
5.0
4.5
4.0
3.5
3.0
2.5
2.0
1.5
1.0
f2 (ppm)
10
20
30
40
50
60
70
80
90
100
110
120
130
f1 (ppm)

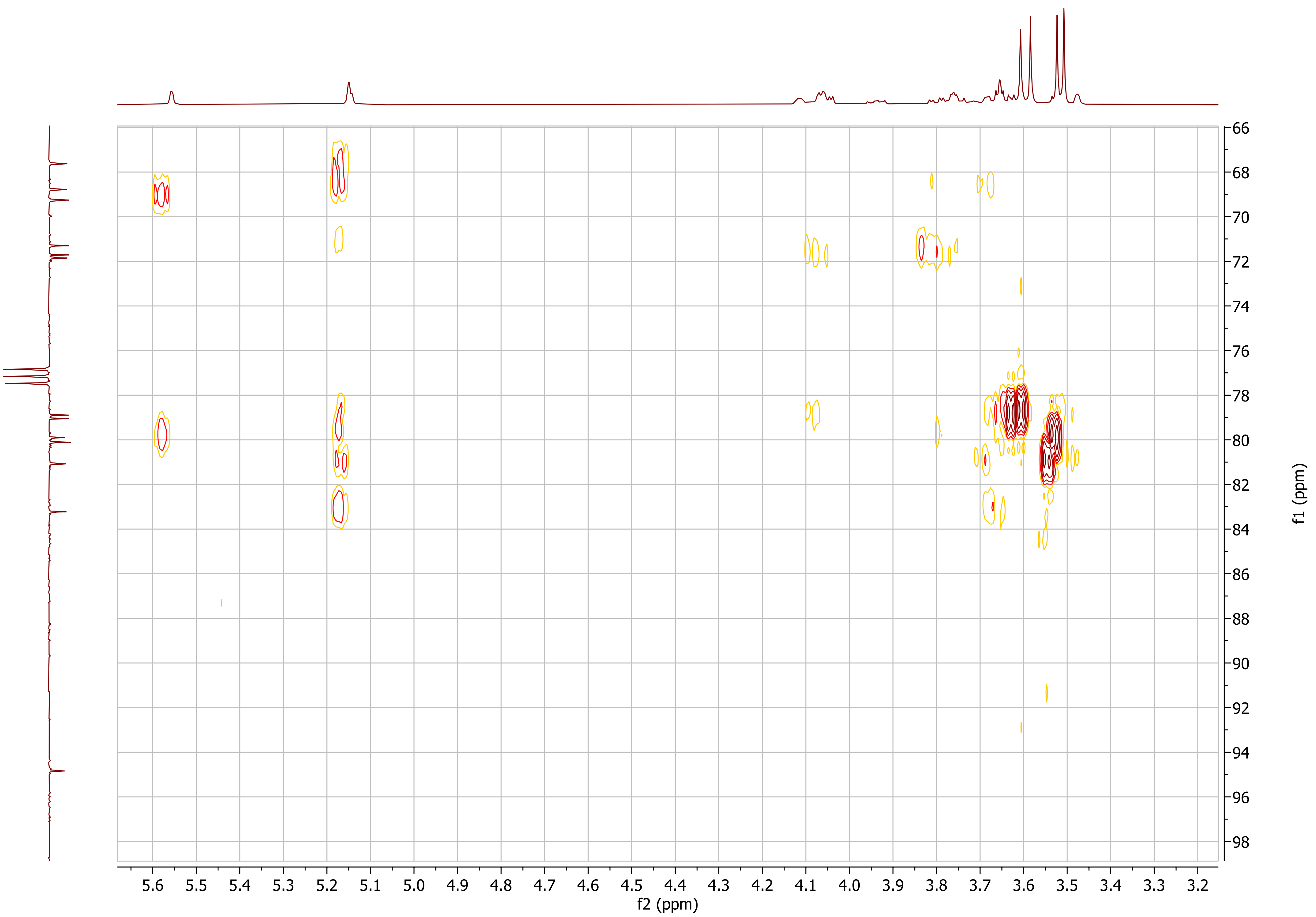

5.6
5.5
5.4
5.3
5.2
5.1
5.0
4.9
4.8
4.7
4.6
4.5
4.4
4.3
4.2
4.1
4.0
3.9
3.8
3.7
3.6
3.5
3.4
3.3
3.2
f2 (ppm)
66
68
70
72
74
76
78
80
82
84
86
88
90
92
94
96
98
f1 (ppm)

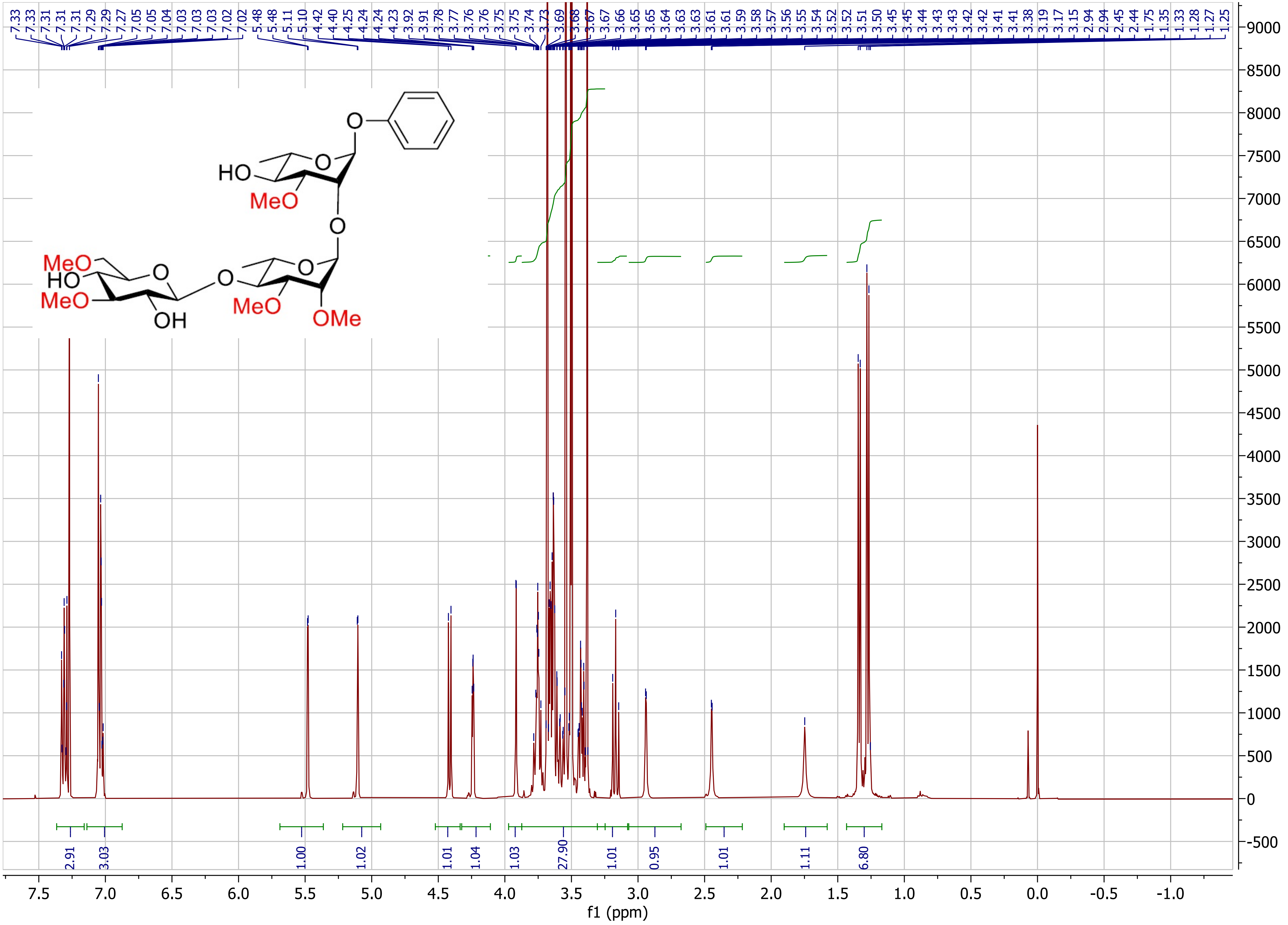

7.33 7.33 7.31 7.31 7.29 7.29 7.27 7.05 7.05 7.04 7.03 7.03 7.03 7.02 7.02 5.48 5.48 5.11 5.10 4.42 4.40 4.25 4.24 4.24 4.23 3.92 3.91 3.78 3.77 3.76 3.76 3.75 3.75 3.74 3.73 3.69 3.68 3.67 3.67 3.66 3.65 3.65 3.64 3.63 3.63 3.61 3.61 3.59 3.58 3.57 3.56 3.55 3.54 3.52 3.52 3.51 3.50 3.45 3.45 3.44 3.43 3.43 3.42 3.42 3.41 3.41 3.38 3.19 3.17 3.15 2.94 2.94 2.45 2.44 1.75 1.35 1.33 1.28 1.27 1.25
HO
MeO
O
MeO
HO
MeO
OH
MeO
OMe
2.91 3.03 1.00 1.02 1.01 1.04 1.03 27.90 1.01 0.95 1.01 1.11 6.80
f1 (ppm)

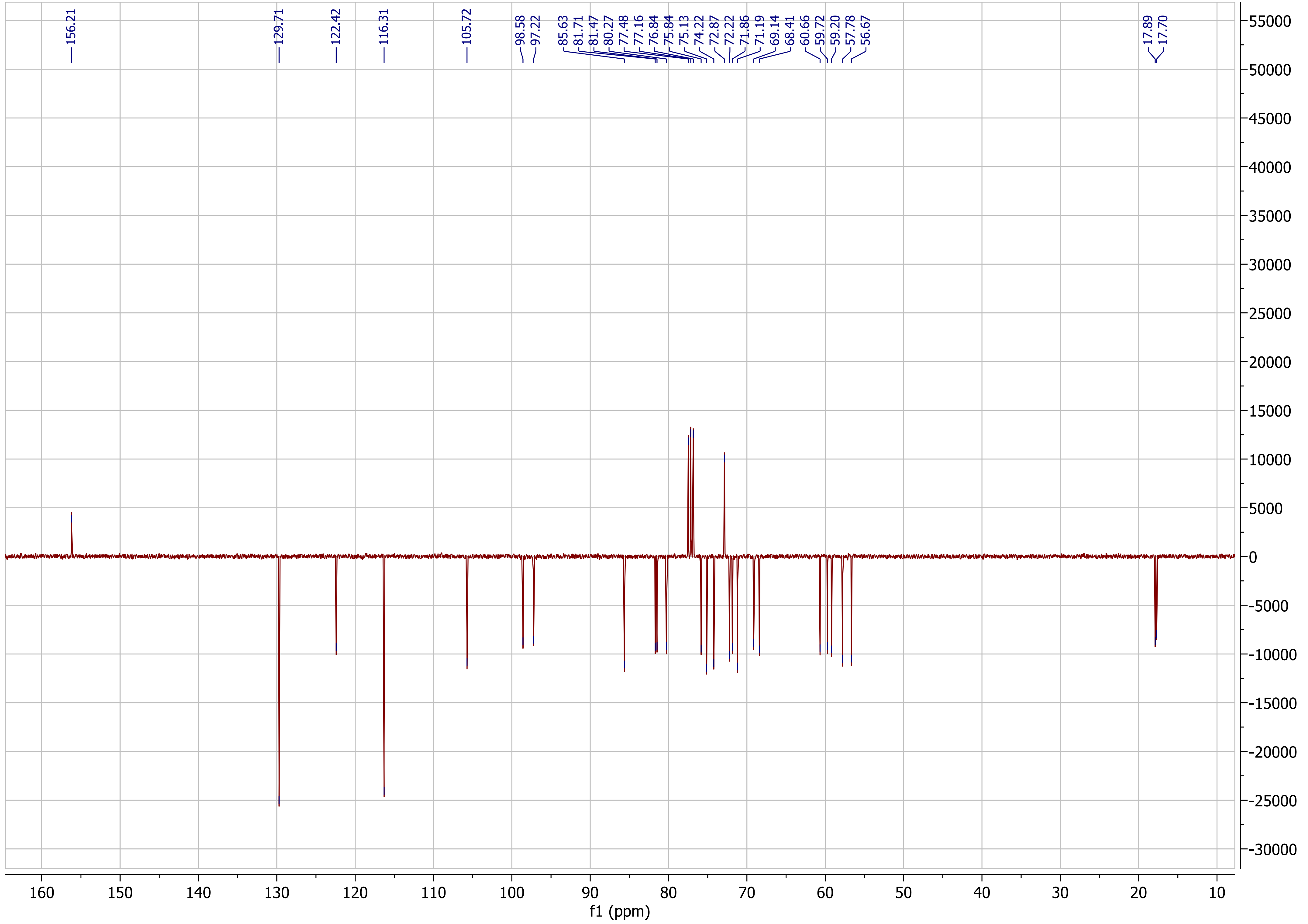

156.21
129.71
122.42
116.31
105.72
98.58
97.22
85.63
81.71
81.47
80.27
77.48
77.16
76.84
75.84
75.13
74.22
72.87
72.22
71.86
71.19
69.14
68.41
60.66
59.72
59.20
57.78
56.67
17.89
17.70
f1 (ppm)

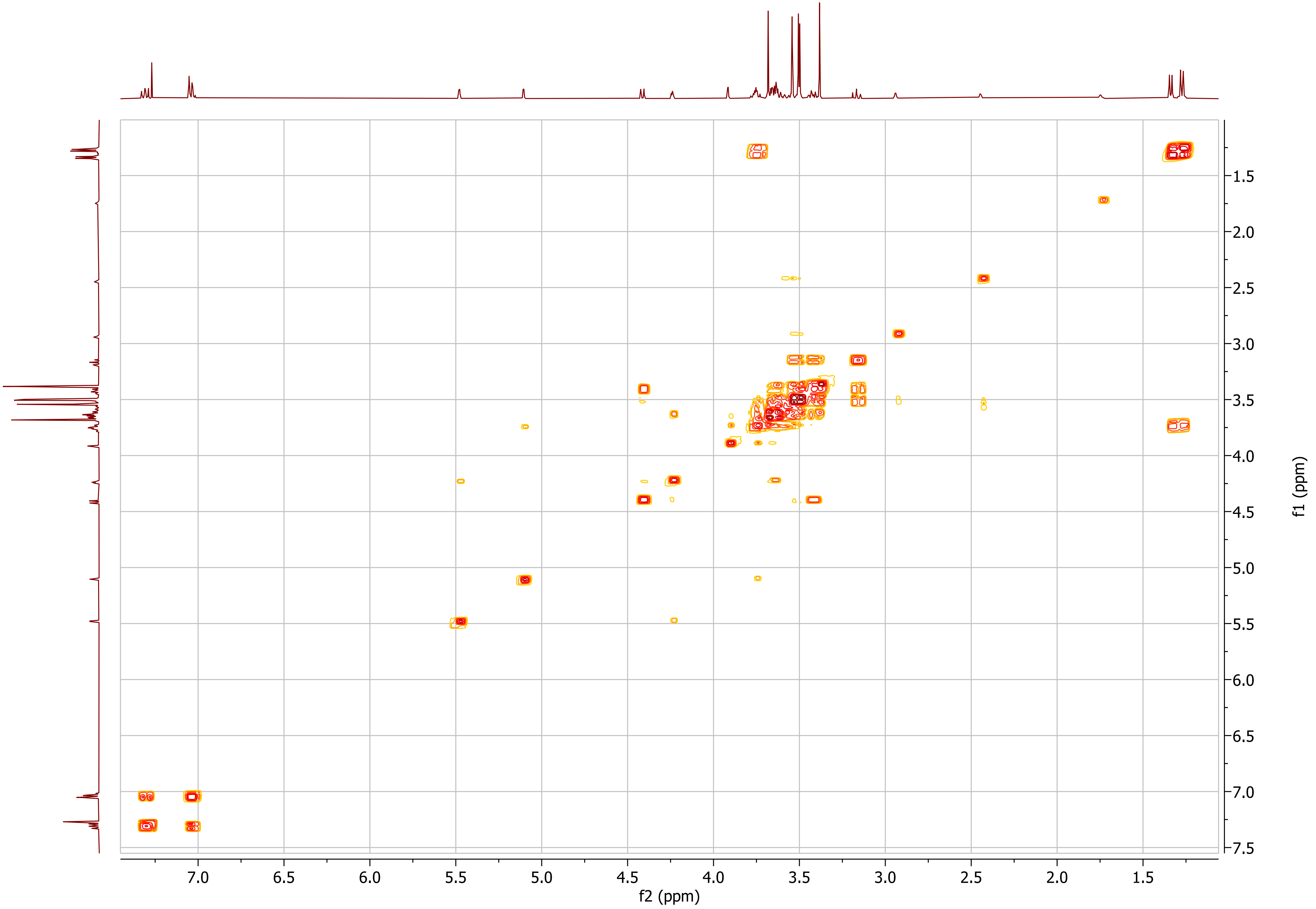

7.0
6.5
6.0
5.5
5.0
4.5
4.0
3.5
3.0
2.5
2.0
1.5
f2 (ppm)
1.5
2.0
2.5
3.0
3.5
4.0
4.5
5.0
5.5
6.0
6.5
7.0
7.5
f1 (ppm)

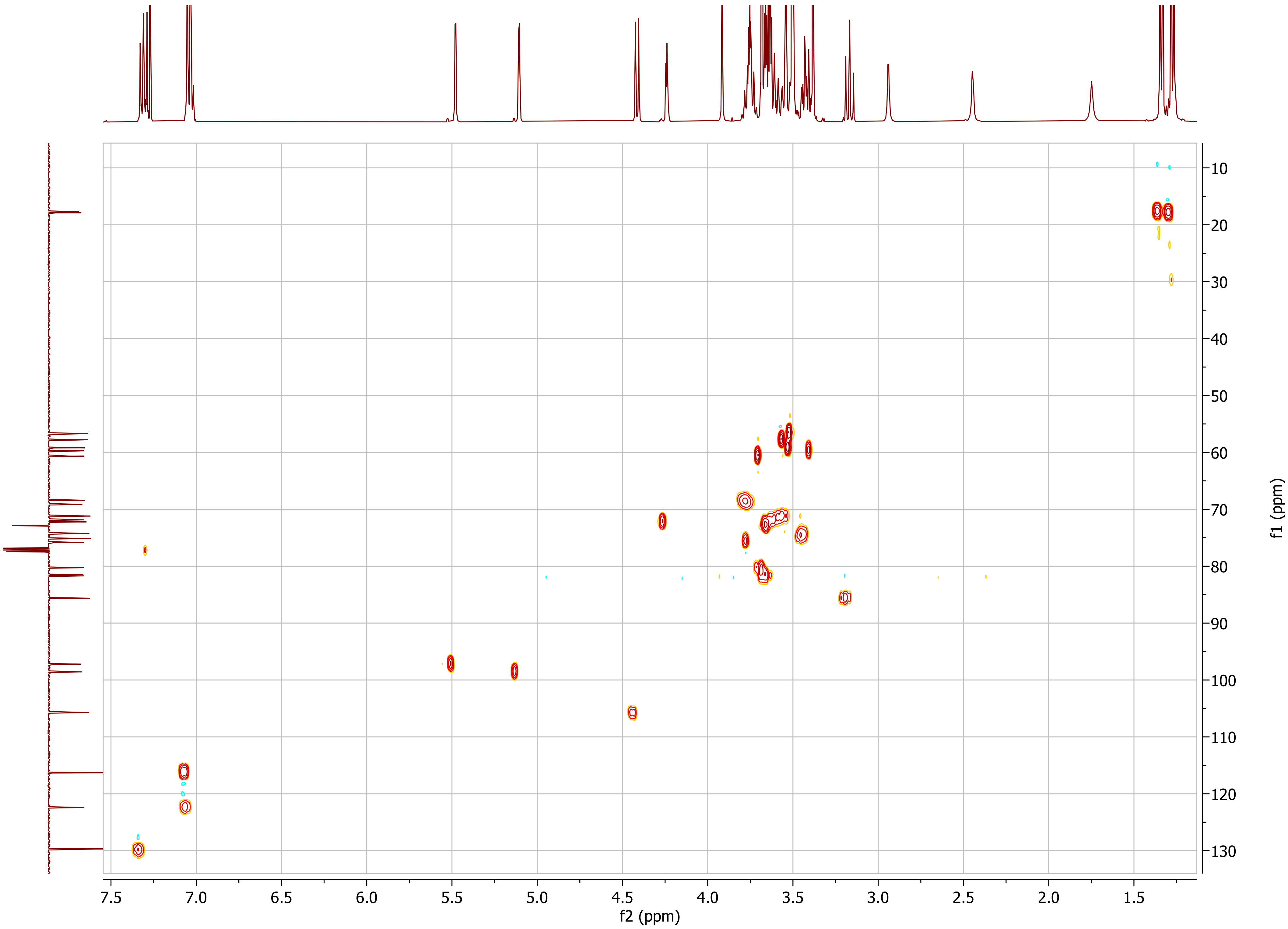

7.5
7.0
6.5
6.0
5.5
5.0
4.5
4.0
3.5
3.0
2.5
2.0
1.5
f2 (ppm)
10
20
30
40
50
60
70
80
90
100
110
120
130
f1 (ppm)

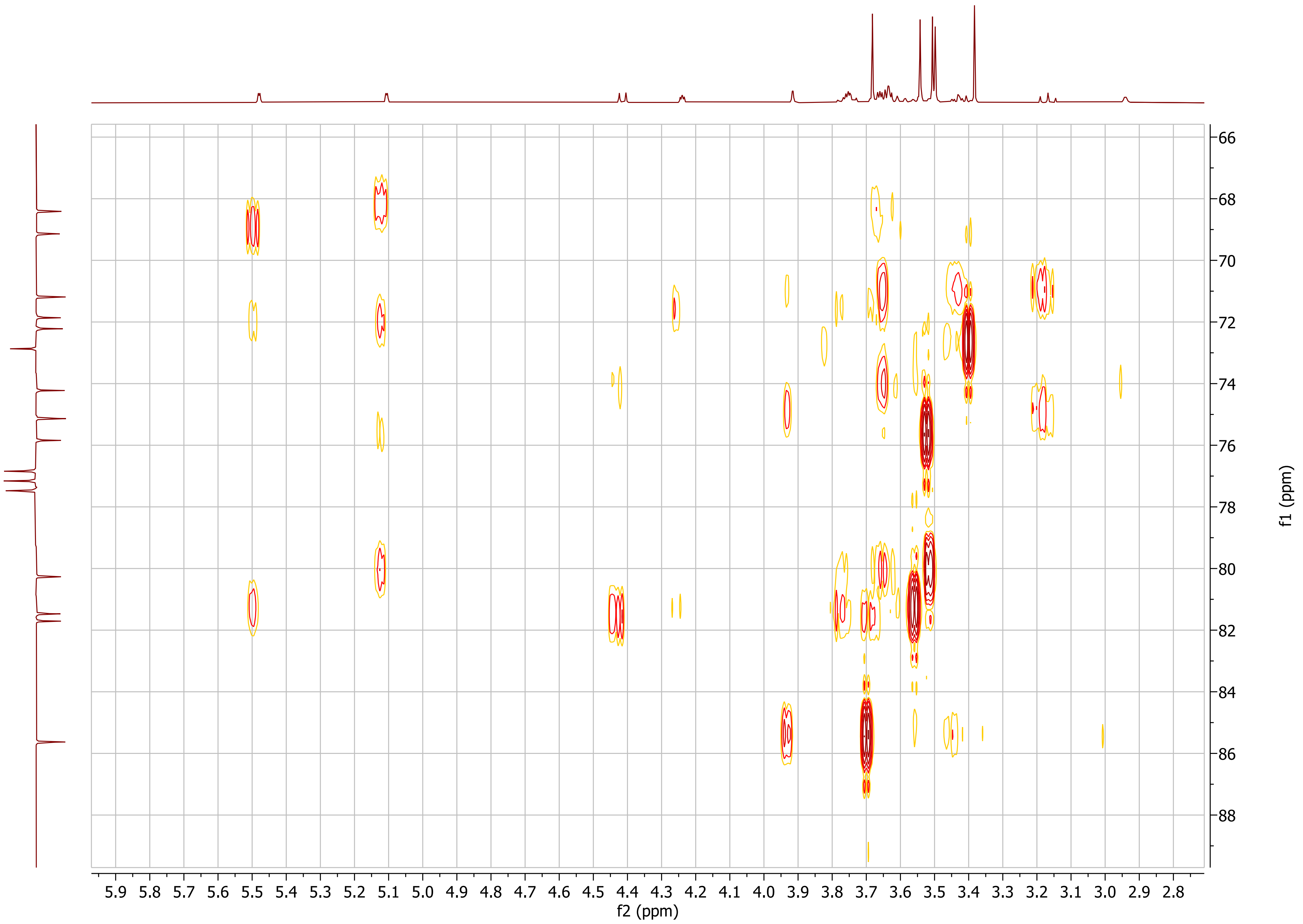

66
68
70
72
74
76
78
80
82
84
86
88
f1 (ppm)
5.9 5.8 5.7 5.6 5.5 5.4 5.3 5.2 5.1 5.0 4.9 4.8 4.7 4.6 4.5 4.4 4.3 4.2 4.1 4.0 3.9 3.8 3.7 3.6 3.5 3.4 3.3 3.2 3.1 3.0 2.9 2.8
f2 (ppm)
